# Evolution of hind limb morphology of Titanosauriformes (Dinosauria, Sauropoda) analyzed via 3D Geometric Morphometrics reveals wide-gauge posture as an exaptation for gigantism

**DOI:** 10.1101/2023.10.04.560839

**Authors:** Adrian Paramo Blazquez, Pedro Mocho, Fernando Escaso, Francisco Ortega

**Affiliations:** University of La Rioja; University of Lisbon; National University of Distance Education

## Abstract

The sauropod hind limb was the main support that allowed their gigantic body masses and a wide range of dynamic stability adaptations. It was closely related to the position of the centre of masses of their multi-ton barrel-shaped bodies, and experienced one of the most noticeable posture changes during macronarian evolution. Deeply branched macronarians achieved increasingly arched hind limbs in what is known as wide-gauge posture. However, it is not clear if this evolutionary trend is related to the evolutionary cascade toward gigantism even though some titanosaurians were the largest terrestrial vertebrates that ever existed. We tested evolutionary changes in hind limb morphology in the Macronaria phylogenetic tree by 3D geometric morphometrics. The macronarian hind limb does become progressively more arched toward deeply-branched groups, specifically Saltasauridae. However, there is morphological convergence between different macronarian subclades. Wide-gauge posture does not correlate with changes in body size deeper in the macronarian evolutionary tree, and acted as an exaptation to gigantism. Despite some titanosaurian subclades becoming some of the largest vertebrates, there is not a statistically-significant trend toward a particular body size but we identify a phyletic body size decrease in Macronaria.

## Main Text

## 1. Introduction

Sauropod dinosaurs evolved a distinct body plan that allowed them to reach some of the largest body masses of terrestrial vertebrates (e.g., Bonaparte and Coria, 1993; Carballido et al., 2017; Lacovara et al., 2014; Sander 2013; Sander et al., 2011) by one order of magnitude compared to other vertebrates, including other megaherbivore dinosaurs (e.g., Maher et al., 2020). Sauropods were dominant herbivorous dinosaurs throughout much of the Mesozoic (Sander, 2013; Sander et al., 2011). Several morphological features can be related to the evolutionary cascade that allowed the acquisition of their colossal sizes, such as the vertebral pneumaticity related to avian-like air sacs, high metabolic rates, cranial morphology, and feeding mechanisms (including the characteristic long necks among several other traits, see Sander 2013). Several features on its appendicular skeleton are related to the evolution of columnar limbs. The mechanical stability of the columnar limbs allowed them to support their multi-ton body masses (Bates et al., 2016; Lefebvre et al., 2022; Salgado et al., 1997; Sander, 2013; Ullmann et al., 2017).

Previous studies based on sauropod anatomical description, systematics, traditional morphometrics, biomechanics and GMM suggested that there was an important evolutionary trend in the appendicular skeleton toward the acquisition of a stable limb posture known as wide-gauge (Klinkhamer et al., 2019; Lefebvre et al., 2022; Salgado et al., 1997; Sander, 2013; Ullmann et al., 2017; Upchurch et al., 2004; Voegele et al., 2021; Wilson and Carrano, 1999; Wilson, 2002). This feature appears among deeply-branched Neosauropoda, in particular, among titanosauriforms, which exhibit a progressively arched limb posture (Salgado et al., 1997; Upchurch et al., 2004; Voegele et al., 2021; Wilson and Carrano, 1999; Wilson, 2002). The wide stance may enable enhanced lateral stability during locomotion allowing them to exploit more efficiently inland environments (Mannion and Upchurch, 2010; Ullmann et al., 2017). However, the widening of the body in Titanosauriformes and the acquisition of the wide-gauge stance is still poorly understood. Although the arched limbs and wider postures appeared in the largest sauropods among Titanosauriformes (Bates et al., 2016; Henderson, 2006; Ullmann et al., 2017; Wilson and Carrano, 1999) these features may at least be acquired independently among both small and large titanosauriforms of different subclades (Henderson 2006; Bates et al. 2016). Limb posture may have been related to achieving greater ranges of ecological niches though biomechanical stability rather than allowing increasing their body size itself (e.g., Bates et al., 2016; Henderson, 2006; Ullmann et al., 2017). The study of the hind limb through 2D and 3D geometric morphometrics (GMM) allows us to analyze complex morphological changes across the titanosauriformes phylogeny (e.g., Lefebvre et al., 2022, Ullmann et al., 2017). Here we will use 3D-digitized and reconstructed titanosauriform hind limbs (Table 1, see also Fig. 1) to test whether there is a relationship between wider hind limb posture (as the hind limbs are the main weight support of the sauropod body, see Bates et al. 2016) and their body size, as proxied by the hind limb size. We chose the centroid size of the hind limb because the size of the femur and tibia correlates well with sauropod body mass (Campione & Evans, 2020, Mazzetta et al., 2004), but alternative tests using femoral length or body mass estimations are provided in Appendix S2-4. We will also test for potential morphological convergence between different titanosauriform subclades regardless of their body size. The analysis presented here can be seen as an expansion of the Lefebvre et al. (2022) study on sauropod limb evolution as we analyze a sample comprised mostly of Late Cretaceous lithostrotian sauropods (their study included a broad diversity of sauropodomorph taxa including several titanosaurs).

**Figure 1.**
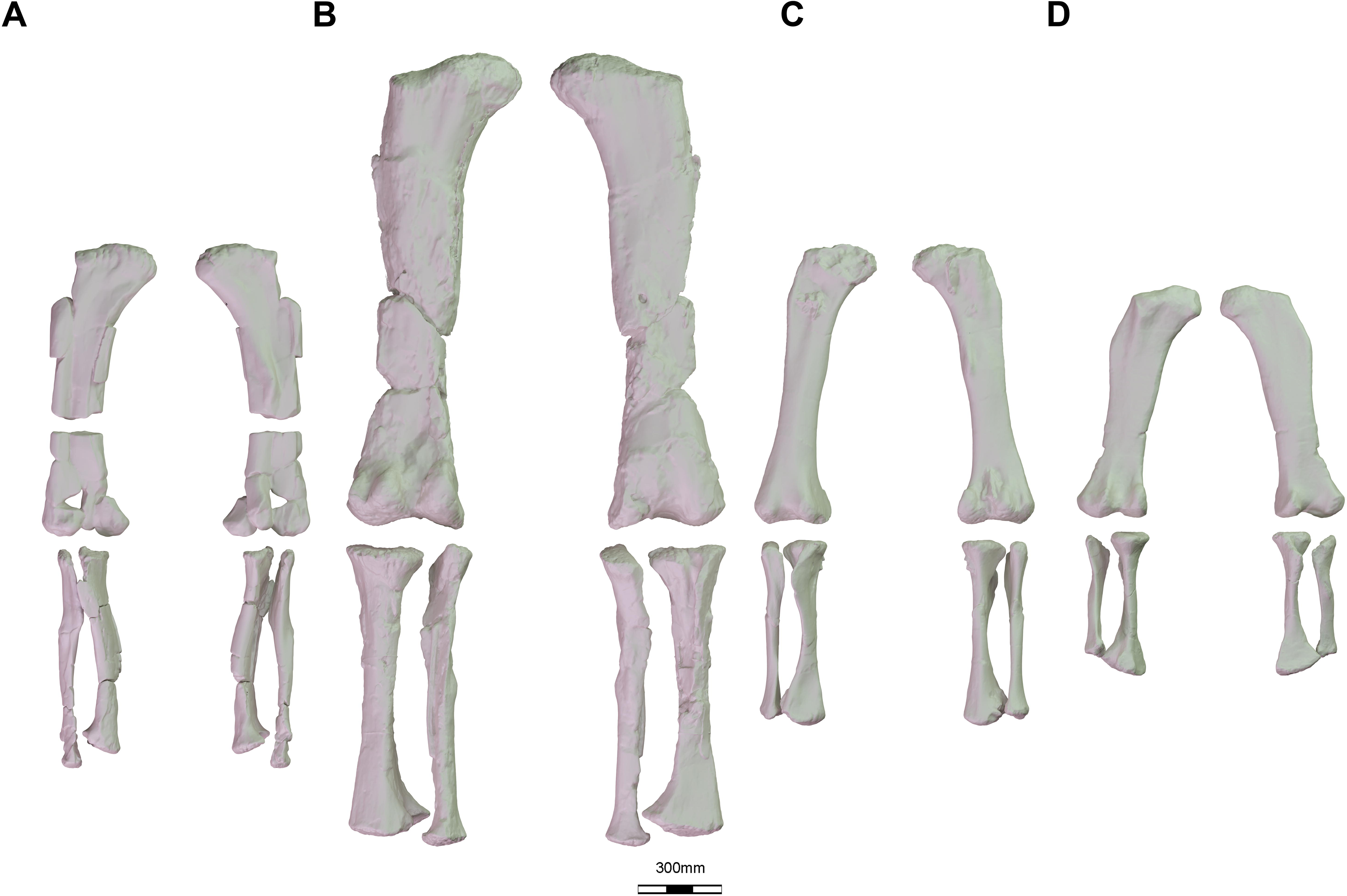
Sample of several 3D reconstruction of macronarian hind limbs used in this study. (a) *Oceanotitan dantasi* in anterior and posterior view; (b) *Ligabuesarusu leanzai* in posterior and anterior view; (c) *Lohuecotitan pandafilandi* in anterior and posterior view; (d) *Saltasaurus loricatus* in posterior and anterior view.

**Table 1.**
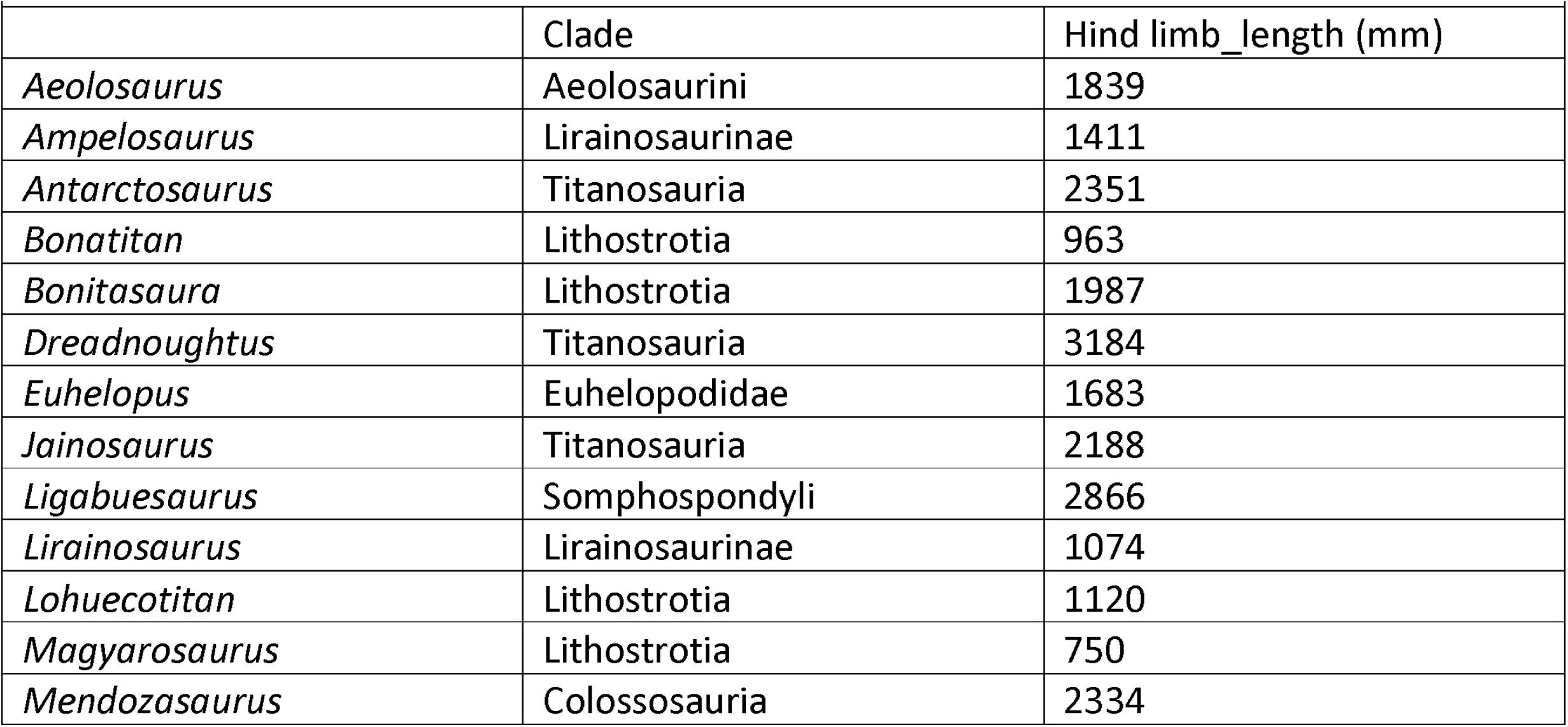
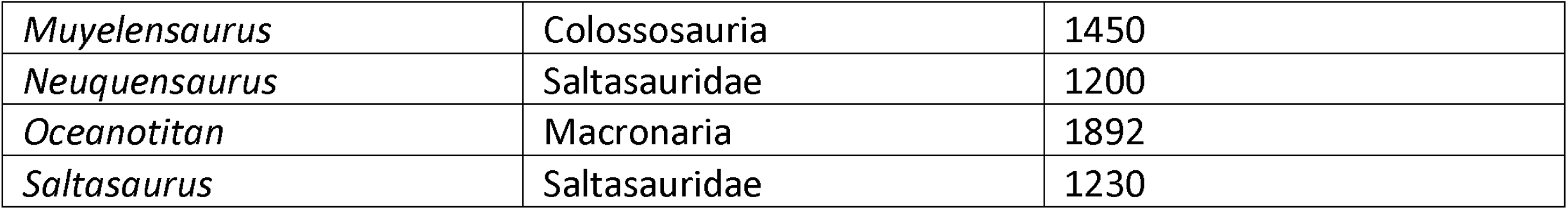
Specimen sample used in this study. Proximodistal hind limb length measured in mm.

## 2. Results

### 2.1. Titanosauriformes Morphospace Occupation

A Principal Component Analysis (PCA) was performed to generate an occupation morphospace, obtaining a total of 16 shape Principal Components (after Anderson’s χ test; PC from now on). The first six PCs accounted for 78.3% of the cumulative morphological variation (Table 2). The non-parametric Kruskal-Wallis test shows that no single shape variable reports significant differences among sauropod subclades (Table 3 and 4). Here we only comment on the results of the first three shape PCs (>50% of the cumulative variance) due to space limitations, but a full description and visualization of the complete PCA and phylomorphospace projections can be found in Appendix S3.

**Table 2.**
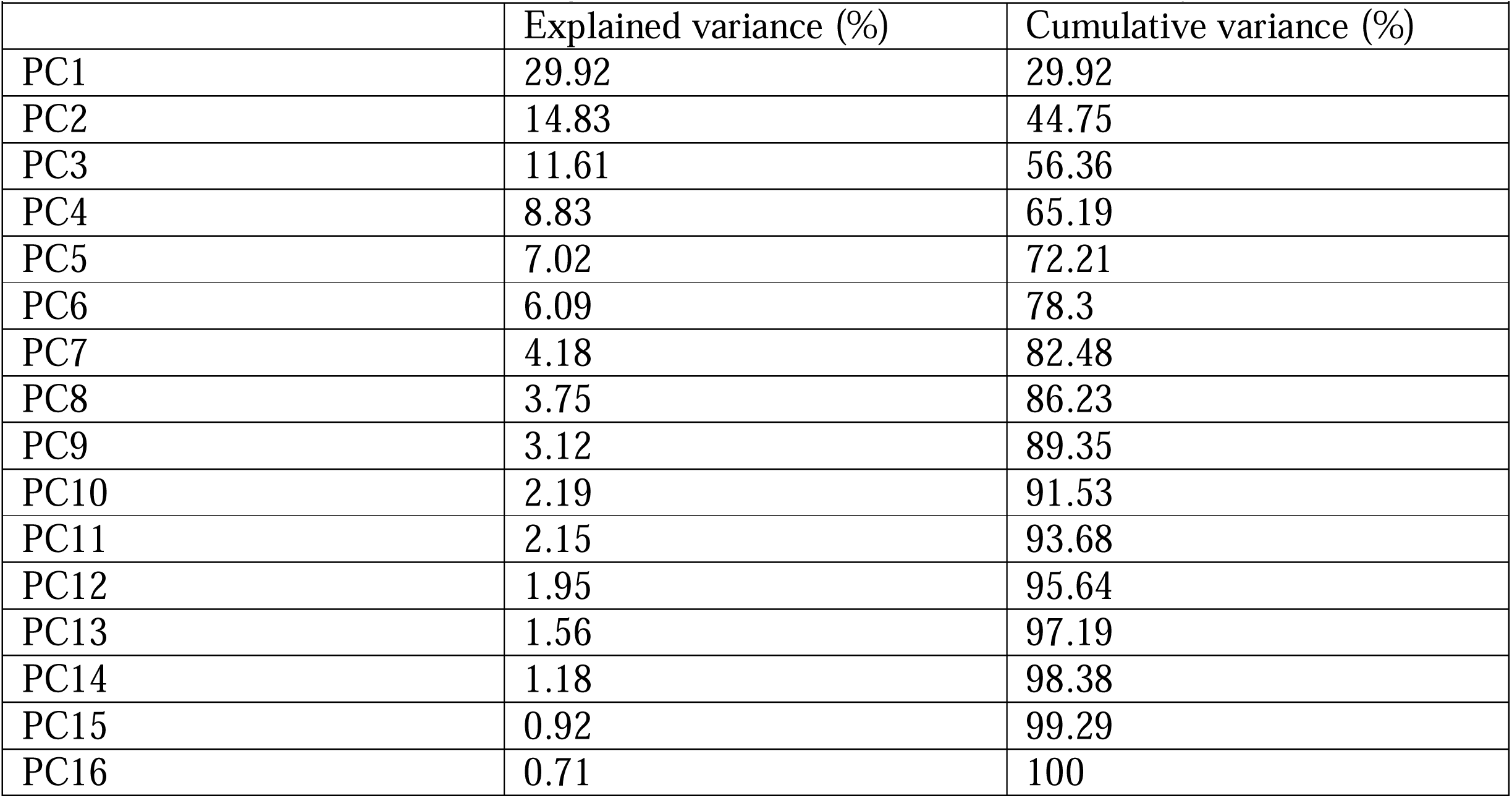
PCA results over GPA aligned coordinates. Variance explained by each shape PC.

**Table 3.** Kruskal-Wallis test on shape PCA variables between the most inclusive subclades analysed.

|  |  |  |  |
| --- | --- | --- | --- |
|  | Chi-sq | p-value | p-adjusted |
| PC1 | 11.954 | 0.153 | 1 |
| PC2 | 5.592 | 0.693 | 1 |
| PC3 | 7.886 | 0.445 | 1 |
| PC4 | 9.647 | 0.291 | 1 |
| PC5 | 5.17 | 0.739 | 1 |
| PC6 | 7.618 | 0.472 | 1 |
| PC7 | 8.915 | 0.35 | 1 |
| PC8 | 8.941 | 0.347 | 1 |
| PC9 | 10.206 | 0.251 | 1 |
| PC10 | 10.422 | 0.237 | 1 |
| PC11 | 4.768 | 0.782 | 1 |
| PC12 | 5.66 | 0.685 | 1 |
| PC13 | 8.886 | 0.352 | 1 |
| PC14 | 9.856 | 0.275 | 1 |
| PC15 | 10.248 | 0.248 | 1 |
| PC16 | 4.578 | 0.802 | 1 |

PC1 (summarizing 29.92% of the total variance; Fig. 2a-b) is associated with characters describing the orientation and compression of the femoral shaft, the length of the distal femoral condyles, the orientation of the fourth trochanter, and the length and width of the zeugopod bones (Fig. 2c). Non-titanosaurian macronarians occupy negative values, whereas non-lithostrotian titanosaurs are distributed between weakly negative (e.g., *Bonatitan*) and positive PC1 values (Fig. 4b). Lithostrotian titanosaurs occupy a wide range of values, with *Aeolosaurus*, the specimens of Colossosauria and *Lirainosaurus* clustering at negative PC1 values (however, *Ampelosaurus* occupies negative PC1 values approaching zero; Fig. 2b). Most of the non-saltasaurid, non-colossosaurian, non-lirainosaurine and non-aeolosaurine lithostrotians are broadly distributed in the morphospace between negative and positive values, with *Bonitasaura* occupying the farthest negative PC1 values and overlapping with Colossosauria and Lirainosaurinae (Fig. 2b). The highest positive PC1 values are occupied by saltasaurids (i.e., *Saltasaurus* and *Neuquensaurus*) (Fig. 2b). There is a trend from non-titanosaurian macronarians at negative values to the titanosaurian node at weakly negative values (Fig. 2b). In this PC1, the Titanosauria node splits near a zero score, with *Lohuecotitan* occupying weakly positive PC1 values near a zero score [in recent phylogenetic analyses, *Lohuecotitan* has been recovered as a member of Lithostrotia (Díez Díaz et al., 2020; Navarro et al., 2022; Mocho et al. 2024a)], but other deeply nested titanosaurs occupy positive scores in PC1 (Fig. 2b). The trend toward positive values follows with several other deeply nested lithostrotians. However, both Colossosauria and the two analysed members of Lirainosaurinae fall into negative PC1 values (Lirainosaurinae was not recovered as a monophyletic group in our current topology; Fig. 2b). There are no significant differences between the different sauropod subclades in this PC (Appendix S4).

**Figure 2.**
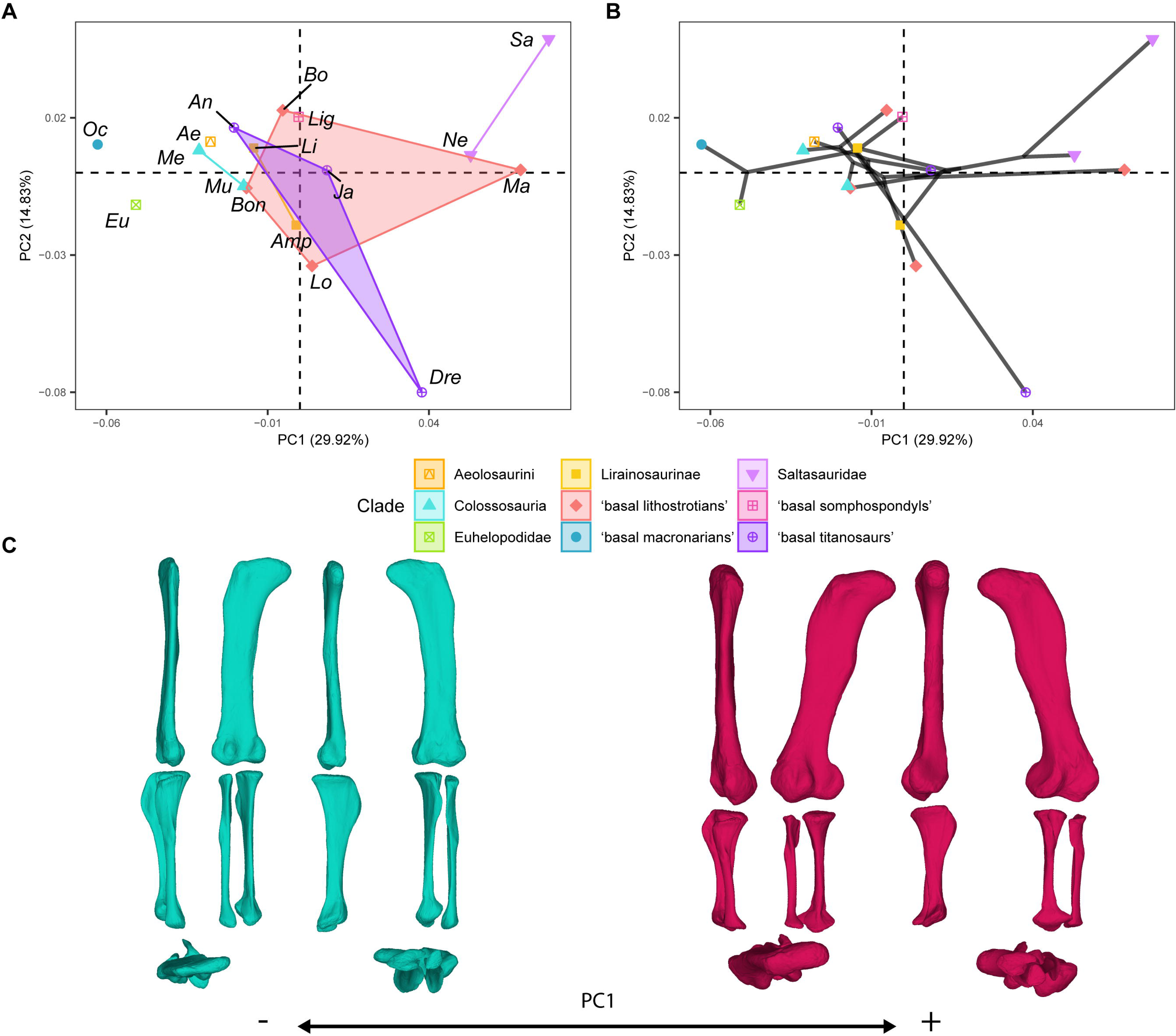
PCA results on the GPA aligned landmark and semilandmark curves of the hind limb. (a) PC1-PC2 biplot. (b) PC1-PC2 phylomorphospace with projected phylogenetic tree. (c) Representation of the shape change along PC1, blue are negative scores, red are positive scores. Percentage of variance of each PC in brackets under corresponding axis. *Ae* – *Aeolosaurus*, *Amp* – *Ampelosaurus*, *An* – *Antarctosaurus*, *Bo* – *Bonatitan*, *Bon* – *Bonitasaura*, *Dre* – *Dreadnoughuts*, *Eu* – *Euhelopus*, *Ja* – *Jainosaurus*, *Li* – *Lirainosaurus*, *Lig* – *Ligabuesaurus*, *Lo* – *Lohuecotitan*, *Ma* – *Magyarosaurus*, *Me* – *Mendozasaurus*, *Mu* – *Muyelensaurus*, *Ne* – *Neuquensaurus*, *Sa* – *Saltasaurus*.

PC2 (summarizing 14.83% of the variance; Fig. 3a-b) is associated with characters describing the orientation of the femoral shaft; the length and orientation of the femur proximal end; the length, width and orientation of the distal femoral condyles; the orientation of the tibial shaft; the length and orientation of the cnemial crest; the length, width and orientation of the tibial distal end; the orientation of the tibial ascending process; the orientation of the fibular shaft and fibular anterior crest (Fig. 3c). Both the early branching macronarian *Oceanotitan* - a possible member of Somphospondyli (Mocho et al., 2019) - and the euhelopodid somphospondyli *Euhelopus* occupy positive and negative values near a zero score, whereas the early-diverging somphospondylan *Ligabuesaurus* is distributed at slightly more positive PC2 values than *Oceanotitan* (Fig. 3b). However, there is no clear pattern among the titanosaurian subclade morphospace (Fig. 3a) or the phylomorphospace (Fig. 3b) among progressively deeply branching titanosauriforms. The saltasaurids and *Aeolosaurus* occupy positive PC2 scores (Fig. 3b) showing some overlap with Colossosauria at weakly positive values of PC2 (Fig. 3b). Lirainosaurines and colossosaurians are distributed at weakly negative to positive values in PC2 (Fig. 3b). However, non-saltasaurid, non-aeolosaurine, non-colossosaurian, and non-lithostrotian titanosaurs are broadly distributed across the morphospace; most of them between negative values (i.e. *Dreadnoughtus*) and positive PC2 values (i.e *Jainosaurus*, *Antarctosaurus*) (Fig. 3a-b). The hind limb occupation of the titanosaur *Dreadnoughtus* at strongly negative PC2 values is noteworthy (Fig. 3a-b). The Mann-Whitney’s U test found no significant differences in the PC2 in any of the pairwise comparisons (Appendix S4).

**Figure 3.**
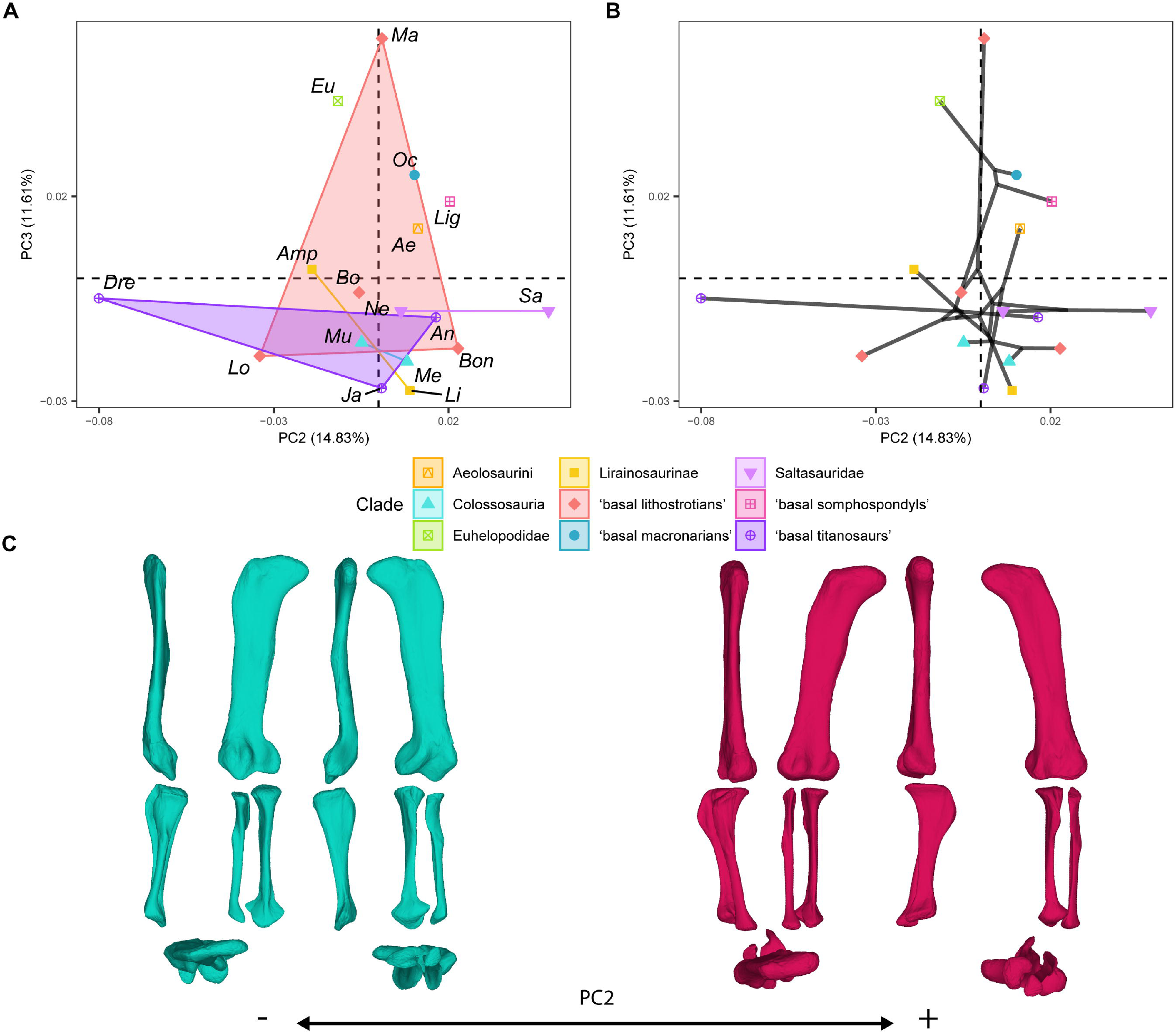
PCA results for the GPA aligned landmark and semilandmark curves of the hind limbs. (a) PC2-PC3 biplot. (b) PC2-PC3 phylomorphospace with projected phylogenetic tree. (c) Representation of the shape change along PC2, blue are negative scores, red are positive scores. Percentage of variance of each PC in brackets under corresponding axis. *Ae* – *Aeolosaurus*, *Amp* – *Ampelosaurus*, *An* – *Antarctosaurus*, *Bo* – *Bonatitan*, *Bon* – *Bonitasaura*, *Dre* – *Dreadnoughuts*, *Eu* – *Euhelopus*, *Ja* – *Jainosaurus*, *Li* – *Lirainosaurus*, *Lig* – *Ligabuesaurus*, *Lo* – *Lohuecotitan*, *Ma* – *Magyarosaurus*, *Me* – *Mendozasaurus*, *Mu* – *Muyelensaurus*, *Ne* – *Neuquensaurus*, *Sa* – *Saltasaurus*.

**Figure 4.**
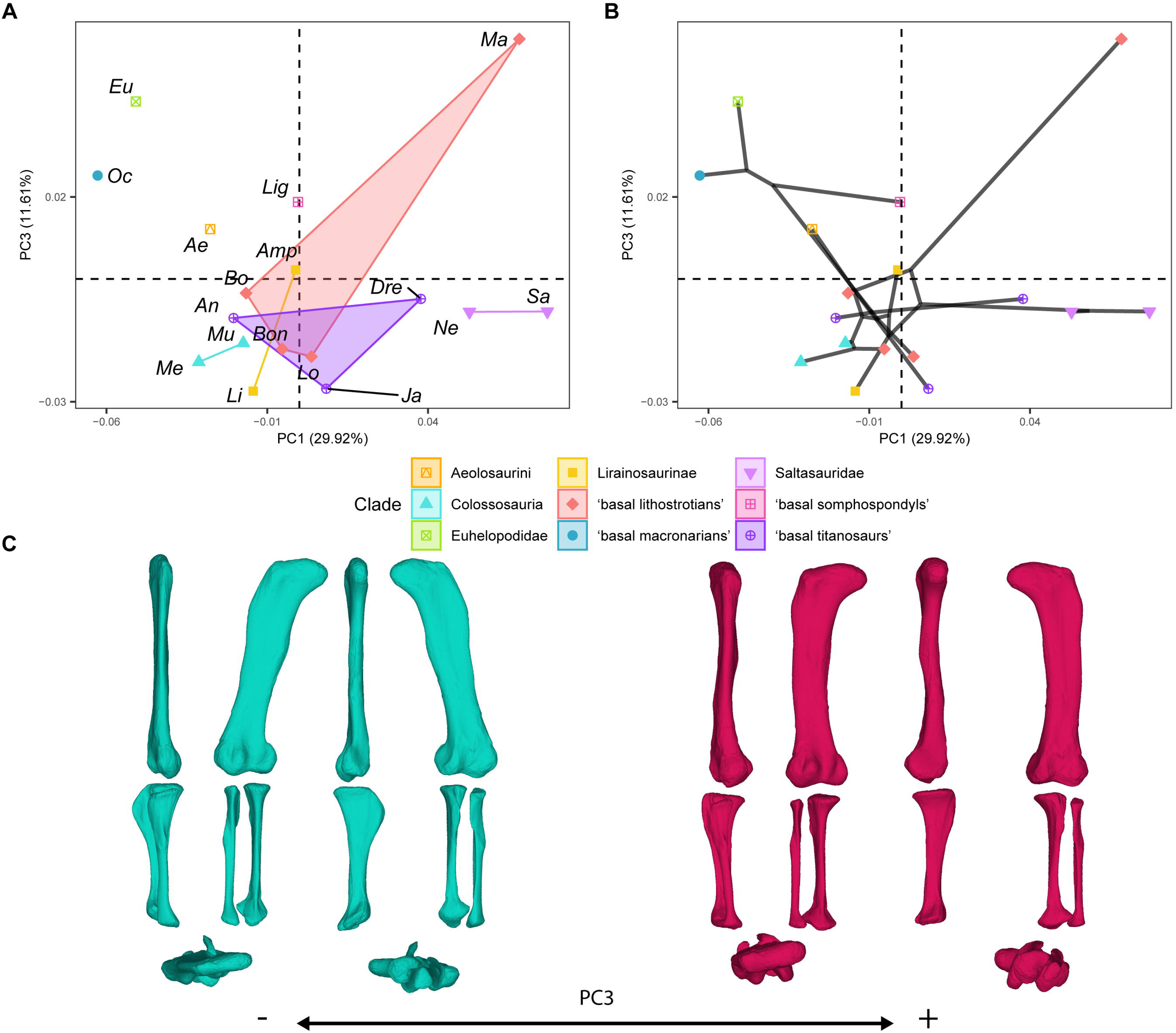
PCA results for the GPA aligned landmark and semilandmark curves of the hind limbs. (a) PC1-PC3 biplot. (b) PC1-PC3 phylomorphospace with projected phylogenetic tree. (c) Representation of the shape change along PC3, blue are negative scores, red are positive scores. Percentage of variance of each PC in brackets under corresponding axis. *Ae* – *Aeolosaurus*, *Amp* – *Ampelosaurus*, *An* – *Antarctosaurus*, *Bo* – *Bonatitan*, *Bon* – *Bonitasaura*, *Dre* – *Dreadnoughuts*, *Eu* – *Euhelopus*, *Ja* – *Jainosaurus*, *Li* – *Lirainosaurus*, *Lig* – *Ligabuesaurus*, *Lo* – *Lohuecotitan*, *Ma* – *Magyarosaurus*, *Me* – *Mendozasaurus*, *Mu* – *Muyelensaurus*, *Ne* – *Neuquensaurus*, *Sa* – *Saltasaurus*.

Finally, PC3 (summarizing 11.61% of the variance; Fig. 3a-b, 4a-b) is related to characters describing the orientation of the femoral shaft and head; the length of the femoral lateral bulge; the location and orientation of the fourth trochanter; the orientation of the distal femoral condyles; the length and orientation of the tibial cnemial crest; the length of the tibial distal end; the length of the tibial ascending process; and the length and morphology of the fibula (Fig. 4c). *Oceanotitan* and *Euhelopus* were plotted toward progressively more positive values (Fig. 4a) and the node was estimated at slightly more negative values of PC3 than *Euhelopus* (Fig. 4b). *Ligabuesaurus* occupies positive values of PC3 closer to zero than early branching titanosauriforms (Fig. 4a). In the phylomorphospace, there is a trend from positive values to negative values of PC3 (Fig. 4b) from more basally branching titanosauriforms to more deeply nested ones. Non-lithostrotian titanosaurs occupy negative PC3 values but the non-saltasaurid, non-lirainosaurine and non-colossosaurian lithostrotians trends towards positive PC3 values (Fig. 4a-b), whereas Lirainosaurinae, Colossosauria and Saltasauridae occupy negative values of PC3 (Fig. 4a-b). Colossosauria, Saltasauridae and *Lirainosaurus*, the most deeply-branched representatives of Lirainosaurinae according to our phylogenetic hypothesis, plotted toward increasingly more negative PC3values (Fig. 4a-b). *Magyarosaurus* is the only lithostrotian occupying high positive values, clearly separate from all other sauropods (Fig. 4a-b). The pair-wise Mann-Whitney’s U test found no significant differences between sauropod subclades in this PC (Appendix S4).

### 2.2. Size distribution

There is a trend in hind limb centroid size distribution from large, non-titanosaurian macronarians in the Early Cretaceous (e.g., *Euhelopus*) to small, deeply nested lithostrotian titanosaurs in the Late Cretaceous (Fig. 5). This trend coincides with the morphospace occupation recovered by PC1 (early-branching and larger non-titanosaurian macronarians among negative PC1 values and progressively more deeply nested and smaller titanosaurs toward positive PC1 values (Fig. 5). The smallest lithostrotians are concentrated at deeply branching nodes, including the Ibero-Armorican lirainosaurines, *Bonatitan reigi*, *Magyarosaurus* spp. and members of Saltasaurinae (Fig. 5).

**Figure 5.**
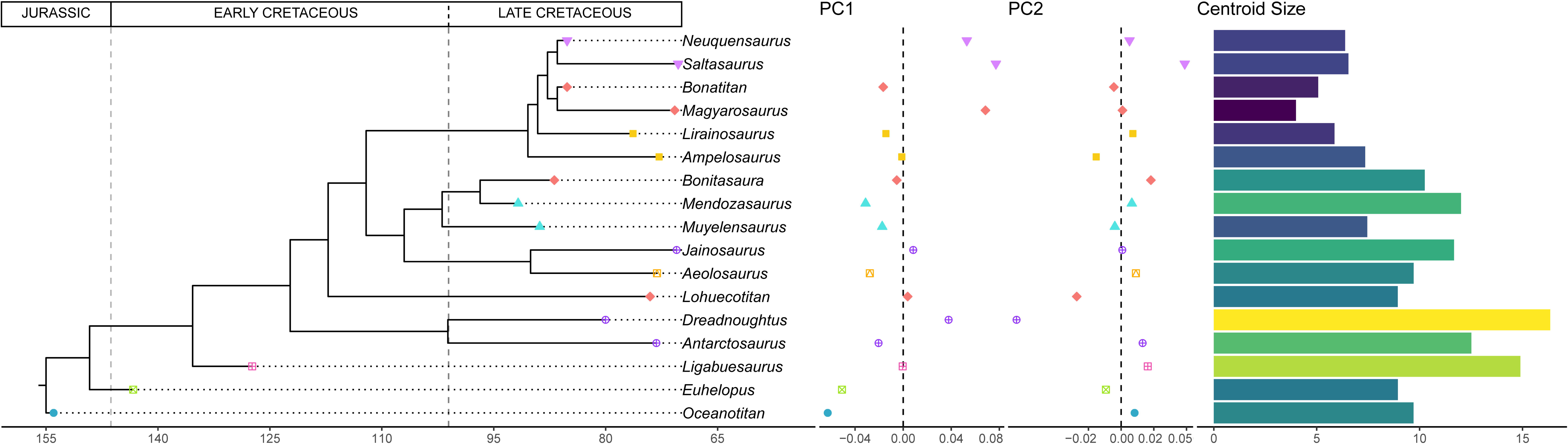
Time calibrated supertree with PC1-PC2 results and hind limb centroid size for each sauropod.

However, the RMA models found no significant correlation between the shape variables and the log-transformed centroid size (Table 6, see Fig. 6 and Appendix S3). The PC1 model (r^2^ = 0.105, p-value= 0.204; Fig. 6a) found negative allometry but no significant correlation and the percentage of variance explained by hind limb size differences was small. Almost all of the RMA found a negative relationship, except for several sub-sampled RMA models like the PC2 against log-transformed centroid size for the sample of lithostrotian titanosaurs only (Fig. 6b). None of the RMA models for the sub-samples found a significant correlation (see Appendix S3).

**Figure 6.**
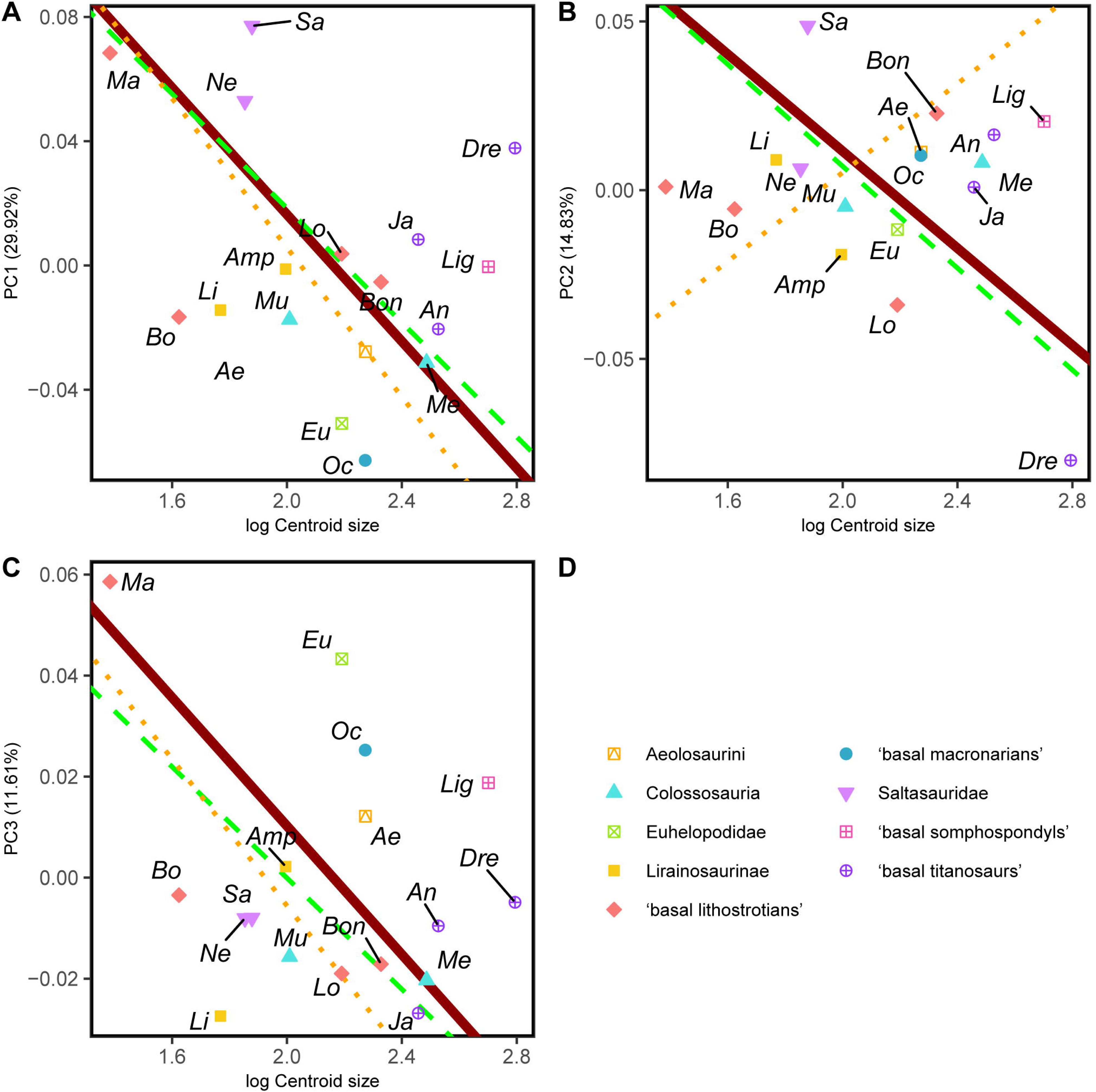
RMA results of the first three shape PCs against the logarithm of the hind limb centroid size. (a) PC1 against log-Centroid size, all taxa RMA in dark red: intercept = 0.221, slope = −0.102, *r^2^* = 0.105, *p* = 0.204; Titanosauria only partial RMA in dashed green: intercept = 0.203, slope = −0.092, *r^2^* = 0.118, *p* = 0.229; Lithostrotia only partial RMA in dotted orange: intercept = 0.246, slope = −0.120, *r^2^* = 0.319, *p* = 0.07; (b) PC2 against log-Centroid size, all taxa RMA in dark red: intercept = 0.155, slope = −0.072, *r^2^* = 0.054, *p* = 0.371; Titanosauria only partial RMA in dashed green: intercept = 0.158, slope = −0.075, *r^2^* = 0.117, *p* = 0.232; Lithostrotia only partial RMA in dotted orange: intercept = −0.127, slope = 0.066, *r^2^* = 0, *p* = 0.952; (c) PC3 against log-Centroid size (Csize), all taxa RMA in dark red: intercept = 0.137, slope = −0.064, *r^2^* = 0.055, *p* = 0.363; Titanosauria only partial RMA in dashed green: intercept = 0.110, slope = −0.055, *r^2^* = 0.236, *p* = 0.078; Lithostrotia only partial RMA in dotted orange: intercept = 0.140, slope = −0.073, *r^2^* = 0.313, *p* = 0.074. *Ae* – *Aeolosaurus*, *Amp* – *Ampelosaurus*, *An* – *Antarctosaurus*, *Bo* – *Bonatitan*, *Bon* – *Bonitasaura*, *Dre* – *Dreadnoughuts*, *Eu* – *Euhelopus*, *Ja* – *Jainosaurus*, *Li* – *Lirainosaurus*, *Lig* – *Ligabuesaurus*, *Lo* – *Lohuecotitan*, *Ma* – *Magyarosaurus*, *Me* – *Mendozasaurus*, *Mu* – *Muyelensaurus*, *Ne* – *Neuquensaurus*, *Sa* – *Saltasaurus*.

### 2.3. Phylogenetic trends

Pagel’s lambda (λ) estimation shows a significative phylogenetic signal in log-transformed hind limb centroid size (λ = 0.982), and PC1 (λ = 0.715), PC3 (λ = 0.760), PC5 (λ = 0.778) and PC6 (λ = 0.697) therefore exhibiting a trend in the evolution of Titanosauriformes (Table 7). We estimated ancestral characters (ACEs) using log-transformed hind limb centroid size and those shape variables (the PCs) that exhibit a significant signal during the evolution of Titanosauriformes and tested for a directionality or trend (Fig. 7). The resulting tests recover significant trends toward a decrease in hind limb size across all titanosauriform subclades, with positive PC1 values including somphospondyli titanosauriformes, and negative PC3 values across all titanosauriform subclades (electronic Appendix S4, table S4.8, S4.9, S4.10).

**Figure 7.**
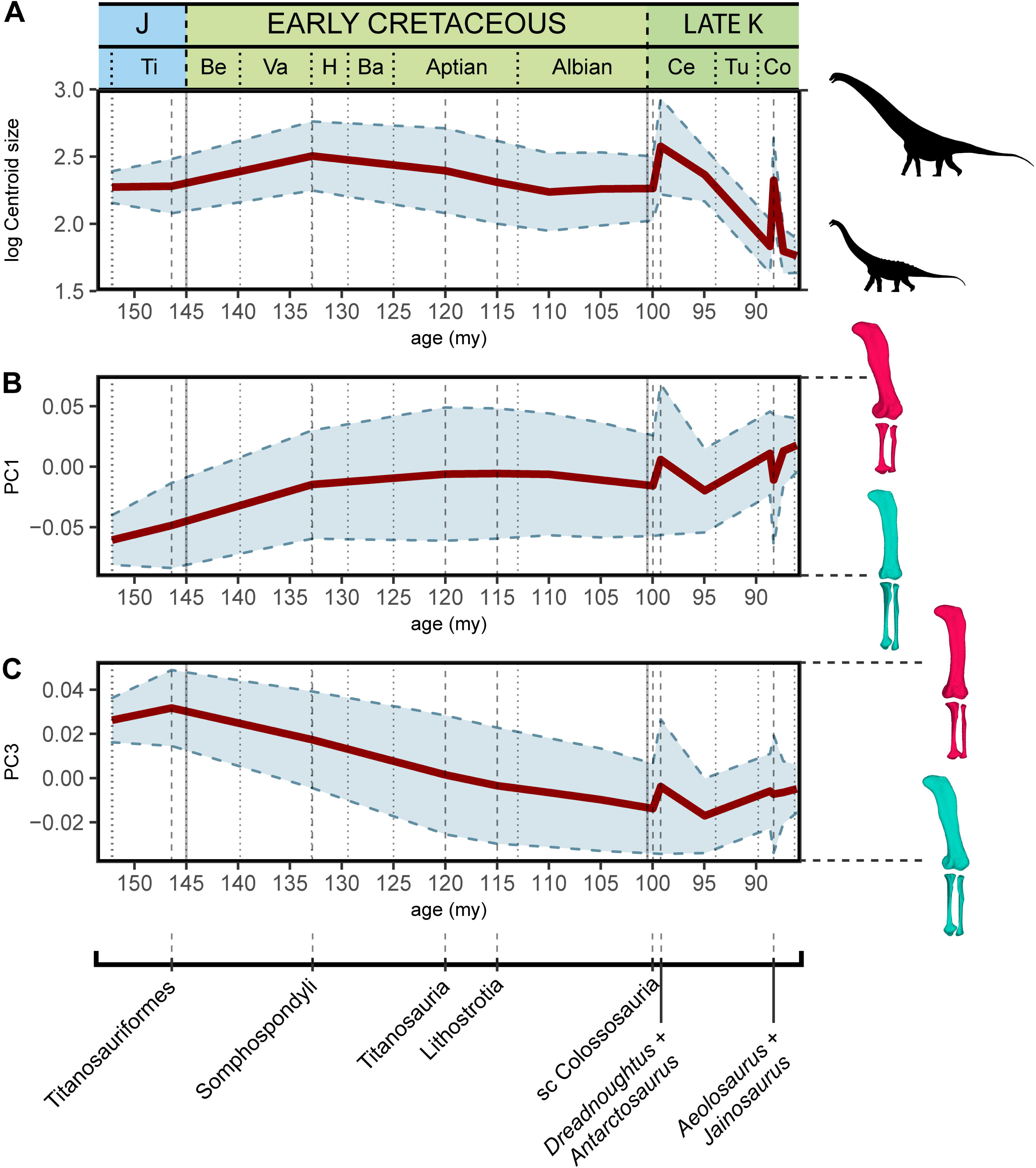
Evolution of log-transformed hind limb centroid size, shape PC1 and PC3 according to our sample and time-calibrated supertree topology. Ba – Barremian, Be – Berriasian, Ce – Cenomanian, Co – Conacian, H – Hauterivian, J – Jurassic, Late-K – Late Cretaceous, Ti – Tithonian, Tu – Turonian, Va – Valanginian. my – million years from present, sc – subclade Colossosauria + *Bonitasaura*.

## 3. Discussion

### 3.1. Hind limb morphological convergence in Titanosauriformes

Analysis of the shape variables extracted by PCA on the Procrustes coordinates of the sauropod taxa reveals a large overlap between the different titanosauriform subclades, in particular within Titanosauria (e.g., Fig. 2-4), across all the resulting shape PCs. Both the non-parametric tests on the hind limb shape variables and the size, and the phylogenetic ANOVA accounting for the time-calibrated supertree topology, suggest the lack of sufficient and significant morphological differences between the different titanosauriform subclades studied in this analysis. Based on the lack of significant phylogenetic differences and the presence of morphological similarities, the evolutionary pattern observed for the titanosaurian hind limb may be explained by convergent evolution, consistent with previous analyses (Lefebvre et al., 2022; Páramo et al., 2020; Ullmann et al., 2017).

Considering the analysed sample, the acquisition of wide-gauge locomotion would be the main source of hind limb morphological variability in titanosaurian sauropods (Fig. 2, 5) and possesses a significative phylogenetic signal. The trend toward a more arched limb posture persists in the more deeply nested titanosaurs. *Oceanotitan dantasi*, a possible representative of Somphospondyli, exhibits a columnar hind limb characterized by no deflection of the femoral head with regard to the tibial condyle (though some titanosauriforms exhibits this feature, see Royo-Torres, 2009; recent phylogenetic approaches suggest that *O. dantasi* might represent non-titanosauriform macronarian, see Mocho et al. 2024b), and a straight and long, lateromedially-narrow zeugopod with few anterior rotations of the fibula (Fig. 2). In contrast, the more deeply branching titanosauriforms exhibit the typical titanosaurian hind limb configuration with a more arched posture, lateromedially more robust femora, and increased medial or proximo-medial deflection of the femoral head. Distally, the hind limb exhibits a slight rotation of the femoral distal ends, and increasingly lateromedially robust zeugopods. The robust zeugopod elements are the only resemblance to *O. dantasi* as our outgroup (e.g., Fig. 2). The highest positive PC1 (Fig. 6) values correlate with several hyper-robust taxa of different titanosaur subclades that exhibit lateromedially and anteroposteriorly wide stylopods and zeugopods. In these taxa, the zeugopod bones are extremely shorter proximodistally, extremely arched with a predominance of tibiae characterized by short cnemial crests but somewhat rotated, interlocking with anteriorly deflected and robust fibulae as in the saltasaurine *Saltasaurus loricatus*, the lithostrotian *Magyarosaurus* spp. and the possible non-lithostrotian titanosaur *Dreadnoughtus schrani* (based on the super-tree: Fig. 2,5). The acquisition of this particular morphology is correlated with the development of gigantism within Titanosauria (Carrano, 2005; Lefebvre et al., 2022; Ullmann et al., 2017), at least when analysing early branching members of Lithostrotia. However, specimens occupying higher PC1 values exhibit the most hyper-robust and arched hind limbs including representatives of the smallest lithostrotians and the largest non-lithostrotian titanosaur studied (Fig. 5). The analyses of evolutionary trends presented here reveal that the trend toward titanosauriform gigantism shifted toward adaptation to dwarfism in some lithostrotian titanosaurs like *Magyarosaurus* and *Neuquensaurus* (see discussion on hind limb size variability below). In general, the results obtained here confirm the previously proposed trend toward the acquisition of robust and arched hind limbs in Titanosauria (Fig. 7). Nevertheless, once hind limb mechanical stability was acquired (following e.g., Ullmann et al., 2017) the increasingly arched and robust morphologies established within Titanosauria cannot be fully related to an increase in body mass (Fig. 6a) and are better explained as convergence between different subclades (Figs. 2 and 6, see evolutionary trend breakdown in Table 7). Saltasaurine lithostrotians are characterized by this type of extreme morphology, with hyper-robust limb bones. Even large saltasaurid sauropods like *Opisthocoelicaudia skarzyinski* exhibit this type of hind limb (Borsuk-Bialynicka, 1977). However, this morphology is not exclusive of saltasaurids, since other titanosaurs exhibit homoplastic hyper-robust and arched hind limbs (i.e., *Dreadnoughtus* and *Magyarosaurus*). This progressively arched limb was probably hard-coded in the macronarian bauplan and, after the somphospondylan stable posture was acquired, was still present but relatable to a significant variability of biomechanical adaptations (Voegele et al., 2021) as our analyses suggest. Despite biomechanical diversity associated with hind limb morphology is still unclear, several studies point out that morphological differences in the fore limb elongation in sauropods may be related to different feeding niche capabilities (Bates et al., 2016; Upchurch et al., 2021; Vidal et al., 2020; Voegele et al., 2020), including discussion on possible bipedal/tripodial rearing abilities that are much more developed, particularly in sauropods with an hyper-robust of hind limb (Upchurch et al., 2021). Interestingly, *Dreadnoughtus* exhibits this hyper-robust hind limb with subequal autopodial lengths and a wedged sacrum (Vidal et al., 2020; Voegele et al., 2021), an anteriorly placed body Centre of Masses (CoM; Bates et al., 2016) and possibly high browsing feeding capabilities (e.g., Vidal et al., 2020). While the small saltasaurids in this study exhibit short necks, stout bodies and a posteriorly placed CoM (Bates et al., 2016) the pneumaticity of some saltasaurid tails (Upchurch et al., 2021; Zurriaguz and Cerda, 2017) may suggest the possibility of a more anterior location of the CoM. It is possible that the acquisition of a hyper-robust and arched hind limb morphology is related to improved skeletal support in high browsing and extremely large sauropods like *Dreadnoughtus*, which in combination with their size and neck dorsiflexion capabilities (e.g., Vidal et al., 2020) allowed them to feed in a high niche stratification, independent of the acquisition of rearing capabilities.

Although PC2 does not provide a phylogenetic signal for the titanosaurian evolution, it reveals significant differences among smaller titanosaurian taxa with robust hind limbs and especially hyper-robust zeugopodia (Fig. 3), like *Magyarosaurus* and the saltasaurines *Neuquensaurus* and *Saltasaurus*, which can be related to different roles in the necessary mechanical stability for rearing capabilities (following Upchurch et al., 2021). *Magyarosaurus*, *Neuquensaurus* and *Saltasaurus* occupy increasingly positive PC2 values (Fig. 3). *Magyarosaurus* exhibits a short zeugopod in which the tibia is slightly laterally rotated, the cnemial crest is laterally projected and the fibula is slightly sigmoidal with its proximal third displaying an anterior projection and a medially deflected anterolateral crest. Saltasaurines exhibit an extreme condition characterized by short and robust zeugopods. Members of this clade show laterally rotated tibiae with posteriorly deflected distal ends, and sigmoidal fibulae that articulate in an oblique position with the anteriorly projected proximal third and anteriorly placed lateral crest. Additionally, this anatomical fibular configuration produces a more anterior displacement of the distal attachment of the *M. iliofibularis* and a probable different distribution of the stress as the main beam of the shafts is rotated from the hind limb trunk, for which previous myological studies suggest that there is no evident body size/phylogenetic pattern (Voegele et al., 2021). In contrast, *Dreadnoughtus* exhibits a slightly arched hind limb posture with a robust zeugopod, a medially deflected cnemial crest of the tibia and a sigmoidal fibula that is slightly posteriorly deflected, and it is projected at negative PC2 (Fig. 3). In this taxon, the fibula, although sigmoidal, articulates in a mostly straight anatomical position with the particularly robust tibia. The differences among robust hind limb titanosaurian morphologies may suggest that their convergence is due to other sources of variation (i.e., different biomechanical adaptations). It is also important to note that all the lithostrotian titanosaurs that exhibit a slightly rotated fibula, whether sigmoid or not, and extremely arched hind limb morphology independent of their size, exhibit femora with reduced fibular epicondyles (after Otero et al., 2020).

Previous studies also pointed to the distinct morphology of the members of Colossosauria, especially in their extremely elongate and gracile zeugopod (González Riga et al., 2018; Páramo et al., 2020). Here we found a distinctive plotting for the rinconsaurian hind limb of *Muyelensaurus* and *Mendozasaurus*, often overlapping with *Aeolosaurus,* as in the PC1 and PC3 morphospaces (Figs. 2-4). It is especially relevant that these specimens occupy distinctive PC1 and PC3 scores within Rinconsauria/Colossosauria, which exhibit a signal regarding titanosaurian evolution and corresponding to a specific lineage of this titanosaurian subclade. In addition, the phylomorphospaces indicate that some lithostrotians shift away from the main early-branching lithostrotian area of the morphospace towards a straighter, elongated hind limb convergent with non-titanosaurian macronarians (Fig 2a-b, Fig. 4a-b). In the case of *Aeolosaurus* the phylomorphospace produces a long, branched and wide morphospace occupation among titanosaurs (Figs. 2-4) as the whole titanosaurian clade re-occupies an entirely new area of the morphospace. Representatives of this group exhibit extremely distinct morphologies, with *Aeolosaurus* and *Antarctosaurus* often much more similar to the early branching non-titanosaurian titanosauriformes than to other deeply-branched titanosaurs (e.g., Fig. 2b, 4b). Colossosauria includes several of the largest titanosaurs ever known (Carballido et al., 2017; González Riga et al., 2019) as well as some of the smallest lithostrotian titanosaurs in South America that are placed within Rinconsauria, which has been recently recovered as an early branching lineage of Colossosauria (Carballido et al., 2017; González Riga et al., 2019). Additionally, some recent phylogenetic hypotheses recovered the aeolosaurine lithostrotians as deeply branching rinconsaurians as well (Carballido et al., 2017; Silva Junior et al., 2022). Whether the appendicular skeleton of Aeolosaurini and Colossosauria exhibits morphological convergence between these two different lithostrotian subclades or they represent the same morphological trend within the same lineage of specialization is still unknown. This depends on whether the latter phylogenetic hypotheses including Aeolosaurini within Colossosauria are accepted as valid. It is important to note that despite the small to medium size of *Muyelensaurus* and *Aeolosaurus*, *Mendozasaurus* can be considered a large sauropod and it exhibits morphological differences from other large sauropods in our sample (i.e., the titanosaurs *Dreadnoughtus* and *Jainosaurus* or the non-titanosaurian somphospondylan *Ligabuesaurus*), similar to previous analyses of separate specimens (Páramo et al., 2020). *Antarctosaurus* seems to be the only large non-colossocaurian titanosaur that shares a similar morphospace with the members of Colossosauria analysed here (i.e., PC1-PC3, Figs. 2-4). The inclusion of hind limb elements of other members of this clade (i.e., *Patagotitan*, *Epachthosaurus*) in future analyses might shed light on this morphological pattern, as the zeugopod elements of several of these taxa resemble those of other non-colossosaur titanosaurs (e.g., *Dreadnoughtus* and *Patagotitan*; see also *Argentinosaurus* in Páramo et al. 2020). Also, *Aeolosaurus* exhibits a different morphology from other post-early Coniacian lithostrotians titanosaurs, thus producing a widening of the occupied morphospace (Fig. 7b-c).

Despite these small differences in the hind limb morphology in key features that may be related to the giant titanosaurian body size, our analyses indicate the great morphological convergence between the different titanosaurian subclades (e.g., *Aeolosaurus* and Colossosauria; or the latter closer to the plesiomorphic morphology of *Euhelopus*, Fig. 2; see also Table 3-4). Also, no significant differences were found in any of the shape variables, either in the pairwise subclade comparisons (Appendix S4), or in the shape PCs that indicate slightly less variance than expected by evolution under Brownian motion (i.e., shape PC1, PC3 in this text, Table 5; shape PC5-6 in Appendix S4).

**Table 4.** Phylogenetic ANOVA test on shape PCA variables between the most inclusive subclades studied.

|  |  |  |  |  |  |  |  |  |
| --- | --- | --- | --- | --- | --- | --- | --- | --- |
|  |  | Df | SS | MS | R<sup>2</sup> | F | Z | Pr(>F) |
| PC1 | ~Clade | 8 | 0.073 | 0.009 | 0.466 | 0.872 | -0.105 | 0.531 |
|  | Residuals | 8 | 0.083 | 0.010 | 0.533 | - | - | - |
|  | Total | 16 | 0.157 | - | - | - | - | - |
| PC2 | ~Clade | 8 | 0.001 | 1.50E+04 | 0.685 | 0.218 | 0.573 | 0.313 |
|  | Residuals | 8 | 5.52E+04 | 6.90E+09 | 0.314 | - | - | - |
|  | Total1 | 16 | 0.002 | - | - | - | - | - |
| PC3 | ~Clade | 8 | 1.54E+04 | 1.93E+09 | 0.285 | 0.399 | -0.925 | 0.82 |
|  | Residuals | 8 | 3.86E+04 | 4.83E+08 | 0.714 | - | - | - |
|  | Total | 16 | 5.40E+04 | - | - | - | - | - |
| PC4 | ~Clade | 8 | 1.49E+04 | 1.87E+09 | 0.3534 | 0.546 | -0.496 | 0.658 |
|  | Residuals | 8 | 2.73E+04 | 3.41E+09 | 0.646 | - | - | - |
|  | Total | 16 | 4.22E+04 | - | - | - | - | - |
| PC5 | ~Clade | 8 | 3.77E+04 | 4.72E+09 | 0.556 | 125.362 | 0.121 | 0.452 |
|  | Residuals | 8 | 3.01E+04 | 3.77E+09 | 0.443 | - | - | - |
|  | Total | 16 | 6.79E+04 | - | - | - | - | - |
| PC6 | ~Clade | 8 | 3.24E+04 | 4.06E+09 | 0.590 | 144.158 | 0.590 | 0.264 |
|  | Residuals | 8 | 2.25E+04 | 2.82E+09 | 0.409 | - | - | - |
|  | Total | 16 | 5.50E+04 | - | - | - | - | - |
|  | ~Clade | 8 | 1.63E+04 | 2.05E+09 | 0.434 | 0.768 | -0.315 | 0.625 |
|  | Residuals | 8 | 2.13E+04 | 2.66E+09 | 0.565 | - | - | - |
|  | Total | 16 | 3.76E+04 | - | - | - | - | - |

**Table 5.** RMA models of the shape PCs against log-transformed Centroid size. CI – confidence interval.

|  | intercept | slope | CI 2.5% Slope | CI 97.5% Slope | r<sup>2</sup> | P |
| --- | --- | --- | --- | --- | --- | --- |
| PC1 | 0.221 | -0.102 | -0.168 | -0.062 | 0.105 | 0.204 |
| PC2 | 0.155 | -0.072 | -0.12 | -0.043 | 0.054 | 0.371 |
| PC3 | 0.137 | -0.064 | -0.106 | -0.038 | 0.055 | 0.363 |
| PC4 | 0.12 | -0.055 | -0.093 | -0.033 | 0.026 | 0.534 |
| PC5 | -0.107 | 0.049 | 0.03 | 0.082 | 0.079 | 0.275 |
| PC6 | -0.1 | 0.046 | 0.028 | 0.076 | 0.086 | 0.254 |

**Table 6.** Estimated phylogenetic signal via Pagel’s lambda. *p* value of log-likelihood ratio test after 1000 simulations. * Indicates significant relationships for an alpha of 0.05. log-Csize – Log-transformed hind limb centroid size.

|  | Lambda | p |
| --- | --- | --- |
| log-Csize | 0.982 | 0.003* |
| PC1 | 0.715 | 0.000* |
| PC2 | 0 | 1 |
| PC3 | 0.76 | 0.002* |
| PC4 | 0 | 1 |
| PC5 | 0.778 | 0.031* |
| PC6 | 0.697 | 0.01* |

**Table 7.**
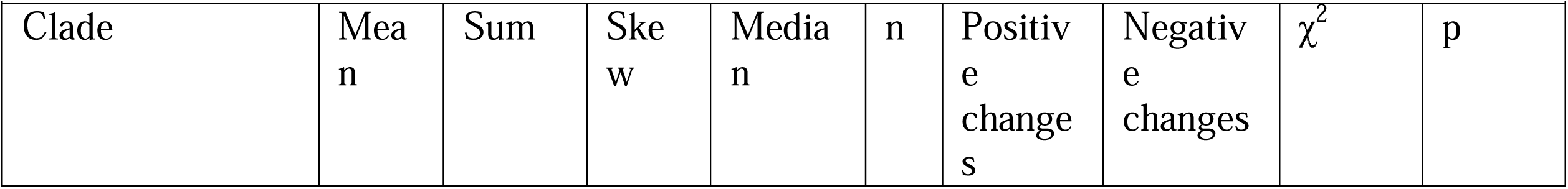
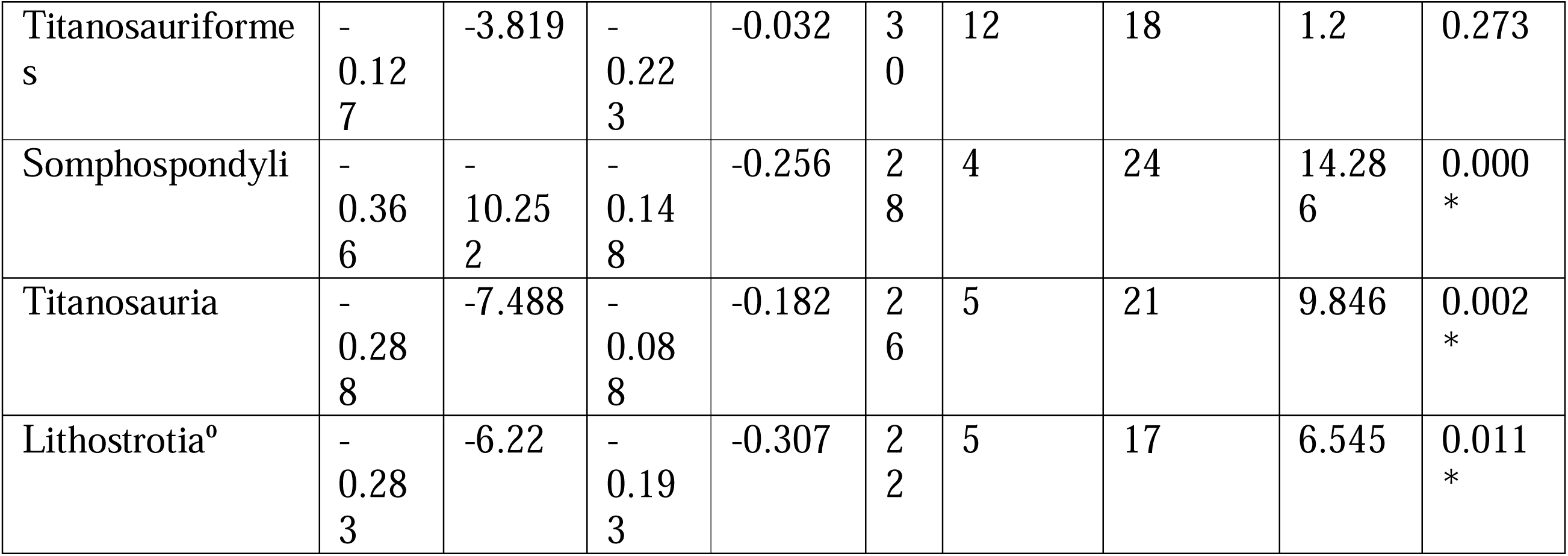
Results of pairwise ancestor-descendant comparisons for log-transformed centroid size in macronarian sauropods in our time-calibrated supertree (17 terminal taxa). n = ingroup internal nodes + terminal taxa. *= accepted as significant with alpha <0.05. °= Lithostrotia + *Antarctosaurus*.

| Clade | Mean | Sum | SKEW | Median | n | Positive changes | Negative changes | $\chi^2$ | p |
| --- | --- | --- | --- | --- | --- | --- | --- | --- | --- |
| Titanosauriformes | -0.127 | -3.819 | -0.223 | -0.032 | 30 | 12 | 18 | 1.2 | 0.273 |
| Somphospondyli | -0.366 | -10.252 | -0.148 | -0.256 | 28 | 4 | 24 | 14.286 | 0.000* |
| Titanosauria | -0.288 | -7.488 | -0.088 | -0.182 | 26 | 5 | 21 | 9.846 | 0.002* |
| Lithostrotia<sup>o</sup> | -0.283 | -6.22 | -0.193 | -0.307 | 22 | 5 | 17 | 6.545 | 0.011* |

### 3.2. Hind limb size evolution

Our results with the current time-calibrated supertree topology indicate that there is a trend in the evolution of titanosaurs towards a decrease in size (Fig. 5 and 7, Tables 5-6). In the light of our current sample, once many of the hind limb features relatable to wide-gauge posture are acquired, there is a phylogenetical trend, close to a pure Brownian motion model, toward an overall decreasing size (log-transformed hind limb centroid size λ = 0.982), which is consistent with previous results that indicated that lithostrotians (or even all macronarian sauropods) may not follow the Cope’s rule (Carrano, 2006; de Souza and Santucci, 2014). This could be due to our current lithostrotian sample as some of the Saltasaurinae or closely related taxa (i.e., *Opisthocoelicauda* and *Alamosaurus*, which are usually considered members of a more inclusive clade, Saltasauridae) are also large lithostrotians, especially *Alamosaurus*. However, it is important to note that we also found several key traits in our shape variables (most importantly summarized in PC1 and PC3, 41.53% of the total sample variance between them; Table 7) that are usually related to the acquisition of gigantism and that exhibit significant signals about titanosaurian evolution. When we tested for a significant correlation between these traits and the log-transformed hind limb centroid size, no significant correlation was found between any of the shape variables and size (Table 4, Fig. 6).

Within Titanosauria, the hind limb arched morphology (increasing wide-gauge posture) does not correspond to the significant trend toward a size decrease observed in this sauropod clade (see Fig. 7, Table 6-7). Large titanosaurs show very different hind limb morphologies. Some lithostrotians exhibit plesiomorphic columnar, slightly arched hind limbs with elongated or even gracile zeugopod elements (*Antarctosaurus*, *Aeolosaurus* or the extremely slender hind limb of the colossosaurians *Mendozasaurus* and *Muyelensaurus*) (Fig. 2), whereas other large titanosaurs (i.e. *Dreadnoughtus)* exhibit an extremely robust hind limb with slightly reduced zeugopods as may be expected from the typical trend toward the acquisition of an arched morphology in Titanosauriformes (Fig. 2a). In this context, it is remarkable that most of the lithostrotians that exhibit extremely arched hind limbs, robust elements and reduced zeugopod elements decrease in size, like *Saltasaurus* or *Magyarosaurus* (Fig. 2, see discussion on morphological convergence above). Our test found that the trend toward this type of hind limb breaks in titanosaurian sauropods, with more variable morphologies and large overlapping of morphospaces whereas the hind limb size decreases (Fig. 7, Table 6-7). Most importantly, PC3 exhibits some of the morphological features that are classically related to wide-gauge arching morphology (e.g.) and our analyses found a significant trend toward increasingly arched hind limbs (Table 8). However, the post-Cenomanian lithostrotians exhibit great morphological variability, with several taxa exhibiting plesiomorphic columnar femora (i.e., *Ampelosaurus*, *Aeolosaurus*, *Magyarosaurus*) and slightly straight zeugopodial elements, with anteriorly expanded tibial proximal ends, and straight non-rotated fibula expanded in the anterior view (Fig. 4,7; see discussion on morphological convergence above). Among these lithostrotians, *Ampelosaurus* and *Aeolosaurus* are medium-sized sauropods, whereas *Magyarosaurus* exhibits both plesiomorphic hind limb morphology and the smallest size. Similarly, the large size variability in the post-early Coniacian cannot be related to the morphological features that are classically associated with the progressively arched morphology (Fig. 7), as discussed before. Therefore, titanosaurs that produce large PC1-PC3 changes at this peak are both moderate-to-large in size, inducing a displacement of the morphospace: (i) toward the plesiomorphic hind limb morphology, plotted in negative PC1 and positive PC3 values (i.e., *Aeolosaurus*) or (ii) toward a morphology intermediate between the main PC1-PC3 typical robust arched titanosaurian hind limb morphology (i.e., *Jainosaurus*) (Fig. 7).

**Table 8.** Results of ancestor-descendant pairwise comparisons for shape PC1 in macronarian sauropods in our time-calibrated supertree (17 terminal taxa). n = ingroup internal nodes + terminal taxa. *= accepted as significant with alpha <0.05. °= Lithostrotia + *Antarctosaurus*.

| Table 8. Results of ancestor-descendant pairwise comparisons for shape PC1 in macronarian sauropods in our time-calibrated supertree (17 terminal taxa). n = ingroup internal nodes + terminal taxa. *= accepted as significant with alpha <0.05. <sup>o</sup>= Lithostrotia + *Antarctosaurus*. |  |  |  |  |  |  |  |  |  |
| --- | --- | --- | --- | --- | --- | --- | --- | --- | --- |
| Clade | Mean | Sum | SKEW | Median | n | Positive changes | Negative changes | $\chi^2$ | p |
| Titanosauriformes | 0.051 | 1.518 | 0.98 | 0.043 | 30 | 29 | 1 | 26.133 | 0.000* |
| Somphospondyli | 0.019 | 0.538 | 1.158 | 0.012 | 28 | 20 | 7 | 6.259 | 0.012* |
| Titanosauria | 0.011 | 0.291 | 1.076 | 0.003 | 26 | 14 | 10 | 0.667 | 0.414 |
| Lithostrotia<sup>o</sup> | 0.011 | 0.24 | 1.079 | 0.002 | 22 | 11 | 10 | 0.048 | 0.827 |

**Table 9.** Results of ancestor-descendant pairwise comparisons for shape PC3 in macronarian sauropods in our time-calibrated supertree (17 terminal taxa). n = ingroup internal nodes + terminal taxa. *= accepted s significant with alpha <0.05. °= Lithostrotia + *Antarctosaurus*.

| Table 9. Results of ancestor-descendant pairwise comparisons for shape PC3 in macronarian sauropods in our time-calibrated supertree (17 terminal taxa). n = ingroup internal nodes + terminal taxa. *= accepted as significant with alpha <0.05. <sup>o</sup>= Lithostrotia + *Antarctosaurus*. |  |  |  |  |  |  |  |  |  |
| --- | --- | --- | --- | --- | --- | --- | --- | --- | --- |
| Clade | Mean | Sum | SKEW | Median | n | Positive changes | Negative changes | $\chi^2$ | p |
| Titanosauriformes | -0.035 | -1.047 | 1.697 | -0.038 | 30 | 2 | 28 | 22.533 | 0.000* |
| Somphospondyli | -0.023 | -0.639 | 2.174 | -0.024 | 28 | 2 | 26 | 20.571 | 0.000* |
| Titanosauria | -0.008 | -0.21 | 2.483 | -0.009 | 26 | 3 | 23 | 15.385 | 0.000* |
| Lithostrotia<sup>o</sup> | -0.003 | -0.074 | 2.323 | -0.004 | 22 | 4 | 17 | 8.048 | 0.005* |

Here it is important to point out that when comparing our results with other lithostrotian titanosaurs not included in the current analysis, similar hind limb posture variability not relatable to hind limb size increases (and therefore body size) is observed. The arched morphology with extremely robust zeugopodial elements exhibited in saltasaurid sauropods (e.g., *Saltasaurus*) is also found in the dwarf non-lithostrotian titanosaur *Diamantinasaurus* (Poropat et al., 2021, 2015). Its tibia shares similarities with those of *Dreadnoughtus* (anteroposteriorly and lateromedially wide proximal end, extremely short and laterally-projected cnemial crest and lateromedially expanded distal end) (Poropat et al., 2015; Ullmann and Lacovara, 2016) whereas the fibula is similar to the slightly straight fibula of *Magyarosaurus*, with a proximally anterior deflection (Poropat et al., 2015; APB direct observation on *Magyarosaurus* sp. specimens). Despite slight differences in the proportion of the zeugopodial elements, the hind limb morphology of *Diamantinasaurus* is similar to those of other small taxa. However, *Rapetosarus* exhibits a completely different lithostrotian hind limb configuration that is more similar to that observed in *Lirainosaurus*. Thus, whereas the fibula is slightly straight with an anteriorly expanded proximal third (Curry Rogers, 2009) as in *Magyarosaurus* and *Diamantinasaurus*, both the tibia and fibula are extremely lateromedially compressed as in *Lirainosaurus* or *Muyelensaurus* (APB pers. obs.). Our results are congruent with the observation in other, non-sampled small titanosaurian taxa that exhibit similar morphological variability. Notice that *Rapetosaurus* is based on juvenile and subadult specimens and it may retain a plesiomorphic slender morphology that change in the adult (see precocial development in the limbs of *Rapetosaurus*) (Curry-Rogers et al. 2016)

Among large titanosaurs, *Elaltitan* exhibits a (virtually restored) robust femur, a slightly robust and plesiomorphic straight tibia but with a posteriorly rotated distal end, and an extremely sigmoidal and anteriorly projected fibula like those of members of Saltasaurinae but slightly lateromedially narrow compared to the latter (Páramo et al., 2020). Despite some differences, other robust large lithostrotians (i.e., *Dreadnoughtus*) exhibit similar hind limb morphology. However, other large lithostrotians, like *Argentinosaurus,* exhibit a plesiomorphic straight fibula (i.e., Páramo et al., 2020). The hind limb of *Patagotitan* exhibits a less arched posture, with a slightly straighter femur whereas the fibula is robust, sigmoidal and has an anterior expansion of the proximal third (Otero et al., 2020). The stylopodium is not as robust and arched as in *Dreagnoughtus* but also exhibit the slightly plesiomorphic titanosaurian straight hind limb, which is also part of the shift toward robust morphology after the earlier branching Colossosauria, such as *Mendozasaurus* (Figs. 2 and 4). *Patagotitan* still lacks the extreme medial deflection of the femoral head seen in our PC1 positive and PC3 negative values. *Petrobrasaurus* and *Narambuenatitan* are both large titanosaurs that exhibit similar medial deflection of the femoral head to *Dreadnoughtus*, and in the case of *Narambuenatitan*, even more than *Saltasaurus* and *Neuquensaurus* (Páramo et al., 2020). However, the tibia of *Petrobrasaurus* is extremely lateromedially narrow as in *Mendozasaurus* (Páramo et al., 2020; APB pers. obs.) instead of the typical robust tibiae of the extremely arched hind limb found in our results (see Fig. 2). Only *Uberabatitan* exhibits a clear morphology like the hind limb morphology found in *Dreadnoughtus* and our positive PC1 scores. Another large titanosaur that exhibits a lateromedially narrow proximal tibial end is *Ruyangosaurus* (Lu et al., 2009). Despite being fragmentary, the femur is straight (with a rounded shaft) and has a tibia with lateromedially narrow proximal end similar to that observed in *Ligabuesaurus* and the members of Colossosauria (which are closely positioned in our analyses) (Figs. 2 and 4). Other more deeply branching lithostrotians either exhibit a hind limb that closely resembles the early-branching colossosaurs or a plesiomorphic tibia as in *Abditosaurus* (Vila et al., 2022). In this taxon, the femur is anteroposteriorly-narrow with a highly eccentric shaft, like other Ibero-Armorican lithostrotians (e.g., *Ampelosaurus*). *Abditosaurus* also preserves an extremely lateromedially narrow tibial proximal end (Vila et al., 2022). Only the fibula is sigmoid and anteriorly projected, but lateromedially narrow with an expanded anteromedially crest, which is larger anteroposteriorly than in *Lohuecotitan*, *Lirainosaurus* (Vila et al., 2022) and probably like those of *Magyarosaurus*, and is the single character that resembles our PC1 results.

Moreover, we must consider that our age estimations for the nodes of our supertree are extremely conservative and are based on the topology after the reduced tips of our sample (see Table 1). Titanosaurian sauropods appeared unambiguously during the Early Cretaceous (D’Emic, 2012; Mannion et al., 2019b; Poropat et al., 2016) just as lithostrotian titanosaurs appeared early after the Valanginian-Hauterivian (Mannion et al., 2019b; Poropat et al., 2016).

Considering several recent phylogenetic hypotheses, the colossosaurian node may have branched in the early Albian (Gorscak et al., 2023; Sallam et al., 2018; Vila et al., 2022). Saltasauridae have also often been estimated at the transit between the Early Cretaceous and Late Cretaceous, with *Jiangshanosaurus* considered an early-branching saltasaurid (Poropat et al., 2016). However, the recent redescription of *Jiangshanosaurus* material has shed light on its phylogenetic affinities and it may be an euhelopodid or at least not as a deeply branched titanosaur (Mannion et al., 2019a). Saltasaurid lithostrotians are nevertheless traced back to the late Albian in recent studies (Sallam et al. 2018; Vila et al., 2022). The trends observed in our study may be accentuated with still a general body size decrease among lithostrotian sauropods (Fig. 7a). However, a series of convergences in the appendicular skeleton: e.g., the acquisition of the plesiomorphic columnar titanosaur hind limb among medium to large lithostrotians, seem to evolve independently of body size (e.g., Fig. 7b). Recent phylogenetic hypotheses show uncertain phylogenetic affinities for some of the sauropods studied, particularly for some of the lithostrotian taxa. This is the case of the opisthocoelicaudiine affinities that have been suggested for some members of Lirainosaurinae and *Lohuecotitan*, which is also proposed as a subclade within Saltasauridae (Vila et al., 2022) or Saltasauroidea (Mocho et al. 2024a) as Opisthocoelicaudiinae. *Atsinganosaurus* an Ibero-Armorican lithostrotian was recently recovered as member of Lirainosaurinae (Díez Díaz et al., 2018, 2020, Mocho et al. (2024a)) or as a lognkosaurian colossosaur (Gorscak and O’Connor, 2019; Vila et al., 2022). *Atsinganosaurus* is a small lithostrotian with a hind limb morphology similar to that of *Lirainosaurus* (Díez Díaz et al., 2018). The extremely gracile limbs of *Atsinganosaurus* resemble those of the small rinconsaurian lithostrotians (González Riga et al., 2019; Pérez Moreno et al., 2023 this study) or even the large *Mendozasaurus* (González Riga et al., 2018). However, its affinities to Colossosauria will still indicate a convergence between small Opisthocoelicaudiinae and lognkosaurian colossosaurs according to the phylogenetic hypothesis of Vila et al. (2022). This phylogenetic hypothesis still indicates that the large-sized titanosaurs like *Dreadnoughtus* and *Alamosaurus* (robust medially bevelled femur and sigmoid fibula; see Lehman and Coulson, 2002; Wick and Lehman, 2014) exhibit a morphologically convergence in the hind limb with the small saltasaurines like *Neuquensaurus* and *Saltasaurus*.

It seems that these morphological similarities are due to other biomechanical aspects after the acquisition of the arched hind limb within Somphospondyli, as well as to other morphological features related to the wide-gauge posture of the appendicular skeleton. It is possible that minor differences that do not show a significant phylogenetic signal and recover in other shape-PCs (Appendix S4), are key features for other adaptations that are also important such as the trade-off between speed and rearing stability, which may have shaped limb morphology of Titanosauriformes, particularly in the zeugopods (e.g., Upchurch et al., 2021).

### 3.3. Caveats of this study

Our analyses include a wide range of titanosaurs from most of the proposed subclades. However, many hind limb elements are fragmentary and required virtual restoration according to Páramo et al. (2020, 2022). Traditional studies propose excluding incomplete specimens from the sample. However, in this case, several of our taxa from an already small sample (n = 17) could not be included in this study because they do not have complete specimens to calculate the mean shape for each hind limb element type (e.g., *Mendozasaurus* with several tibia and fibula specimens, *Oceanotitan* with only the fragmentary elements of its left hind limb). The exclusion of potentially informative areas or taxa may hinder palaeobiological studies (Brown et al., 2012) and landmarks estimation may be a more informative procedure (Arbour and Brown, 2014; Brown et al., 2012). Also, it may be interesting to include several taxa that have been examined in previous studies (e.g., Páramo et al., 2020), but many of these taxa lack one or more hind limb element types. In this case, we did not choose to estimate the entire morphology of an element type, because we lack the necessary and more powerful tools to do so, such as partial least squares estimation methods (e.g., Torres-Tamayo *et al*. 2020). The virtual restoration does not appear to contain large artifacts due deformation of the specimens, and most of the sample is not affected by extreme taphonomic artifacts. Some complete specimens may affect the results nonetheless, as it is common that some shaft might be more eccentric due to crushing (i.e., *Ampelosaurus atacis*, could be even closer to *Lirainosaurus astibiae* in PC1) or the deformation of distal condyles in *Dreadnoughtus schrani* which affects its extreme position in PC2. Despite this, we assessed potential biases following Lefebvre *et al*. (2020; Appendix S2-3), and they do not affect significantly any of the shape variables which exhibit phylogenetic signal (e.g., PC2; Table 6; see in details appendix S3).

Our reconstructions of the analyzed titanosaurian hind limbs can also bias our study. The lack of the astragalus in most of the titanosaurs studied (whether due to the lack of available 3D-scanned specimens or the lack of a preserved astragalus) hinders the estimation of the position of the zeugopodial elements. This study uses the most conservative assumption in the overall position of the distal part of the hind limb as the fibula could be positioned even more distally, without reaching proximally the femur, and interlocked with the tibia in several specimens (e.g., *Neuquensaurus australis*, specimen MCS-5-25/26, APB. pers. obs.). This is especially relevant for the robust and arched hind limb with extremely robust zeugopodia, as *Dreadnoughtus schrani*, *Neuquensaurus* spp., and *Saltasaurus loricatus*, but see also *Uberabatitan riberoi* (Salgado and Carvalho, 2008).

The sample is also small and we did not opt to include several advanced statistical hypothesis tests like phylogenetic convergence (Stayton, 2015) because it did not meet the requirements. Instead, we chose a conservative set of tests to discuss the true morphological convergence over titanosaurian hind limb evolution with a mix of phylogenetic and non-phylogenetic methods. We chose Butler and Goswami (2008) change frequencies analysis because, although its statistical properties are less well known than those of independent contrast analyses, it is less sensitive to topological imprecision and somewhat independent of branch length differences (Butler and Goswami, 2008; de Souza and Santucci, 2014). Following this reasoning, we also decided not to estimate independent phylogenetic contrasts that may be sensitive to large differences in branch length in our lithostrotian-biased sample to test for trait correlation between shape-PCs against log-transformed centroid size, contrary to previous studies (Bates et al., 2016). Instead, we opted for the traditional use of test for correlation between tree tips (specimens) shape-PCs against log-transformed centroid sizes via RMA and without incorporating the phylogenetic tree topology of our current time-calibrated supertree.

## 4. Conclusions

Our results suggest that the main features related to the acquisition of an arched hind limb posture (presence of lateromedially wide femora with robust zeugopods) are typical of more deeply nested titanosaurs such as saltasaurines (Carrano, 2005; Lefebvre et al., 2022; Sander et al., 2011; Ullmann et al., 2017; Vila et al., 2022; Wilson and Carrano, 1999) and exhibit a significant phylogenetic signal about titanosaurian evolution (Table 5, Fig. 7). The arched morphology usually related with wide-gauge posture is an exaptation initially related to increasing body size. However, once fully acquired within Somphospondyli, these features are no longer related to increasing body size, as increases in the arched posture and the development of hyper-robust zeugopods are features that evolved independently in several lineages, and are shared by several of both the smallest and largest taxa in different titanosaurian lineages (Fig. 6, Table 4). Also, there is an evolutionary trend toward decreasing titanosaurian hind limb size (and body size; Fig. 7), based on the size of the hind limb centroid. This trend is congruent with previous studies focused on the size evolution within Titanosauriformes (de Souza and Santucci 2014), despite an increase in arched morphology and robustness of the hind limb in deeply nested titanosaurian subclades.

The lack of correlation between the arched posture, position and robustness of zeugopod, among other traits with the body size may also be related to a wide morphological variation within Lithostrotia (Figs. 2-4). It can be noted that, together with a trend toward increasingly arched hind limbs and reduced zeugopods with increasingly rotated fibulae, there is also a large morphological convergence between the different titanosaurian subclades, as indicated by both our non-phylogenetic and phylogenetic analyses (Table 2-3). Many of the morphological changes without a significant signal in titanosaurian evolution are related to the morphology and anatomical position of zeugopod bones. Large morphological convergences unrelated to body size may be a response to different morphofunctional adaptations. Our results show several differences in the morphology of zeugopod elements without a signal in the evolution of titanosaurs, especially those regarding the fibula (rotation, sigmoidal morphology, anterior or posterior displacement of the proximal end that affects the anteroposterior position of the lateral bulge). These morphological differences are also related to its relative position and articulation with the tibia and correlate with the expression of femoral features like the relative development of the posterior epicondyle of the fibular condyle. These features translate in changes in the length and morphology of the hind limb distal musculature that exhibit no clear evolutionary trend within titanosaurs, as previous studies have indicated. Also, our results show that specimens in different subclades share similar morphologies across the shape-PCs that exhibit no phylogenetical signal in titanosaurian evolution. The observed changes in the zeugopod morphology may be related to different morphofunctional and ecomorphological adaptations and the convergences in hind limb morphology of titanosaurs may explain biomechanical similarities. This may explain the differences observed between small and medium-to-large titanosaurs with either arched or columnar hind limbs (i.e., PC1 similarities) across those shape-PCs that lack a significant signal in the evolution of titanosaurs. However, to test the hypothesis of differences in biomechanical adaptation (i.e., movement speed or differences in feeding niche specialization) further analyses with additional parts of the skeleton must be included.

## Supporting information

Appendix S1 revised

Appendix S2 revised

Appendix S3 revised

Appendix S4 revised

Appendix S5 revised R code

## 5. Acknowledgments

This work is funded by Ministry of Science and Innovation of Spain project PID2019-111488RB-I00 and PID2023-148083NB-I00, and the Junta de Comunidades de Castilla-La Mancha projects (SBPLY/19/180801/000044 and SBPLY/21/180801/000045). Access to fossil vertebrate collections in Argentina was made possible by Spanish Ministry of Economy and Competitiveness grant EEBB-I-16-11875. The authors would like to thank BP. Kear and Uppsala University Natural History Museum staff for facilitating access to their collections. Access to Uppsala University Museum of Natural History paleovertebrate collections was possible thanks to the ERG-2020 EAVP research grant. Author PM was funded by FCT/MCTES for one CEECIND/00726/2017/CP1387/CT0034 individual contract (https://doi.org/10.54499/CEECIND/00726/2017/CP1387/CT0034) and SFRH/BD/68450/2010 PhD scholarship. The authors are very grateful to S. Chapman and Natural History Museum of London (UK) staff for facilitating access to their collections. Access to NHM (UK) was made possible thanks to the European Union Council Synthesys program GB-TAF-6153. They also want to thank the students of the Faculty of Arts in the UCM (Madrid, Spain) and preparation staff for providing the sample for this study. They equally thank many others for access to specimens under their care, including S. Langreo at the Museo de Paleontología de Castilla-La Mancha (Cuenca, Spain); J.A. Ramírez de la Peciña and C. Corral at the MCNA (Vitoria, Spain); R. Coria at the MCF (Plaza Huincul, Argentina); L. Filippi at the MRS (Rincón de los Sauces, Argentina); M. Reguero at the MLP (La Plata, Argentina); A. Otero at CONICET (La Plata, Argentina); S. Devincenzi at the IANIGLA (Mendoza, Argentina); and B. González Riga at the UNCUYO (Mendoza, Argentina); C. Muñoz at the MPCA (Cipolletti, Argentina); I. Cerda fromat the CONICET (Cipolletti, Argentina); P. Ortiz at the PVL (Tucumán, Argentina); A.G. Kramarz and M. Ezcurra at the MACN (Buenos Aires, Argentina),J. Le Loeuff at the Musée des Dinosauries d’Espéraza (Espéraza, France) and B. Silva at the SHN (Torres Vedras, Portugal).. We want to ask M Christopher for access to *D. schrani* skeleton 3D scan. We want to thank P.M. Sander, the editorial team and two anonymous reviewers for their comments which helped improve this manuscript.

## 7. Methods

### 7.1. 3D Geometric Morphometrics

To analyse the morphology, 17 macronarian sauropod hind limbs (Table 1) were 3D-digitized and anayzed using 3D Geometric Morphometrics tool-kit (3D-GMM; Gunz et al., 2005 e.g., Páramo et al., 2020). The 3D digitizing process was based on the methodology proposed in previous analyses (i.e., Mallison, 2011; Páramo et al., 2020) and 3D-GMM analyses were conducted in R statistical software v4.1.3 (R Core Team, 2022).

To reconstruct each taxa hind limb, we first virtually-restored the 3D reconstructions of the digitized femora, tibiae and fibulae and calculated the grand mean shape for each sampled taxon using previous datasets (Páramo et al., 2020). The grand mean specimens for each taxon were mounted on an estimated anatomical position (accounting for the lack of astragalus in our sample to guide the distal zeugopod articulation). A comprehensive description of the methodology can be accessed in Appendix S2.

Each element of the hind limb is separated by an articular cap much larger than in other known extant archosaurs (Bonnan et al., 2013; Holliday et al., 2010; Schwarz et al., 2007; Voegele et al., 2022). The femur, and the hind limb overall, may exhibit the best correlation between cartilage cap and bone morphology constrained by its role as support under most of the stress of the body mass (Bonnan et,al., 2013; Voegele et al., 2022). Although cartilaginous cap thickness is not constant and may vary among taxa, we set up our model hind limbs with a similar constant space between the femur and the zeugopod bones in all the sampled taxa. The anatomical mounts include an additional space of 2% of the element length between stylopod and zeugopod (following Voegele et al., 2022).

A total of 28 landmarks and 12 semilandmark curves were placed on the hind limb bones partly based on previous studies (Lefebvre et al., 2022; Páramo et al., 2020; Table 2, Fig. 2, Appendix S1) using IDAV Landmark Editor software (Wiley et al., 2005) (dataset with the landmark and semilandmark curves coordinates can be accessed in the Appendix S1. Semilandmarks were then slid in R using package “Morpho” (Schlager, 2017) following Gunz et al., (2005). To remove size differences, spatial position and orientation, the resulting landmark and semilandmark configurations were superimposed via Generalized Procrustes Analysis (GPA) using the “procSym” function in *Morpho* package. Morphological variance was analysed with Principal Component Analysis (PCA; results in Table 3), saving the expected number of Principal Components (PCs) which summarize a significant amount of variance after an Anderson Chi’s test (see Bonnan, 2007).

Evolutionary trend analyses were accomplished after the estimation of a consensus tree topology using the MRP-supertree methodology (Bininda-Emonds, 2004) with the *phangorn* package (Schliep et al., 2017; Schliep, 2011). For supertree construction we compiled several of the more recent phylogenetic hypotheses that include all the available sampled data (resulting supertree can be accessed in Appendix 1; trimmed supertree in Fig. 2). The resulting phylogenetic relationships were projected onto the shape-PCA to visualize the phylomorphospace. To analyse true morphological convergences between titanosaurian sub- clades, we tested for morphological differences in the hind limb skeleton using the shape variables (PCs) without the phylogenetic relationships involved using: (i) Mann Whitney U’s test and Kruskal-Wallis non-parametric tests accounting for the uneven distribution of the group (sub-clade) samples and (ii) phylogenetic ANOVA using the time-calibrated supertree topology with the *phytools* R package (Revell, 2012).

### 7.2. Hind limb size distribution and phylogenetic signal

Sauropod body mass was proxied by hind limb centroid size collected from the GPA. Body mass can be calculated preferably using both humeral and femoral measurements (Mazzetta et al., 2004) or a whole-body volumetric estimation (Bates et al., 2016). However, as the hind limb is the main sauropod body mass support, it can be better used as a “conservative-minimal” approach to its body mass, with larger hind limb corresponding to giant titanosaurian taxa. We tested for allometric relationships between the sauropod hind limb shape variables (PCs) and the centroid size as proxy to body mass via Reduced Major Axis (RMA) regression using *lmodel2* R package (Legendre, 2018); but an alternative set of analyses were carried out using femoral length and body mass estimations (see Appendices S2-4).

We used the time-calibrated supertree topology and our shape variables (PCs) and hind limb centroid size to generate the Pagel’s lambda (λ) with the “phytools” R package and test for the phylogenetic signal of these traits. All hypotheses of statistical correlations, dissimilarity tests and phylogenetic signal tests were accepted as significant using an alpha level of 0.05 (a comprehensive report of the results and a copy of the R code and packages used can be accessed in Appendix S4 and S5 respectively). Once those PCs that exhibit significant phylogenetical signal were identified, as well as the log-transformed centroid size, we estimated their ancestral characters (ACEs) using maximum-likelihood and a simple Brownian evolutionary model similar to the one assumed for estimation of the Pagel’s lambda. We used the ACEs to observe trends in the evolution of titanosaurian hind limb size and morphology (based on our shape- PCs). We evaluated the differences in body size (proxied by log-transformed hind limb centroid size ACEs) between terminal taxa and internal nodes and between internal nodes of Titanosauriformes, Somphospondyli, Titanosauria and Lithostrotia, as well as their subclades. The sum of changes, mean change, median change, positive, negative and the total amount of changes were evaluated for each of the above clades following Butler and Goswami (2008). We used a χ2 goodness-of-fit test to evaluate whether body size increasing or decreasing, and shape- PCs occur at the same frequency (50%-50% null hypothesis) or have a positive or negative tendency over titanosaurian evolution.

