## Appendix S2 revised for "Evolution of hind limb morphology of Titanosauriformes (Dinosauria, Sauropoda) analyzed via 3D Geometric Morphometrics reveals wide-gauge posture as an exaptation for gigantism"

Electronic Supplementary Material

Appendix S2 – Methodology

In this appendix we will describe comprehensively the methodology used in this study for reconstructing the 3D digital morphology of the hind limb for each sampled taxon, the Virtual Restoration of the incomplete hind limbs following our previous workflow (Páramo et al., 2020). The virtual restoration and the shape analyses were all carried out via landmark-based 3D Geometric Morphometrics (3D-GMM; e.g., Gunz et al., 2005; Rohlf, 1999). The shape analyses were complemented by a series of tests using a phylogenetic ingroup label (“clade” as per most-inclusive Titanosauriformes subclades). The evolutionary analyses were carried out using the shape variables obtained from the 3D-GMM analyses and a summary of current phylogenetic hypotheses via supertree methodology (Bininda-Emonds, 2004). Using the summary phylogenetic supertree and the shape variables we estimated the ancestral characters (ACE; Pagel, 1999; Pagel et al., 2004), the phylomorphospaces and tested for changes in the hind limb shape across the summary phylogenetic supertree.

We selected a random sample of 16 titanosauriform sauropods and *Oceanotitan dantasi* as representative of non-titanosauriform Macronaria as an outgroup for our study and base for all the evolutionary comparison (17 macronarian sauropod hind limbs in total). The study is slightly biased toward Lithostrotian sauropods nonetheless (see Main Text, potential biases in our results) but we took precautions and conservative conclusion based on our current sample of macronarian sauropods.

### S2.1. 3D digitizing and reconstruction

#### S2.1.1. Specimen 3D Digitizing

For each sampled titanosauriformes taxon of this study, all the available specimens of each bone element were digitized individually. The 3D digitizing of each individual specimen was carried by stereophotogrammetry following previous workflows (Díez Díaz et al., 2021b; Mallison, 2010; Páramo et al., 2020; see also Fig.S2.1). The sequences of pictures had been taken with a Canon EOS 1100D DSLR and a Canon EOS 80D DSLR and Canon 18-55mm f3.5, Canon 50mm f1.8 and Sigma 17-50mm f2.8 lenses. All the lenses used except for Canon 50mm f1.8 produce a noticeable distortion (e.g., Collins and Gazley, 2017) so the pictures are in undistorted RAW format with the specimen centred far from the margins were distortion is concentrated and the variable focus lenses were set to 35mm, as in these focal length most of the lens distortion is reduced. The pictures were processed with Agisoft Metashape® v.1.8.1 and the resulting 3D model reconstruction of each fossil specimen exported as OBJ for retopology under Instant Meshes (Jakob et al., 2015) and mesh correction in Blender® v.2.79b (Blender Online Community, 2018).

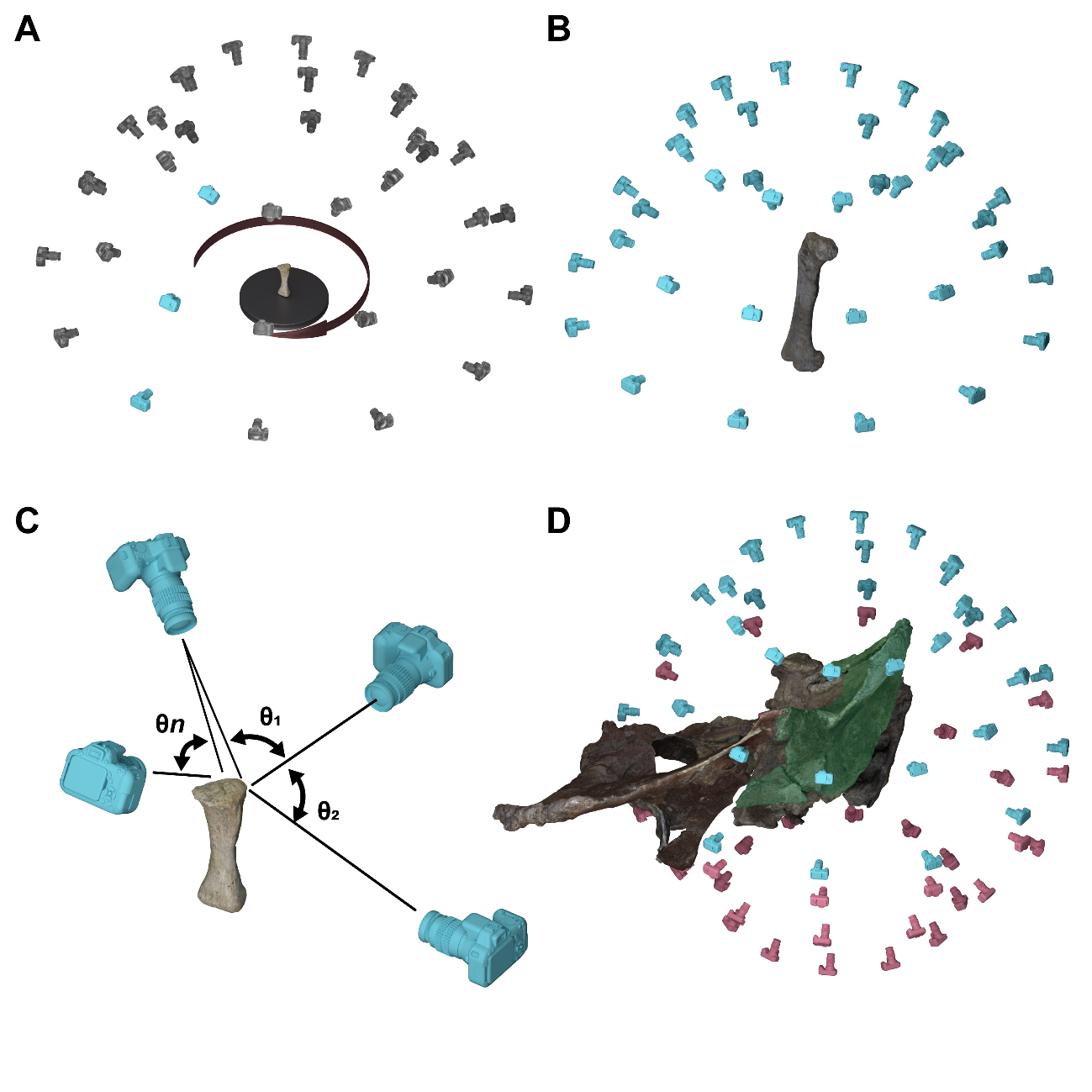

Fig. S2.1. Stereophotogrammetry protocol. A) turntable method of photographing the specimen for small specimens. B) turn-around method for larger specimens C) angles between different pictures to recreate the pixel point-cloud in 3D space in the software D) partial digitizing of separated fragments of the same specimen.

#### S2.1.2. Hind Limb reconstruction

Each element type was sampled with the same set of landmarks and semilandmark curves defined Páramo et al. (2022, 2020; Fig.S2.2 and 3). The landmark and semilandmark curves were sampled in IDAV Landmark Editor® v.3.66 (Wiley et al., 2005) using the Atlas template method (Botton-Divet et al., 2015) and a custom hypothetical macronarian hind limb 3D model used for defining the landmarks. The landmark and semilandmark curves were imported into R statistical software v4.1.3 (R Core Team, 2022) with the packages *geomorph* (Adams et al., 2019) and *Morpho* (Schlager, 2017).

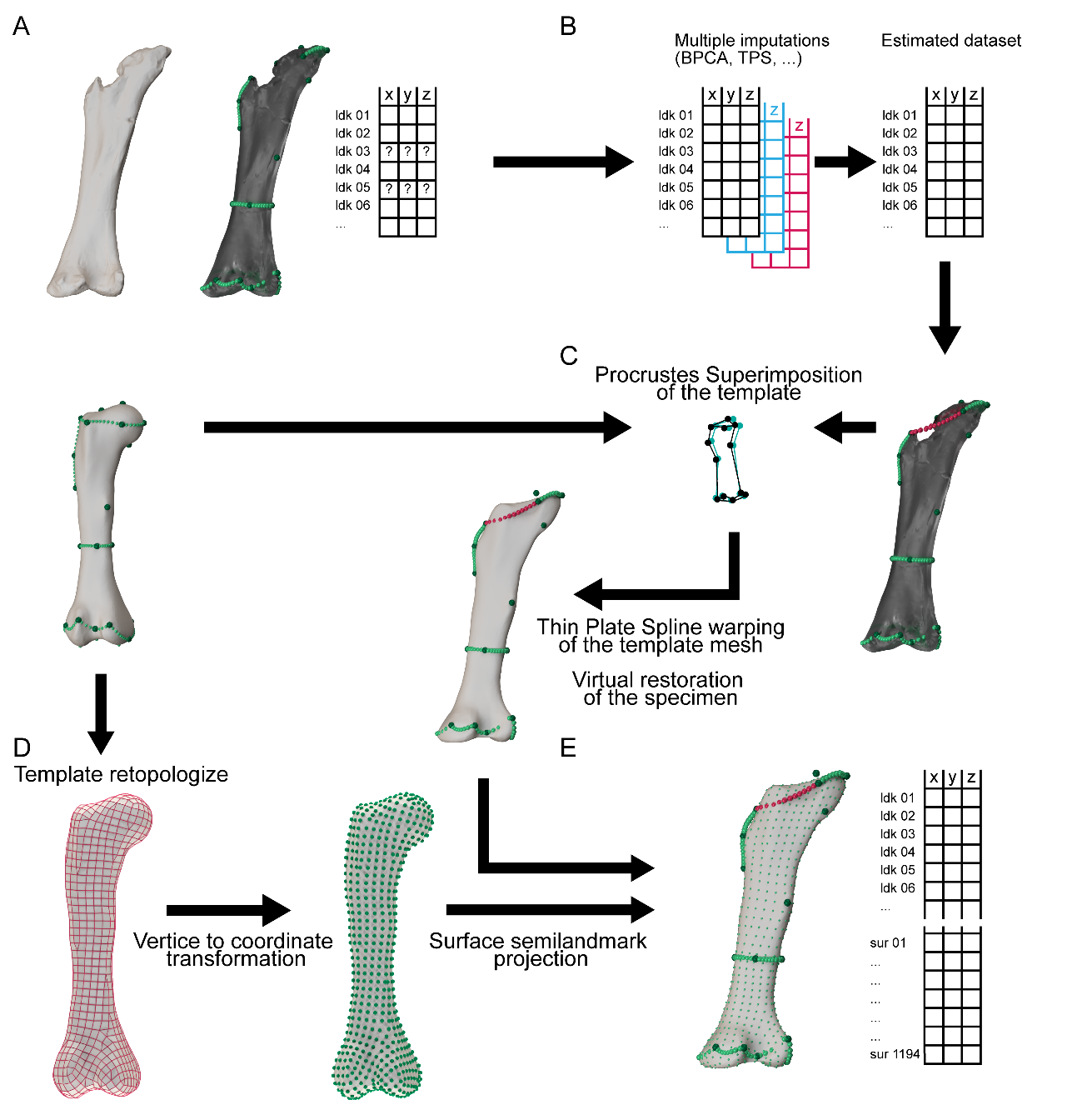

Fig. S2.2. Separate specimen Statistical Virtual Restoration workflow as in Páramo et al. (2020). A) Initial 3D specimen reconstruction landmark sampling B) Multiple imputation methods (TPS in this case) C) Procrustes superimposition of the template mesh and the estimated landmark configuration in order to obtain a reconstructed specimen 3D mesh for anatomical mount.

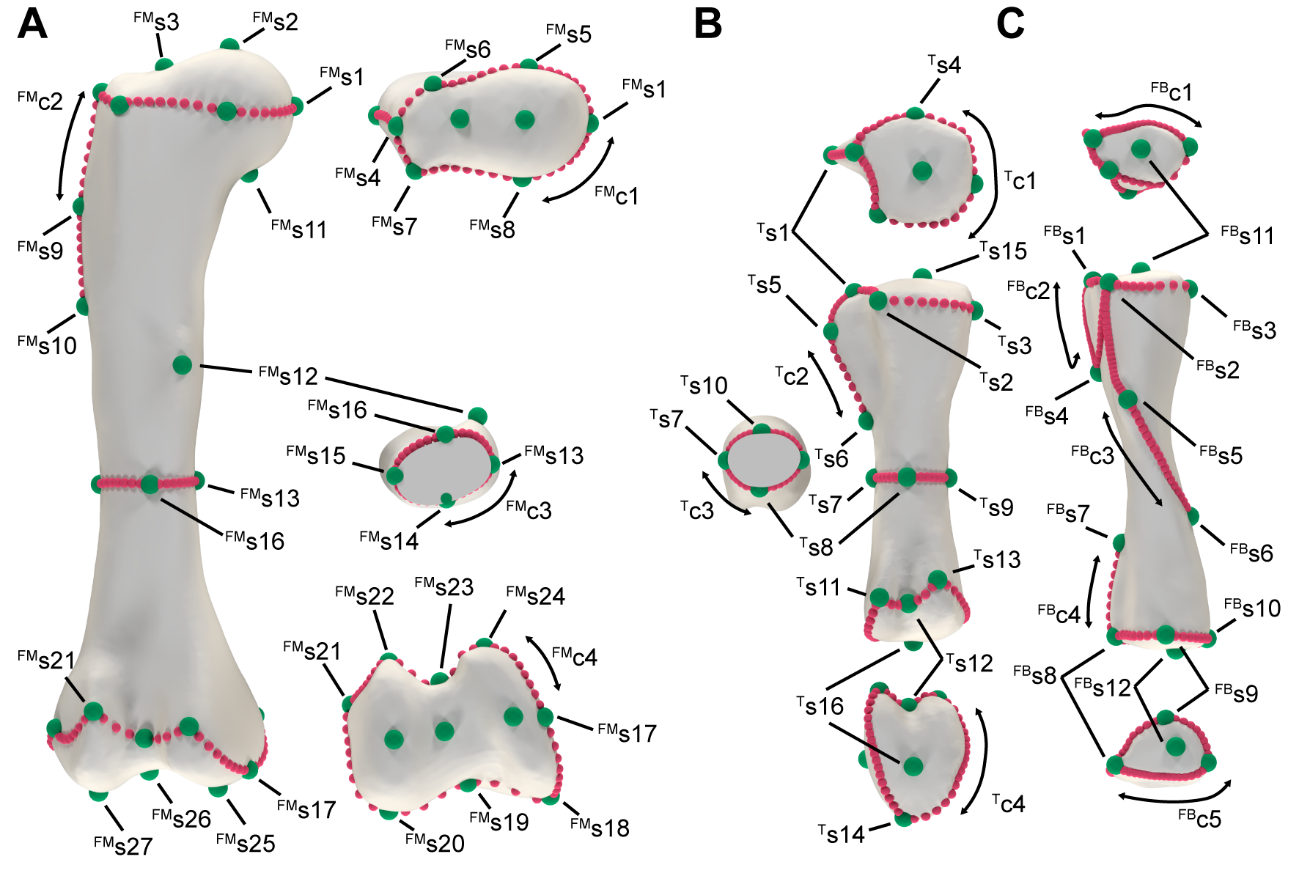

Fig.S2.3. Landmarks and semilandmark curves used for the separated specimen prior landmark estimation during Statistical Virtual Restoration procedure. A) femur B) tibia C) fibula.

Not all the available specimens are complete and we opted for estimation of missing information instead of excluding them form study as the available sample would not be possible to analyse with a rationale conclusion (see considerations prior to missing data estimation instead of specimen exclusion in Brown et al., 2012). Therefore, in order to obtain a “mean shape” for each taxon we firstly need a Virtual Restoration procedure using the same techniques as per the 3D-GMM. We opted for a statistical virtual restoration procedure instead of manually sculpting or undistort a specimen in a 3D modelling software in order to not increase the errors and biases derived from the author manipulation of the starting data prior to the analyses. The single exception was made with *Oceanotitan dantasi* femur (fragmentary, lacking part of the shaft, total length unknown; Mocho et al., 2019). We first estimated the complete length based on several femoral measurements (e.g., lateromedial width of the proximal end of the femur, the lateromedial width of each distal condyle, the proximodistal length of the lateral bulge, etc.) taken in the entire sample of macronarian sauropod specimens as well as some Ibero-Armorican morphotypes included in previous analyses (supplied in Appendix S1). With this linear morphometric dataset we estimated the missing measurements using multiple imputation methods and Predictive Mean Matching (pmm; Allison, 2000; Morris et al., 2014; Royston, 2004) with *mice* package (van Buuren and Groothuis-Oudshoorn, 2011). Using the estimated length, we placed the preserved 3D-digitized reconstruction of the proximal and distal ends of the femur of *Oceanotitan dantasi* in their corresponding positions in Blender®. The resulting femur lacking the shaft but with the proximal and distal end in their estimated position was exported as OBJ after retopologizing and mesh reparation similar to other specimens. The lacking landmark and semilandmark curves, especially concentrated in the missing shaft, were estimated like the other specimens of the sample within R, no manual-resculpt of the missing shaft was carried.

The landmark estimation was carried with the same Thin Plate Spline used for semilandmark sliding (TPS; Brown et al., 2012; Gunz et al., 2009, 2005) using *LOST* package (Arbour and Brown, 2017) and *geomorph* (Fig. S2.2). We obtained a “mean shape” of each element type for each taxon of our sample using the estimated complete set of landmarks and semilandmark curves. We also reconstructed the complete 3D morphology for each element type and sampled taxon warping the template hypothetical 3D mesh to the landmark configuration of each macronarian sauropod. The resulting 3D models were used to reconstruct the macronarian hind limbs, mounting in Blender® in their anatomical position. The femur slightly bevelled to achieve a neutral anatomical position were the distal end offers a somewhat straight plane surface for articulation with the tibia and fibula. The tibia is straight but slightly rotated in proximal view to accommodate the fibular anterior trochanter in the proximal end with the space between the cnemial crest and the shaft (or the cnemial crest and the secondary cnemial crest were present). Also, in order to accommodate the fibula in articulation with the tibia, some fibula are slightly tilted in lateral view in order to position the anterior end within the cnemial crest but also contact the anterolateral crest and distal end of the fibula within the ascending processes of the tibia distal end (e.g., *Lirainosaurus astibiae*; Díez Díaz et al., 2013; Sanz et al., 1999).

Also, each element of the hind limb is separated by an articular cap much larger than in other known extant archosaurs (Bonnan et al., 2013; Holliday et al., 2010; Schwarz et al., 2007; Voegele et al., 2022). The femur, and the hind limb overall, may exhibit the best correlation between cartilage cap and bone morphology constrained by its role as support under most of the stress of the body mass (Bonnan et al., 2013; Voegele et al., 2022). Despite cartilaginous cap thickness is not constant and may vary among taxa, we set up our model hind limbs with a similar constant space between the femur and the zeugopod bones in all the sampled taxa in order to have a standardized initial point for comparisons and that there is no way to estimate the volume of the articular cap in the sampled specimens. The anatomical mounts include an additional space of 2% of the element length between stylopod and zeugopod articular surfaces in anatomical position (following Voegele et al., 2022).

### S2.2. Landmark-based 3D-GMM

#### S2.2.1. 3D Landmark sampling

The complete taxon hind limbs were used as a basis for a new sample of landmarks and semilandmarks, considering the new position of the different bone elements and the need of less landmark information for summarizing morphological changes in this study. Many of the sampled landmarks and semilandmark curves resemble the ones used in the analyses of the hind limb elements separated (Páramo, 2020; Páramo et al., 2022, 2020) or used during the Statistical Virtual Restoration phase previously described. However, the different focus of this study allows us to reduce the number of landmarks of our previous studies (e.g., no curve defined along the femoral or tibial midshaft, no landmarks in the proximal or distalmost surfaces of the femur or zeugopodial elements, etc; see Fig. S2.4). A total of 28 landmarks and 12 semilandmark curves were placed on the hind limb bones partly based on previous studies (Lefebvre et al., 2022; Páramo et al., 2020; see Main Text Table 2 and Fig. 2; data available in Appendix S1) using the same IDAV Landmark Editor® software (Wiley et al., 2005) The semilandmarks were then slid in R using package *Morpho* (Schlager, 2017) following Gunz et al., (2005). To remove size differences, spatial position and the differences of the 3D mesh orientation, the resulting landmark and semilandmark configurations were superimposed via Generalized Procrustes Analysis (GPA) using the “procSym” function in *Morpho* package. Morphological variance was analysed with Principal Component Analysis (PCA; results in Main Text Table 3) saving the expected number of Principal Components (PCs) which summarize a significant amount of variance after an Anderson Chi’s test (see Bonnan, 2007).

| 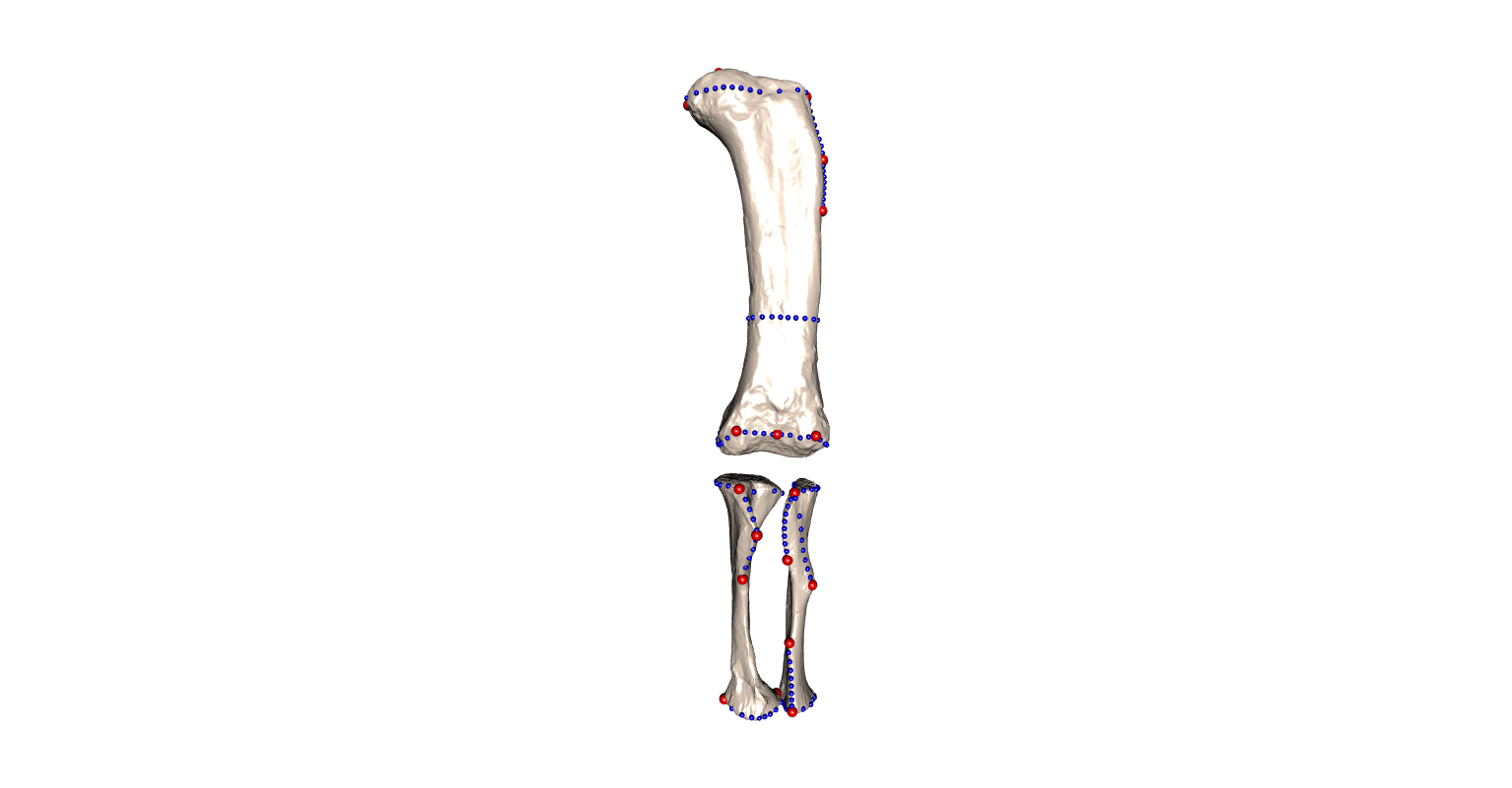 | 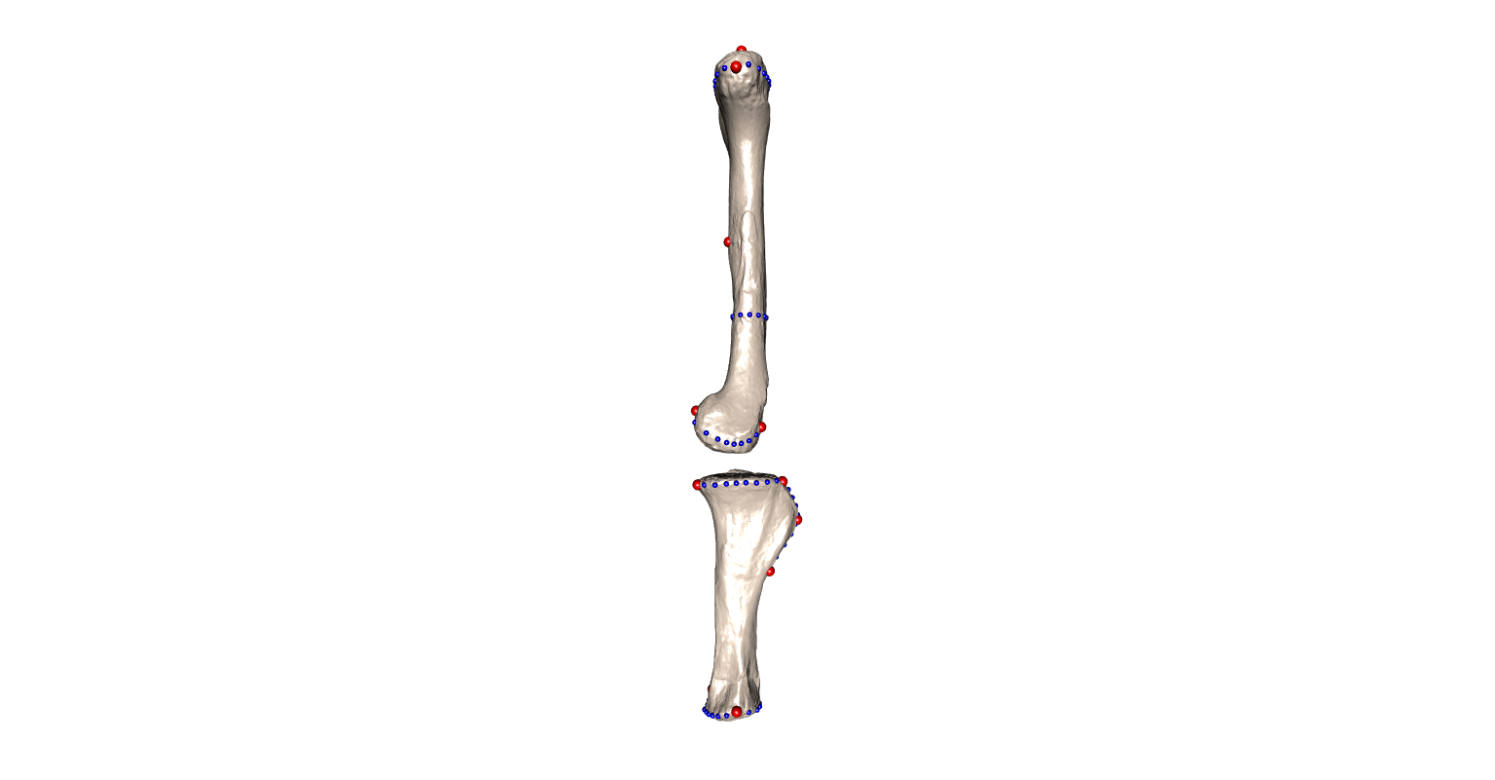 | 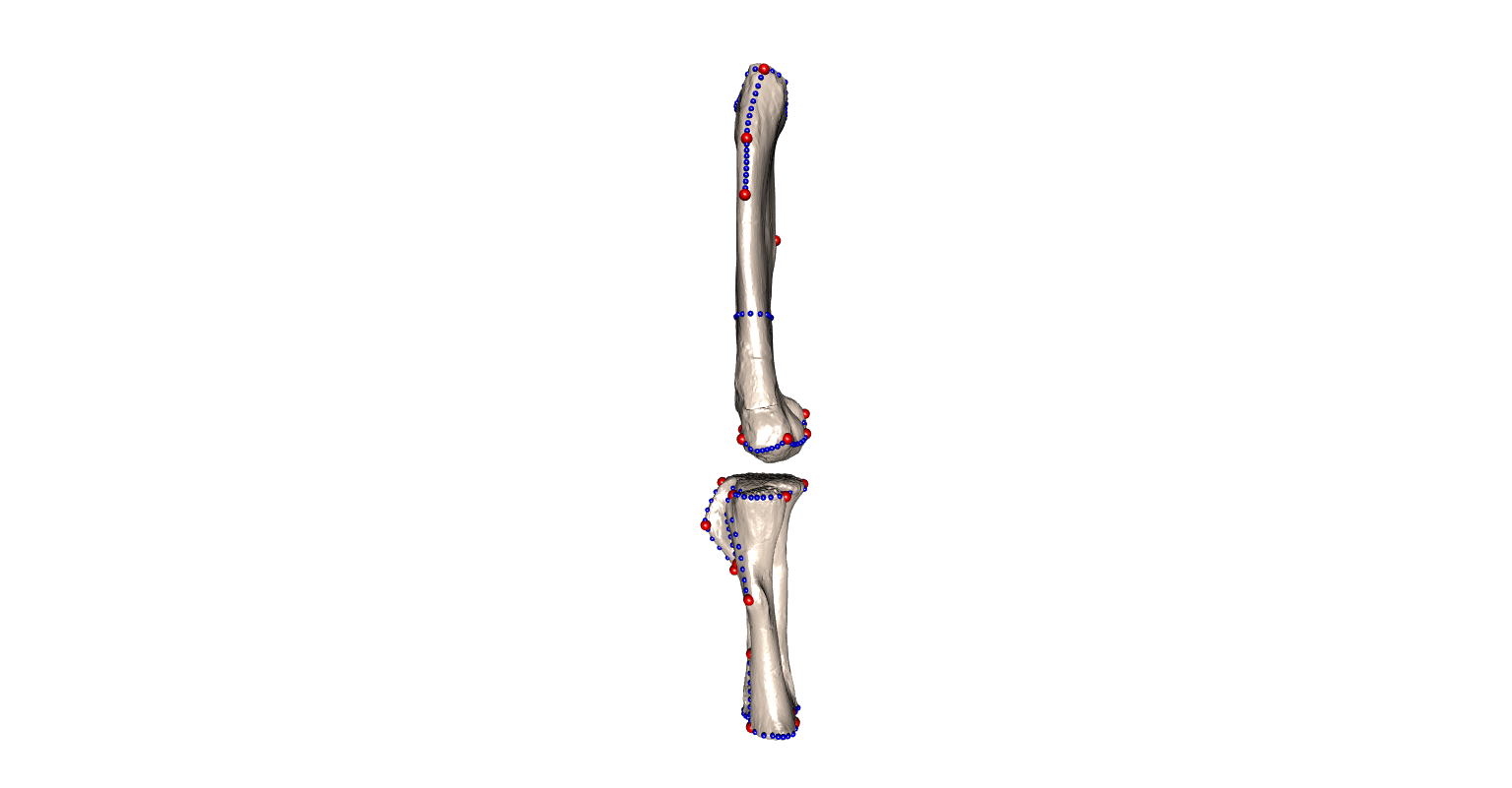 | 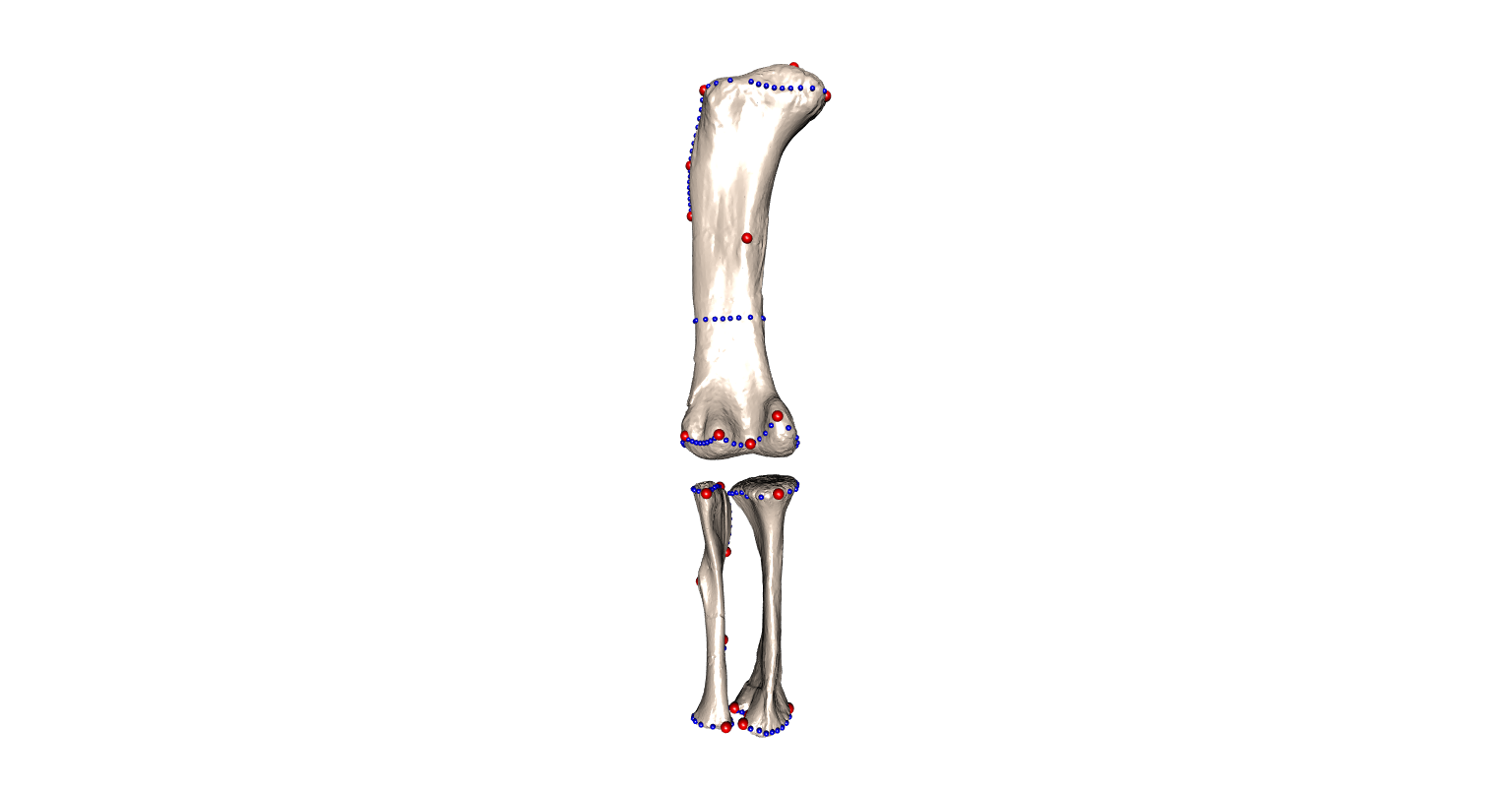 |
| --- | --- | --- | --- |
| Fig. S2.4. Landmark and semilandmark curves used in this study for the entire hind limb. Anterior, medial, lateral and posterior view. | | | |

#### S2.2.2. Summary Phylogenetic Supertree

The analysis of the macronarian hind limb morphology evolution requires a stable phylogenetic tree topology. However, the deeply-branched macronarian systematics and phylogenetics are still debated (e.g., Díez Díaz et al., 2021a; González Riga et al., 2018; Gorscak et al., 2023; Mannion et al., 2019, 2017; Sallam et al., 2018). We estimated a consensus tree topology using the matrix-representation parsimony supertree methodology (MRP; Bininda-Emonds, 2004) with the *phangorn* package (Schliep et al., 2017; Schliep, 2011). For supertree construction we compiled several of the more recent phylogenetic hypotheses that includes all the available sampled taxa (i.e., Carballido et al., 2017; Csiki et al., 2010; Díez Díaz et al., 2018; González Riga et al., 2018; Mannion et al., 2019; Mocho et al., 2019; Fig. S2.5). The resulting supertree (Fig. S2.6) had their tips trimmed to our current macronarian sample. The tree topology was also time-calibrated using the taxon oldest and youngest appearance derived from the bibliography (see cites in age dataset, Appendix S1; see resulting supertree in Fig. S.2.7). The resulting phylogenetic relationships were projected onto the shape-PCA to visualize the phylomorphospaces.

| 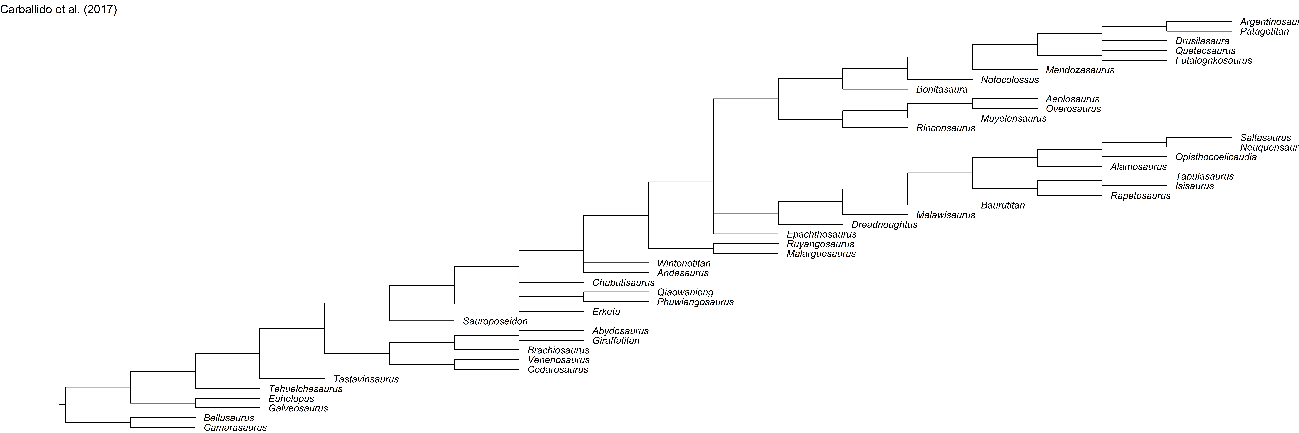 | 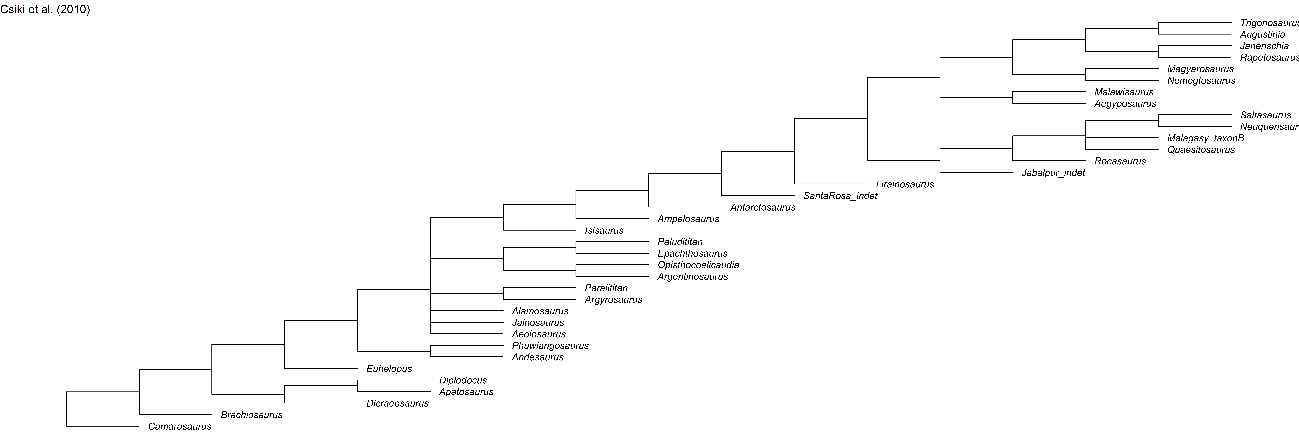 |
| --- | --- |
| Fig. S2.5. Phylogenetic trees used for MRP supertree estimation. | |
| 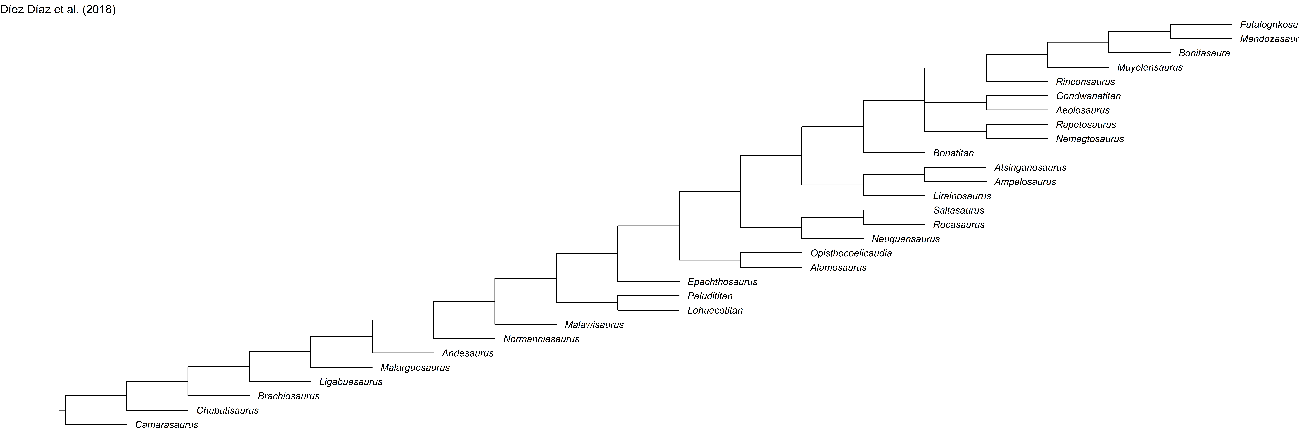 | 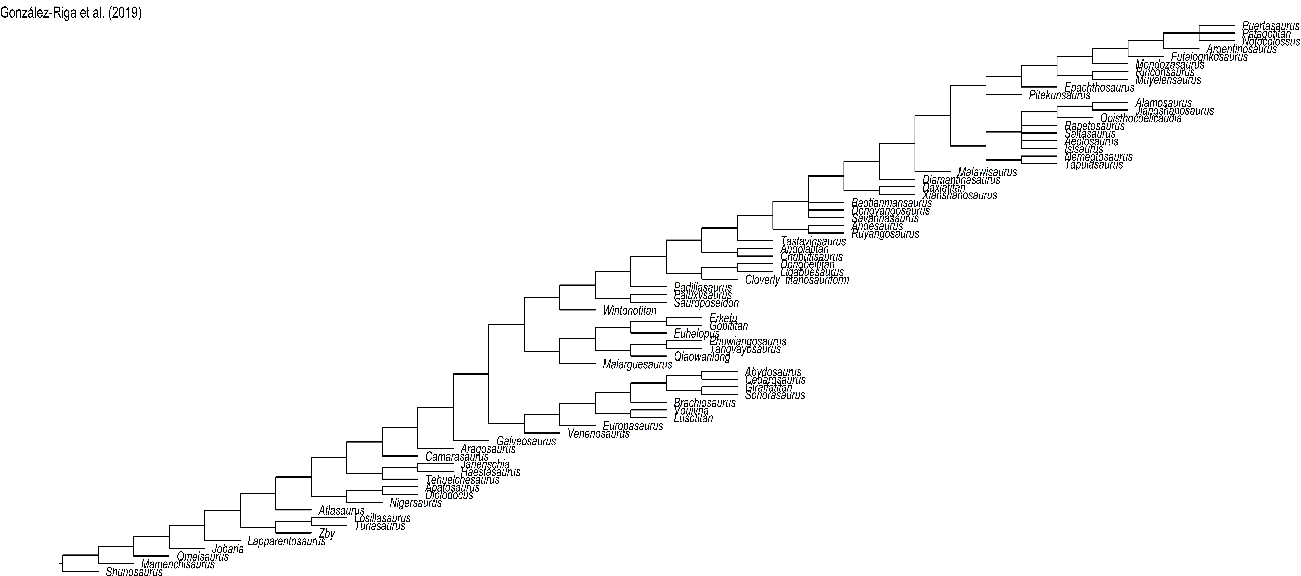 |
| Fig. S2.5. Phylogenetic trees used for MRP supertree estimation. (continues) | |
| 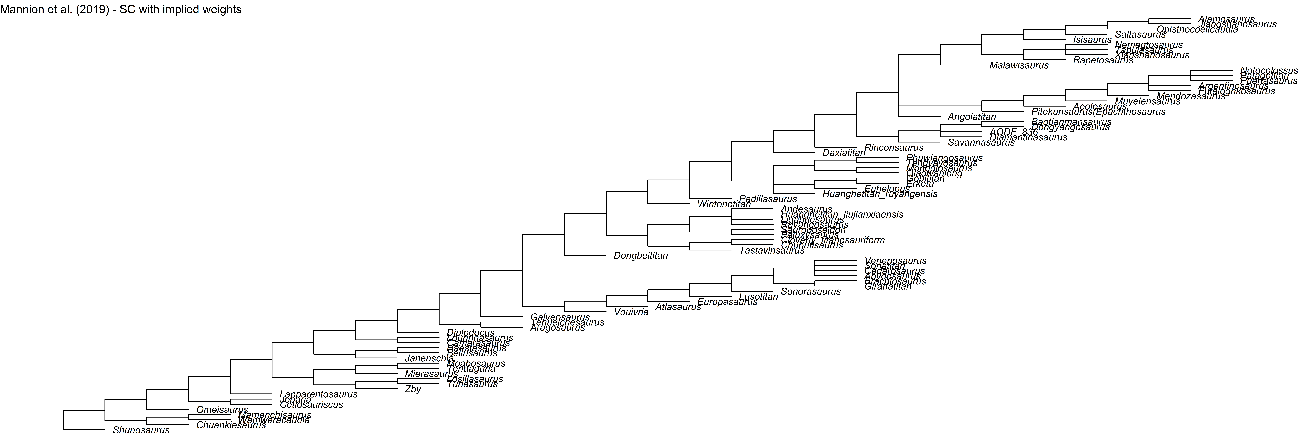 | 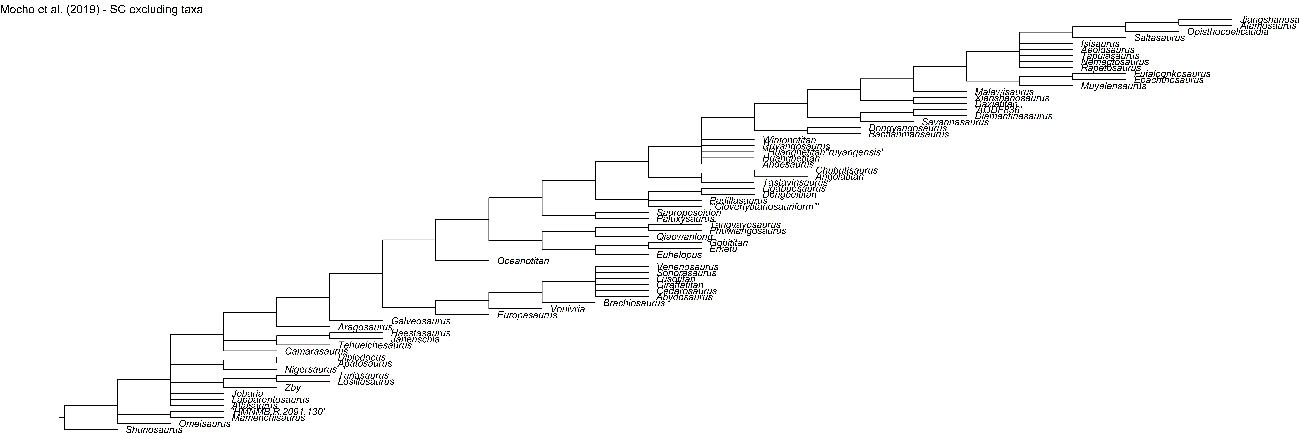 |
| Fig. S2.5. Phylogenetic trees used for MRP supertree estimation. (continues) | |

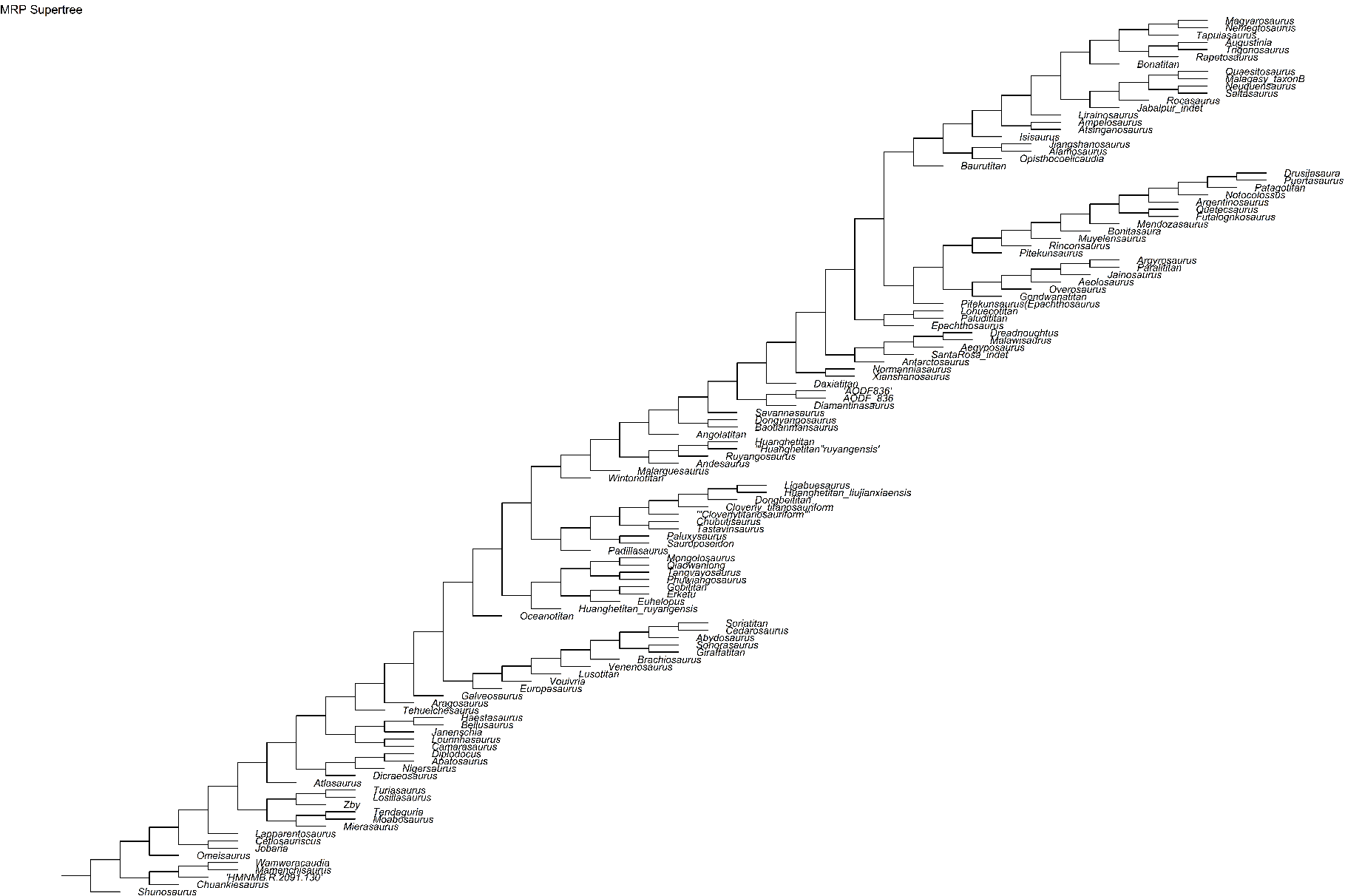

Fig. S2.6. Resulting supertree.

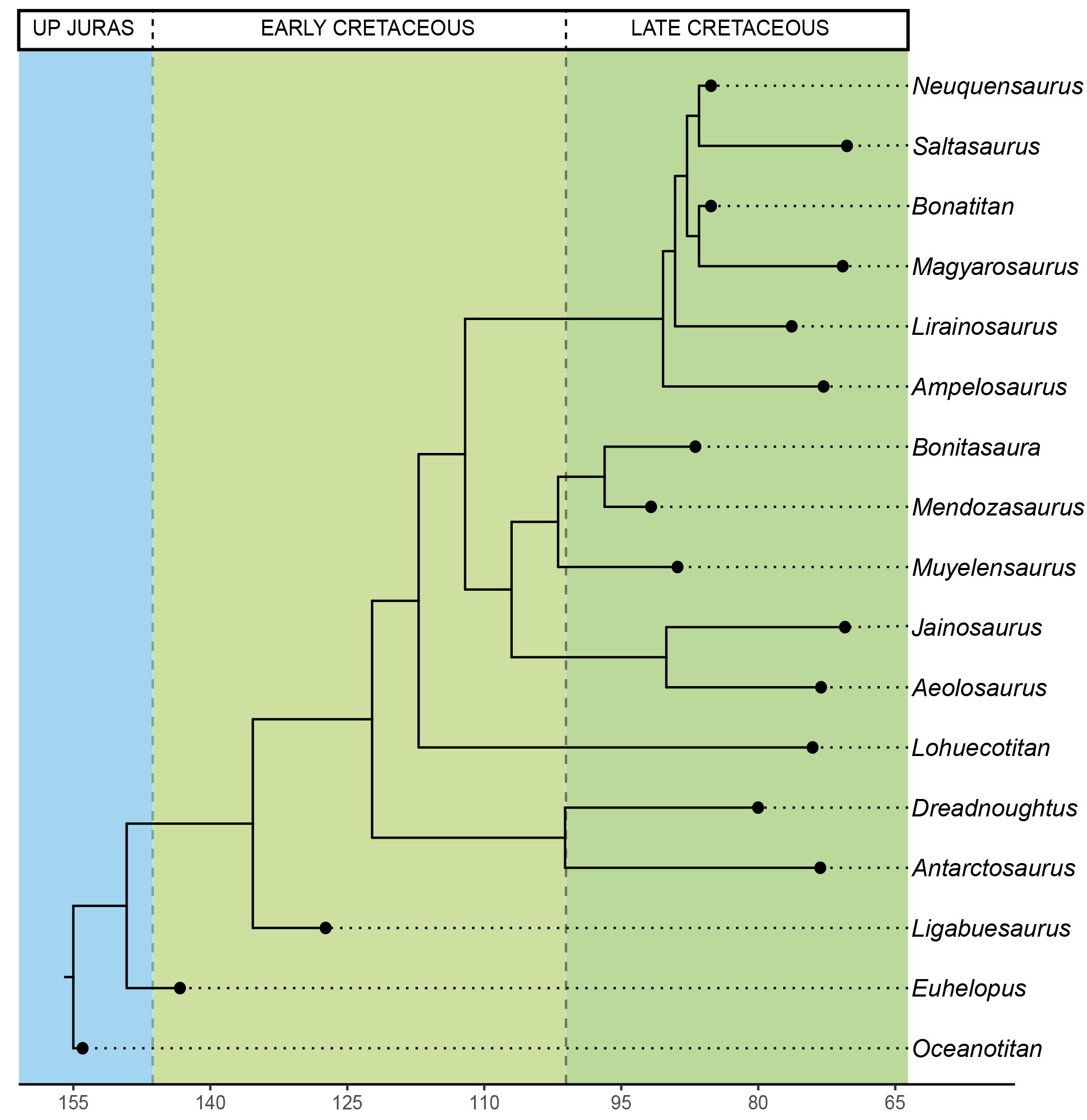

Fig. S2.7. Time calibrated supertree, tips limited to our current sample.

#### S2.2.3. Morphological convergence

In order to test if there is morphological convergence in a phylogenetic context, we conducted several analyses. We used the standard dissimilarity tests without use of phylogeny topology included in the statistic in order to discuss the possible morphological similarities. The testes included the shape variables and for a group label we included the most-inclusive ingroup following the published material for each taxon (e.g., *Oceanotitan dantasi* is considered Macronaria – as non-titanosauriform macronarian; *Saltasaurus loricatus* is considered Saltasauridae, and so on). The differences between the subclades were tested via Mann Whitney U’s test and Kruskal-Wallis non-parametric tests accounting for the uneven distribution of the group (sub-clade) samples.

We also conducted phylogenetic ANOVA using the time-calibrated supertree topology with the *phytools* R package (Revell, 2012).

#### S2.2.4. Body size proxies

The hind limb size as a proxy of sauropod body size or mass is based on the assumption that, despite absence of data on the fore limb size or the sauropod humerus alone, there is a strong correlation between the hind limb elements and the body mass (Benson et al., 2014; Campione, 2017; Mazzetta et al., 2004). The body mass/size estimation could improve if we take in consideration data from the humerus as they’re quadrupedal animals and its hind limb alone may not be enough, however, estimation are still pretty accurate nonetheless (Campione and Evans, 2020). We proxied by a measurement of the landmark configuration specimen centroid size, which is the sum of squared distances between landmarks (Bookstein, 1991; Zelditch et al., 2012) and thus summarizes the size of the stylopodial and zeugopodial elements. Centroid size may be problematic as it is independent of “shape” at individual level but differences in the number of landmarks translate in increasing or decreasing centroid sizes of that particular configuration (centroid size equation in Zelditch et al., 2012). As we use landmark configurations and semilandmarks mostly bounded by type I and II landmarks (see Zelditch et al., 2012), the number of landmarks does not change across the sample. Therefore, the centroid size is comparable as size proxy between the specimens.

The sauropod body mass was proxied by the hind limb centroid size calculated as part of the GPA. Body mass can be estimated preferably by different sets of allometric equations using both humeral and femoral measurements (Mazzetta et al., 2004) or whole-body reconstruction volumetric estimations (Bates et al., 2016). However, it is not easy to apply both methods to our current sample as the entire skeletons were not digitized, nor publicly available and sometimes incomplete. We opted to assume that the hind limb, as it is the main sauropod body mass support, can be better used as a “conservative-minimal” proxy to its body mass, with larger hind limb corresponding to giant macronarian taxa. We tested for allometric relationships between the sauropod hind limb shape variables (PCs) and the centroid size as proxy to body mass via Reduced Major Axis (RMA) regression using *lmodel2* R package (Legendre, 2018).

To assess that centroid size as a proxy is not problematic, we contrasted our findings with alternative body mass estimations using linear measurements of specimen size and extant scaling approach (following Campione and Evans, 2020). We used morphometrics variables such as the humeral circumpherence, femoral length and circumpherence for estimating the evolutionary allometry according to femoral length as proxy of body mass/size (e.g., Bonnan, 2007), humeral-femoral circumpherence for the quadratic equation model from Campione & Evans (2012), and lastly the femoral circumpherence alone following Mazzeta et al. (2004). These equations are derived on the original study of Anderson et al. (1985) and includes slight modifications to tune the model but offer similar results (e.g., Campione, 2017; Campione and Evans, 2020; Mazzetta et al., 2004). The body mass was estimated via *MASSTIMATE* R package (Campione, 2015). The results can be accessed in Appendix S4 and S5.

#### S2.2.5. Ancestral Character Estimation and evolutionary changes

We used the time-calibrated supertree topology and our shape variables (PCs) and hind limb centroid size to generate the Pagel’s lambda (λ) with *phytools* package and test for phylogenetic signal of these traits. All the different hypotheses of statistical correlations, dissimilarity tests and phylogenetic signal tests were accepting as significant using an alpha level of 0.05 (a comprehensive report of the test results and a copy of the R code and packages used can be accessed in Appendix S4 and S5 respectively). Once identified those shape PCs that exhibit significant phylogenetical signal, as well as the log-transformed centroid size, we estimated their ancestral characters (ACEs) using maximum-likelihood and a simple Brownian evolutionary model similar to the one assumed for estimation of the Pagel’s lambda. We used the ACEs to observe trends in the evolution of titanosaur hind limb size and morphology (based on our shape-PCs). We evaluated the differences in body size (proxied by log-transformed hind limb centroid size ACEs) between terminal taxa and internal nodes and between internal nodes of Titanosauriformes, Somphospondyli, Titanosauria and Lithostrotia, as well as their subclades. The sum of changes, mean change, median change, positive, negative and the total amount of changes were evaluated for each of the above clades following Butler and Goswami (2008). We used a χ2 godness-of-fit test to evaluate whether body size increasing or decreasing, and shape-PCs occur at the same frequency (50%-50% null hypothesis) or have either a positive or negative tendency over titanosaurian evolution.
