## Appendix S3 revised for "Evolution of hind limb morphology of Titanosauriformes (Dinosauria, Sauropoda) analyzed via 3D Geometric Morphometrics reveals wide-gauge posture as an exaptation for gigantism"

Electronic Supplementary Material

Appendix S3 – Shape analyses

In this appendix there is a comprehensive figuration of all the shape analyses including those shape PCs explaining up to 75% of total variance. There is also a section of discussion of the effects of taphonomic deformation and virtual restoration method on the analyses.

### S3.1. Shape PCA and Phylomorphospaces

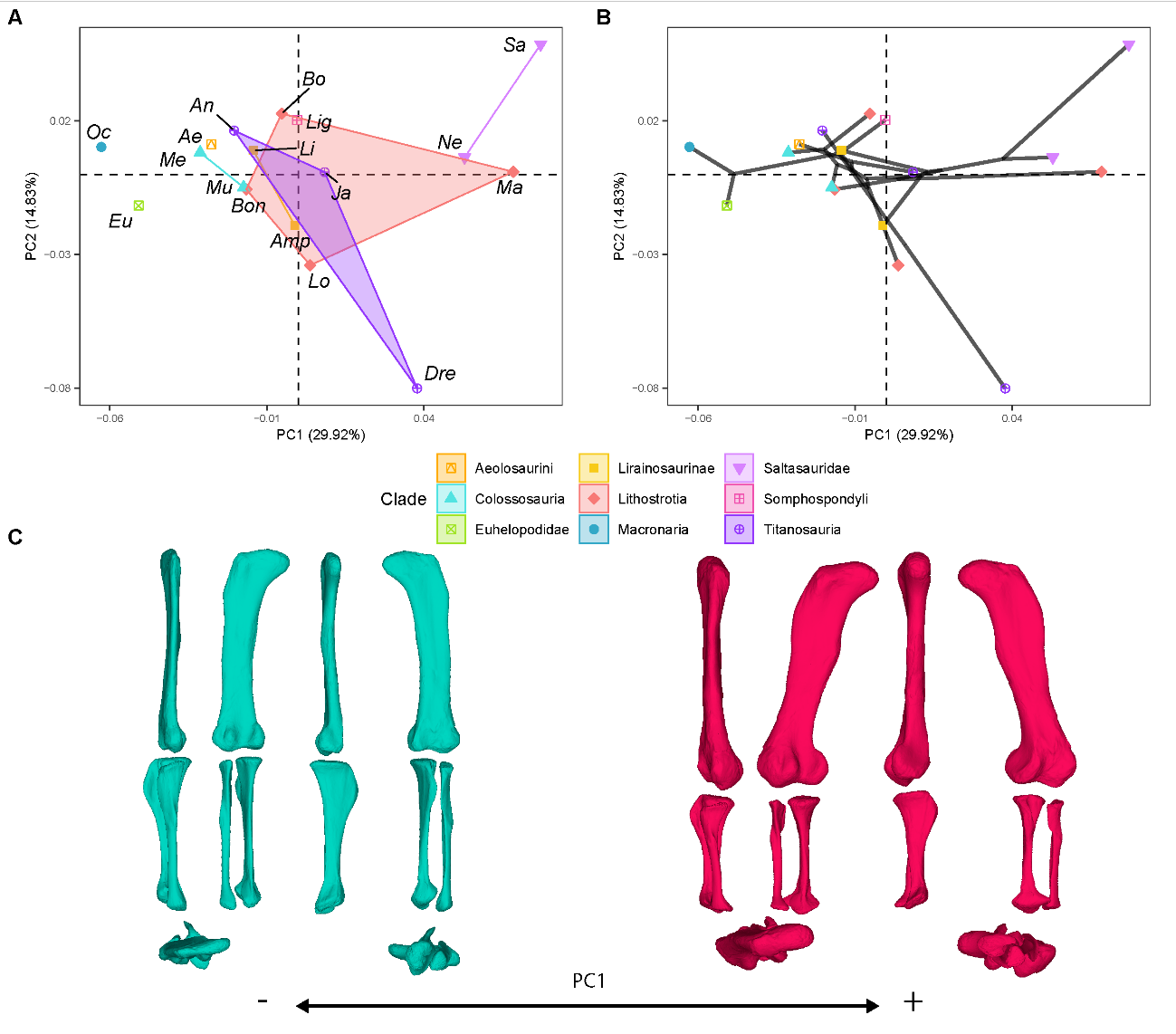

Figure S3.1.1. PCA results on the GPA aligned landmark and semilandmark curves of the hind limb. (a) PC1-PC2 biplot. (b) PC1-PC2 phylomorphospace with projected phylogenetic tree. (c) Representation of the shape change along PC1, blue are negative scores, red are positive scores. Percentage of variance of each PC in brackets under corresponding axis. *Ae* – *Aeolosaurus*, *Amp* – *Ampelosaurus*, *An* – *Antarctosaurus*, *Bo* – *Bonatitan*, *Bon* – *Bonitasaura*, *Dre* – *Dreadnoughuts*, *Eu* – *Euhelopus*, *Ja* – *Jainosaurus*, *Li* – *Lirainosaurus*, *Lig* – *Ligabuesaurus*, *Lo* – *Lohuecotitan*, *Ma* – *Magyarosaurus*, *Me* – *Mendozasaurus*, *Mu* – *Muyelensaurus*, *Ne* – *Neuquensaurus*, *Sa* – *Saltasaurus*.

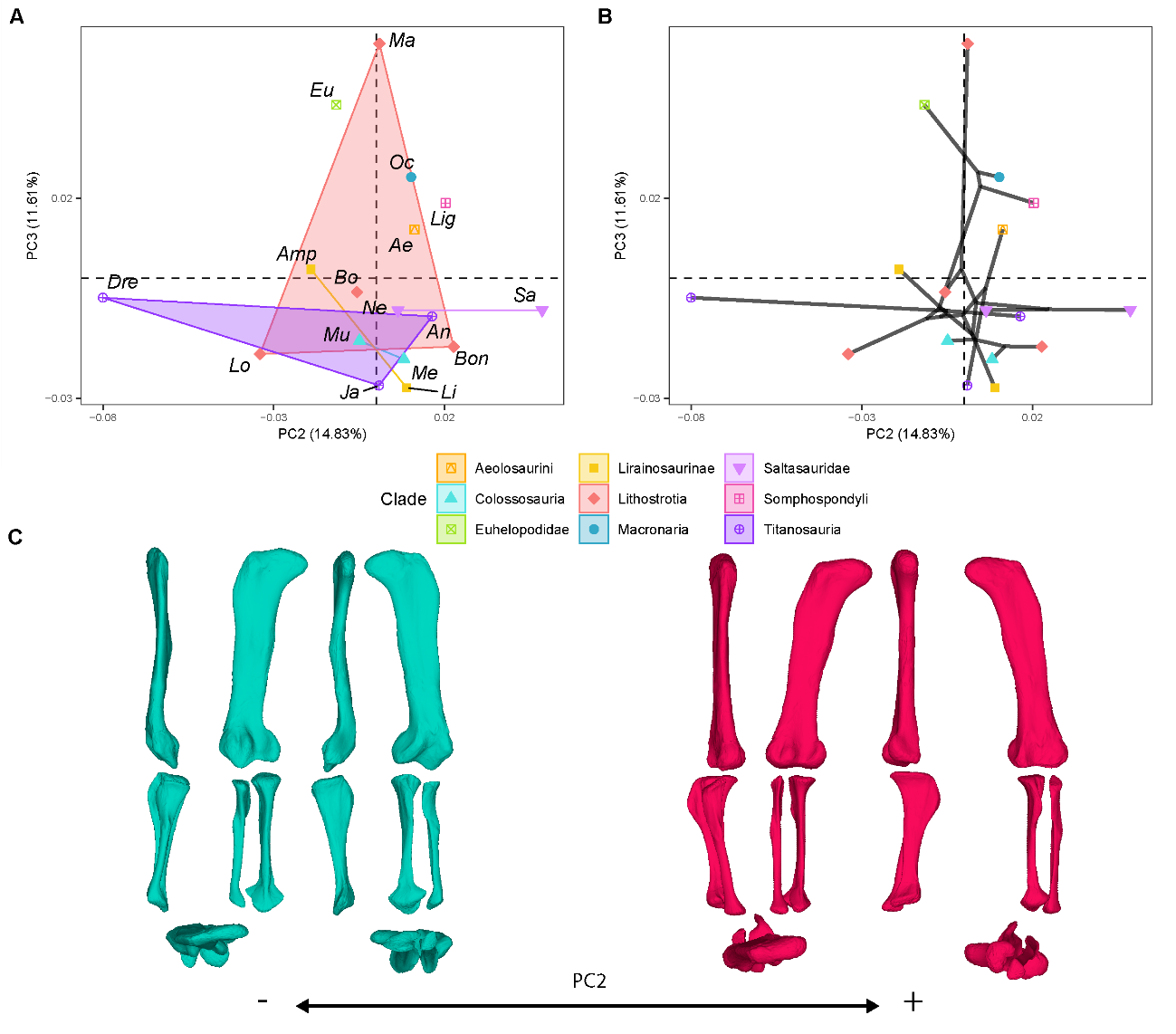

Figure S3.1.2. PCA results on the GPA aligned landmark and semilandmark curves of the hind limb. (a) PC2-PC3 biplot. (b) PC2-PC3 phylomorphospace with projected phylogenetic tree. (c) Representation of the shape change along PC2, blue are negative scores, red are positive scores. Percentage of variance of each PC in brackets under corresponding axis. *Ae* – *Aeolosaurus*, *Amp* – *Ampelosaurus*, *An* – *Antarctosaurus*, *Bo* – *Bonatitan*, *Bon* – *Bonitasaura*, *Dre* – *Dreadnoughuts*, *Eu* – *Euhelopus*, *Ja* – *Jainosaurus*, *Li* – *Lirainosaurus*, *Lig* – *Ligabuesaurus*, *Lo* – *Lohuecotitan*, *Ma* – *Magyarosaurus*, *Me* – *Mendozasaurus*, *Mu* – *Muyelensaurus*, *Ne* – *Neuquensaurus*, *Sa* – *Saltasaurus*.

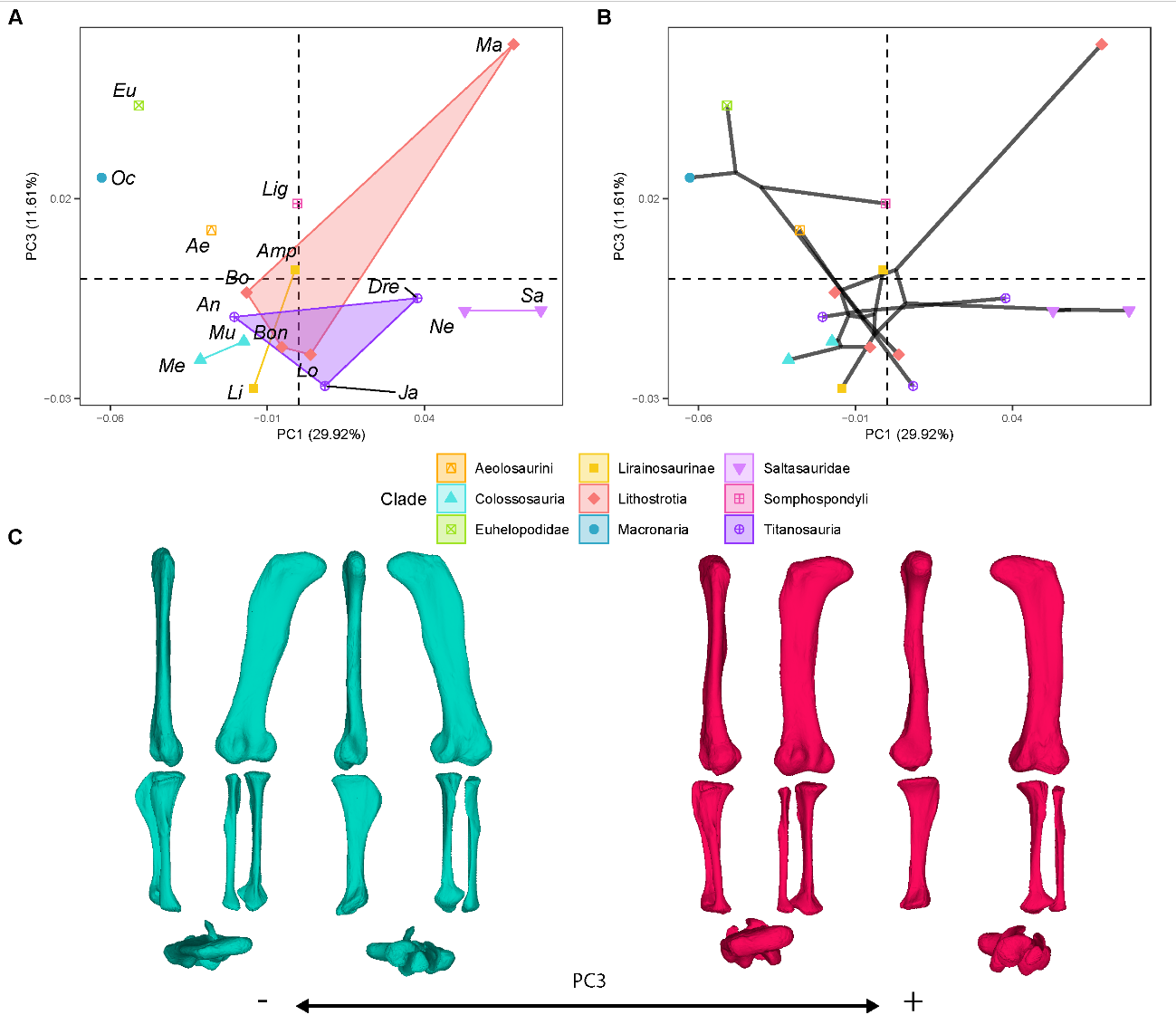

Figure S3.1.3. PCA results on the GPA aligned landmark and semilandmark curves of the hind limb. (a) PC1-PC3 biplot. (b) PC1-PC3 phylomorphospace with projected phylogenetic tree. (c) Representation of the shape change along PC3, blue are negative scores, red are positive scores. Percentage of variance of each PC in brackets under corresponding axis. *Ae* – *Aeolosaurus*, *Amp* – *Ampelosaurus*, *An* – *Antarctosaurus*, *Bo* – *Bonatitan*, *Bon* – *Bonitasaura*, *Dre* – *Dreadnoughuts*, *Eu* – *Euhelopus*, *Ja* – *Jainosaurus*, *Li* – *Lirainosaurus*, *Lig* – *Ligabuesaurus*, *Lo* – *Lohuecotitan*, *Ma* – *Magyarosaurus*, *Me* – *Mendozasaurus*, *Mu* – *Muyelensaurus*, *Ne* – *Neuquensaurus*, *Sa* – *Saltasaurus*.

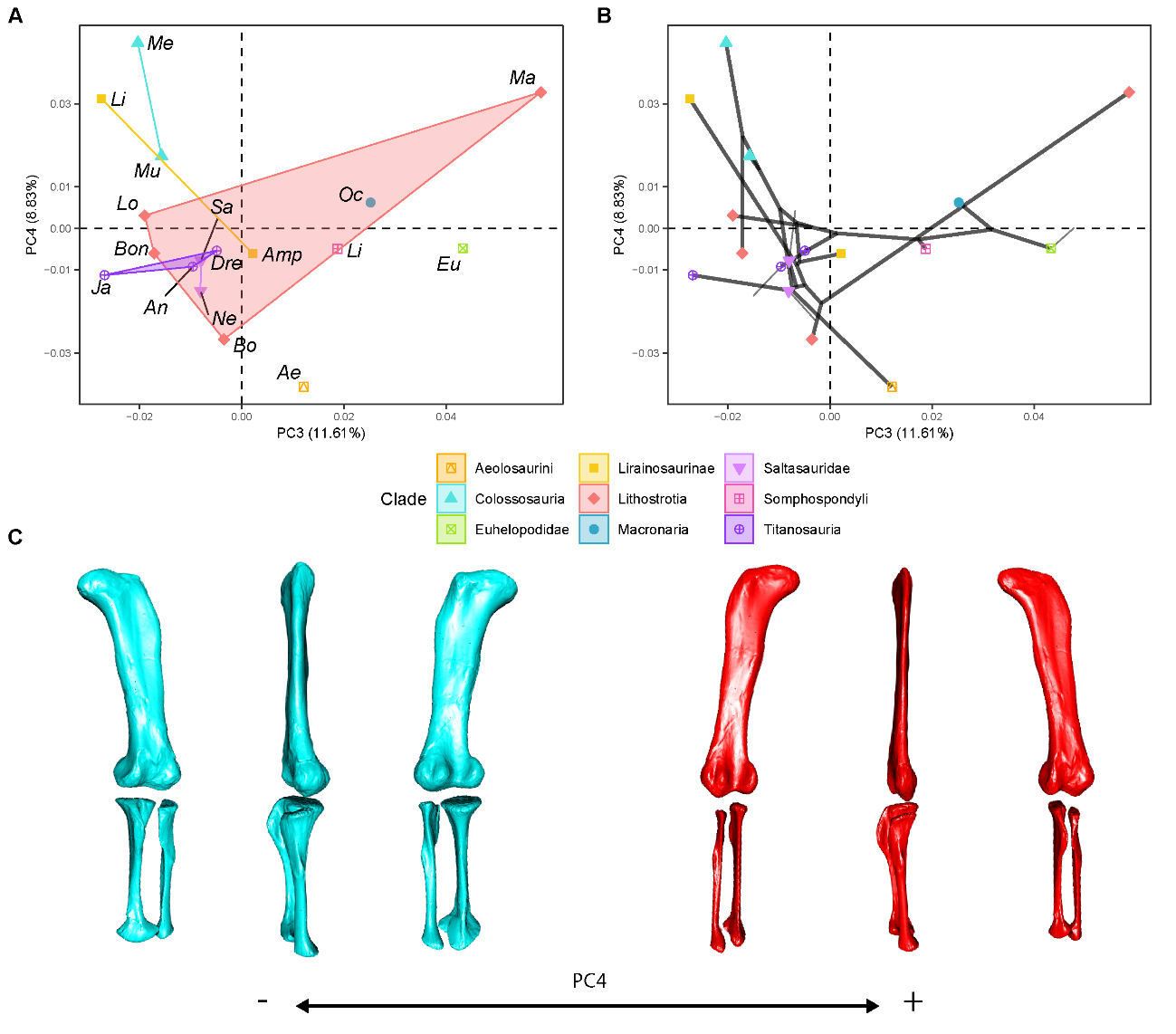

Figure S3.1.4. PCA results on the GPA aligned landmark and semilandmark curves of the hind limb. (a) PC3-PC4 biplot. (b) PC3-PC4 phylomorphospace with projected phylogenetic tree. (c) Representation of the shape change along PC4, blue are negative scores, red are positive scores. Percentage of variance of each PC in brackets under corresponding axis. *Ae* – *Aeolosaurus*, *Amp* – *Ampelosaurus*, *An* – *Antarctosaurus*, *Bo* – *Bonatitan*, *Bon* – *Bonitasaura*, *Dre* – *Dreadnoughuts*, *Eu* – *Euhelopus*, *Ja* – *Jainosaurus*, *Li* – *Lirainosaurus*, *Lig* – *Ligabuesaurus*, *Lo* – *Lohuecotitan*, *Ma* – *Magyarosaurus*, *Me* – *Mendozasaurus*, *Mu* – *Muyelensaurus*, *Ne* – *Neuquensaurus*, *Sa* – *Saltasaurus*.

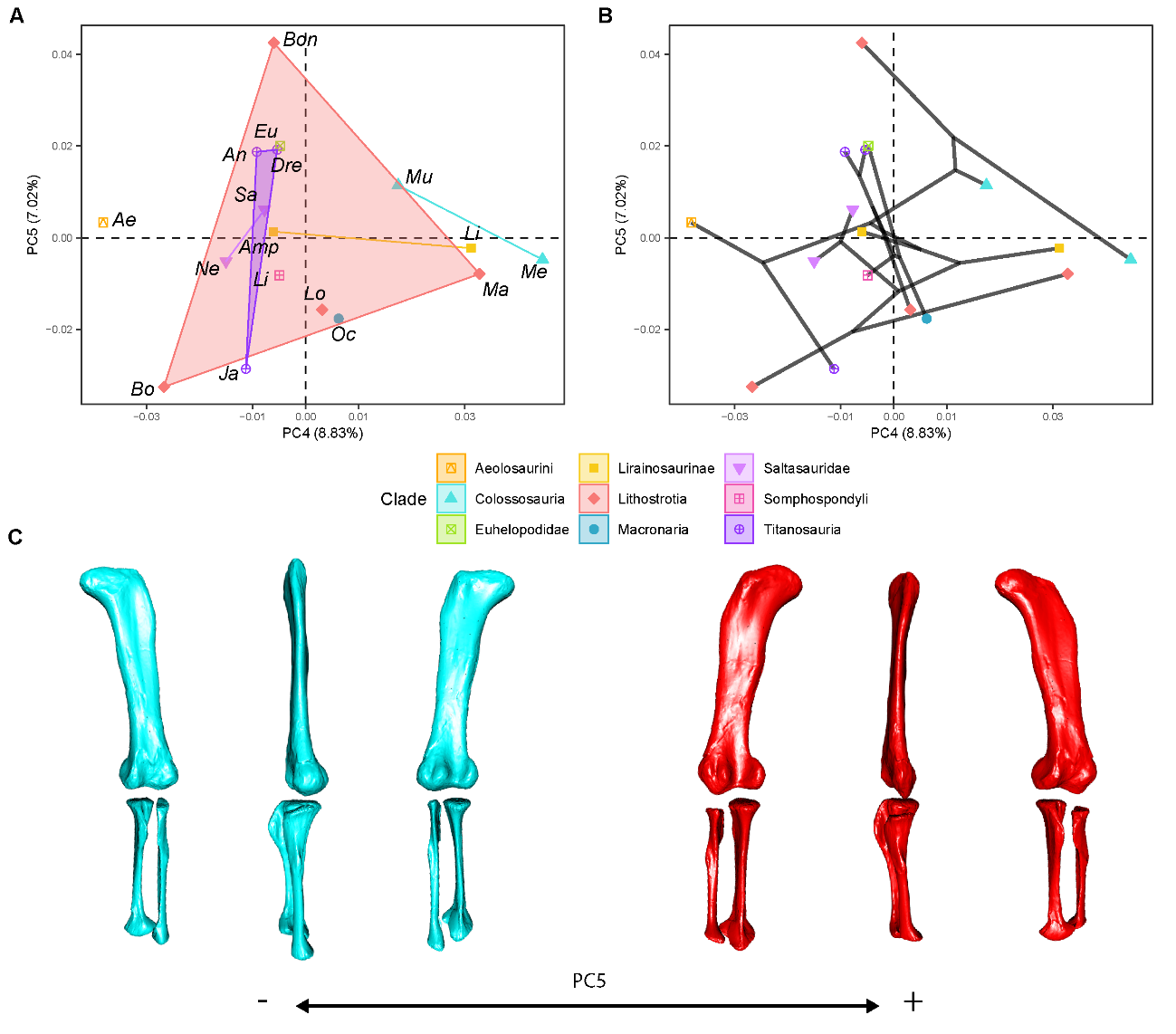

Figure S3.1.5. PCA results on the GPA aligned landmark and semilandmark curves of the hind limb. (a) PC4-PC5 biplot. (b) PC4-PC5 phylomorphospace with projected phylogenetic tree. (c) Representation of the shape change along PC5, blue are negative scores, red are positive scores. Percentage of variance of each PC in brackets under corresponding axis. *Ae* – *Aeolosaurus*, *Amp* – *Ampelosaurus*, *An* – *Antarctosaurus*, *Bo* – *Bonatitan*, *Bon* – *Bonitasaura*, *Dre* – *Dreadnoughuts*, *Eu* – *Euhelopus*, *Ja* – *Jainosaurus*, *Li* – *Lirainosaurus*, *Lig* – *Ligabuesaurus*, *Lo* – *Lohuecotitan*, *Ma* – *Magyarosaurus*, *Me* – *Mendozasaurus*, *Mu* – *Muyelensaurus*, *Ne* – *Neuquensaurus*, *Sa* – *Saltasaurus*.

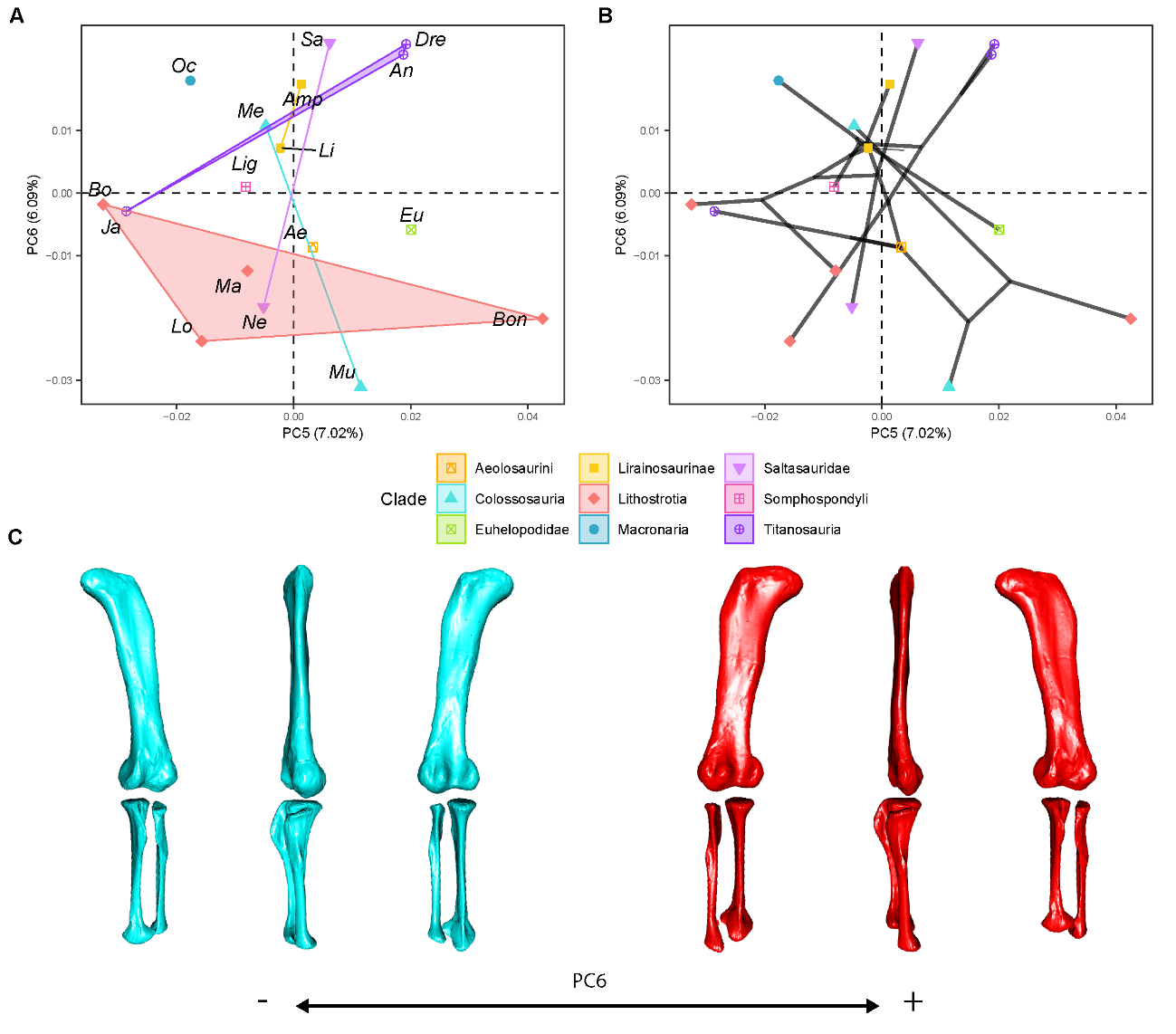

Figure S3.1.6. PCA results on the GPA aligned landmark and semilandmark curves of the hind limb. (a) PC5-PC6 biplot. (b) PC5-PC6 phylomorphospace with projected phylogenetic tree. (c) Representation of the shape change along PC6, blue are negative scores, red are positive scores. Percentage of variance of each PC in brackets under corresponding axis. *Ae* – *Aeolosaurus*, *Amp* – *Ampelosaurus*, *An* – *Antarctosaurus*, *Bo* – *Bonatitan*, *Bon* – *Bonitasaura*, *Dre* – *Dreadnoughuts*, *Eu* – *Euhelopus*, *Ja* – *Jainosaurus*, *Li* – *Lirainosaurus*, *Lig* – *Ligabuesaurus*, *Lo* – *Lohuecotitan*, *Ma* – *Magyarosaurus*, *Me* – *Mendozasaurus*, *Mu* – *Muyelensaurus*, *Ne* – *Neuquensaurus*, *Sa* – *Saltasaurus*.

### S3.2. RMA models with shape PCs – centroid size

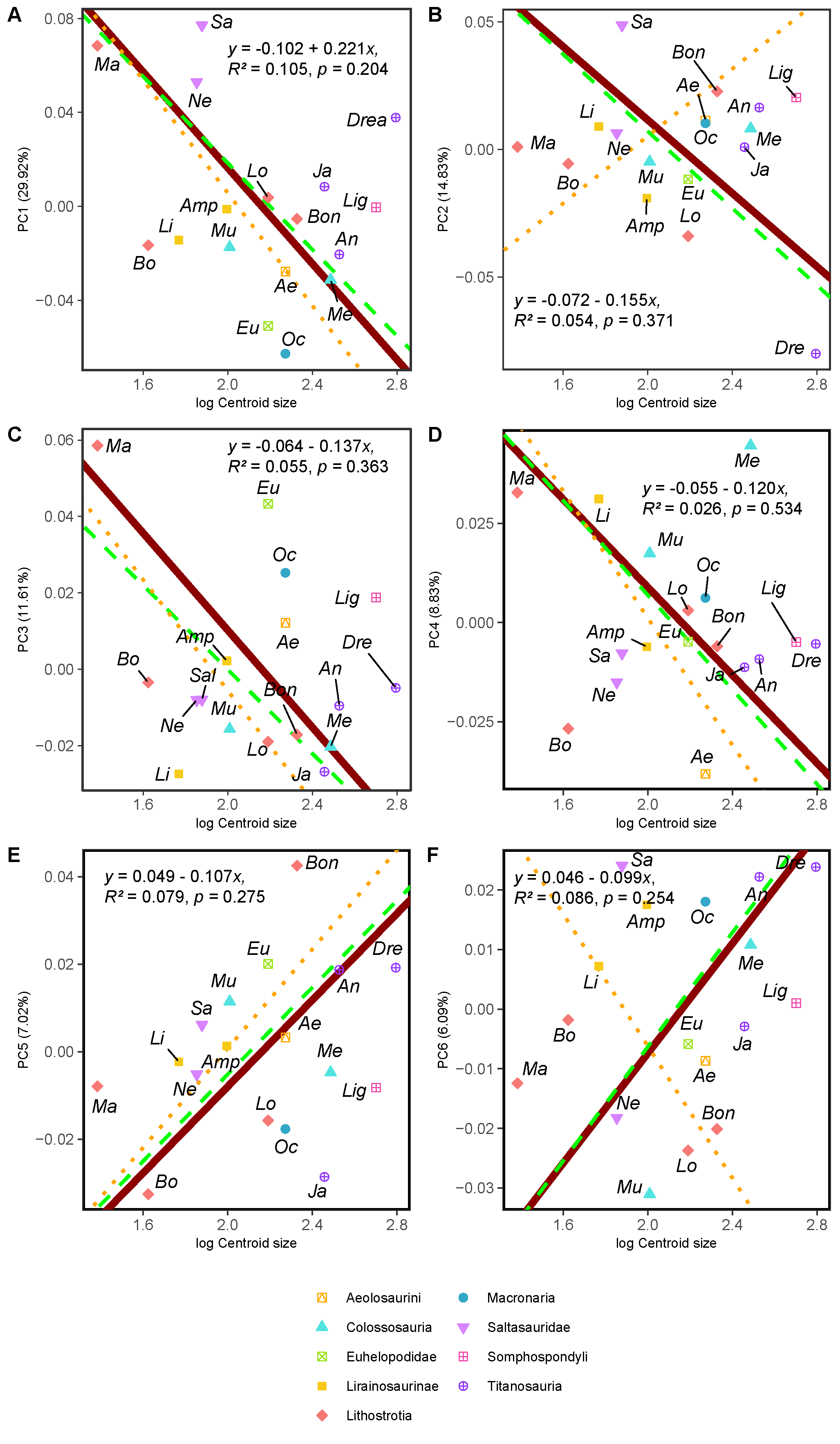

Fig. S3.2.1. RMA results of the first three shape PCs against the logarithm of the hind limb centroid size. (a) PC1 against log-Centroid size, all taxa RMA in dark red: intercept = 0.221, slope = -0.102, *r^2^* = 0.105, *p* = 0.204; Titanosauria only partial RMA in dashed green: intercept = 0.203, slope = -0.092, *r^2^* = 0.118, *p* = 0.229; Lithostrotia only partial RMA in dotted orange: intercept = 0.246, slope = -0.120, *r^2^* = 0.319, *p* = 0.07; (b) PC2 against log-Centroid size, all taxa RMA in dark red: intercept = 0.155, slope = -0.072, *r^2^* = 0.054, *p* = 0.371; Titanosauria only partial RMA in dashed green: intercept = 0.158, slope = -0.075, *r^2^* = 0.117, *p* = 0.232; Lithostrotia only partial RMA in dotted orange: intercept = -0.127, slope = 0.066, *r^2^* = 0, *p* = 0.952; (c) PC3 against log-Centroid size (Csize), all taxa RMA in dark red: intercept = 0.137, slope = -0.064, *r^2^* = 0.055, *p* = 0.363; Titanosauria only partial RMA in dashed green: intercept = 0.110, slope = -0.055, *r^2^* = 0.236, *p* = 0.078; Lithostrotia only partial RMA in dotted orange: intercept = 0.140, slope = -0.073, *r^2^* = 0.313, *p* = 0.074. (d) PC4 against log-Centroid size (Csize), all taxa RMA in dark red: intercept = 0.120, slope = -0.055, *r^2^* = 0.026, *p* = 0.534; Titanosauria only partial RMA in dashed green: intercept = 0.126, slope = -0.060, *r^2^* = 0.025, *p* = 0.592; Lithostrotia only partial RMA in dotted orange: intercept = 0.161, slope = -0.080, *r^2^* = 0.002, *p* = 0.903. (e) PC5 against log-Centroid size (Csize), all taxa RMA in dark red: intercept = -0.107, slope = 0.049, *r^2^* = 0.079, *p* = 0.275; Titanosauria only partial RMA in dashed green: intercept = -0.104, slope = -0.050, *r^2^* = 0.1506, *p* = 0.171; Lithostrotia only partial RMA in dotted orange: intercept = -0.113, slope = 0.057, *r^2^* = 0.204, *p* = 0.163. (f) PC6 against log-Centroid size (Csize), all taxa RMA in dark red: intercept = -0.100, slope = 0.046, *r^2^* = 0.086, *p* = 0.254; Titanosauria only partial RMA in dashed green: intercept = -0.102, slope = -0.048, *r^2^* = 0.089, *p* = 0.302; Lithostrotia only partial RMA in dotted orange: intercept = 0.105, slope = -0.055, *r^2^* = 0.003, *p* = 0.871. *Ae* – *Aeolosaurus*, *Amp* – *Ampelosaurus*, *An* – *Antarctosaurus*, *Bo* – *Bonatitan*, *Bon* – *Bonitasaura*, *Dre* – *Dreadnoughuts*, *Eu* – *Euhelopus*, *Ja* – *Jainosaurus*, *Li* – *Lirainosaurus*, *Lig* – *Ligabuesaurus*, *Lo* – *Lohuecotitan*, *Ma* – *Magyarosaurus*, *Me* – *Mendozasaurus*, *Mu* – *Muyelensaurus*, *Ne* – *Neuquensaurus*, Oc – *Oceanotitan*, *Sa* – *Saltasaurus*.

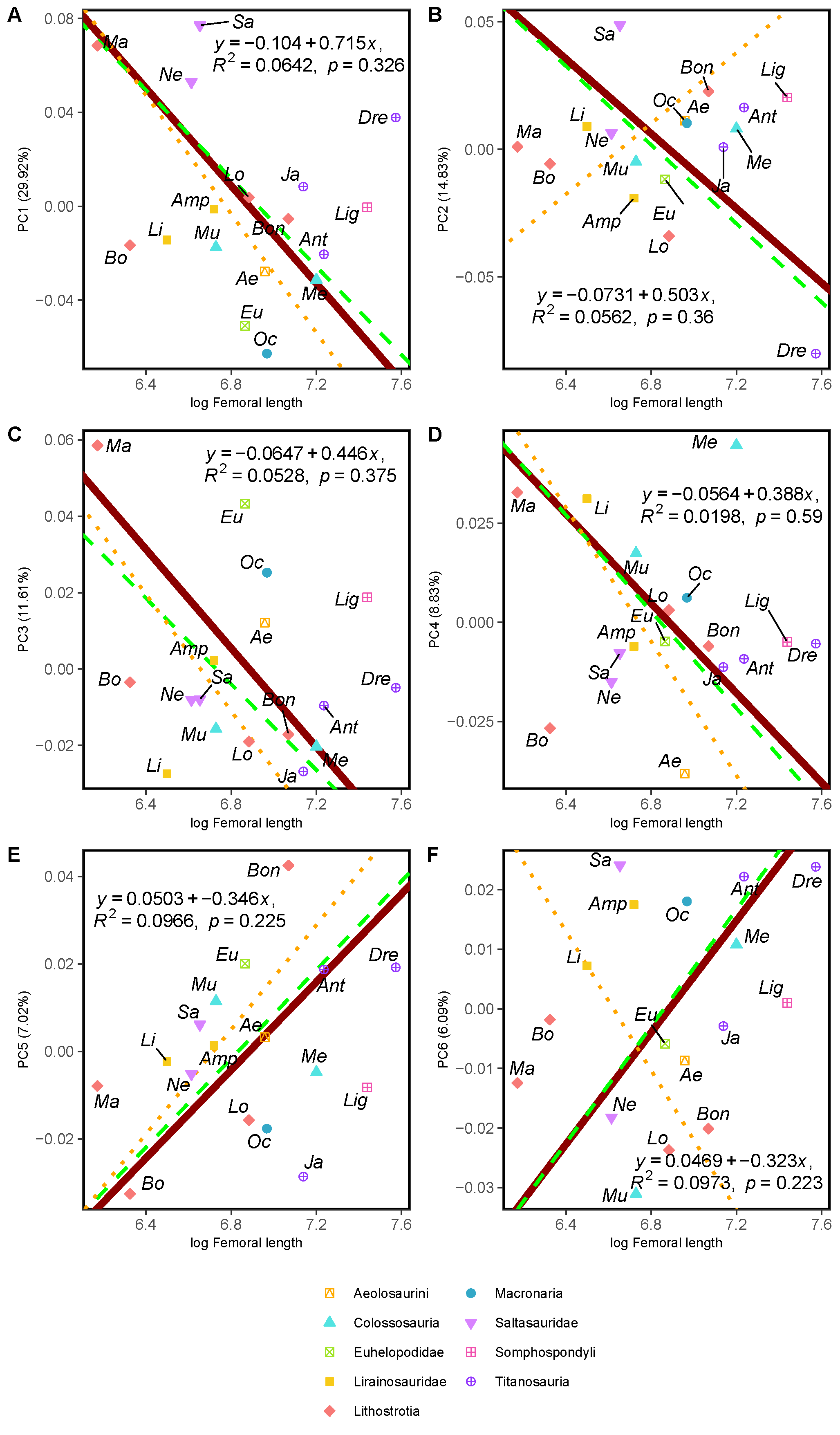

Fig. S3.2.3. RMA results of the first three shape PCs against the logarithm of the hind limb femoral length in mm. (a) PC1 against log-femoral length, all taxa RMA in dark red: intercept = 0.715, slope = -0.104, *r^2^* = 0.064, *p* = 0.326; Titanosauria only partial RMA in dashed green: intercept = 0.651, slope = -0.094, *r^2^* = 0.177, *p* = 0.337; Lithostrotia only partial RMA in dotted orange: intercept = 0.856, slope = -0.126, *r^2^* = 0.252, *p* = 0.115; (b) PC2 against log-Femoral length, all taxa RMA in dark red: intercept = 0.503, slope = -0.073, *r^2^* = 0.056, *p* = 0.36; Titanosauria only partial RMA in dashed green: intercept = 0.526, slope = -0.077, *r^2^* = 0.125, *p* = 0.215; Lithostrotia only partial RMA in dotted orange: intercept = -0.461, slope = 0.069, *r^2^* = 0.006, *p* = 0.823; (c) PC3 against log-Femoral length, all taxa RMA in dark red: intercept = 0.446, slope = -0.065, *r^2^* = 0.053, *p* = 0.375; Titanosauria only partial RMA in dashed green: intercept = 0.378, slope = -0.056, *r^2^* = 0.207, *p* = 0.102; Lithostrotia only partial RMA in dotted orange: intercept = 0.509, slope = -0.076, *r^2^* = 0.296, *p* = 0.084. (d) PC4 against log-Femoral length, all taxa RMA in dark red: intercept = 0.388, slope = -0.056, *r^2^* = 0.02, *p* = 0.59; Titanosauria only partial RMA in dashed green: intercept = 0.416, slope = -0.061, *r^2^* = 0.018, *p* = 0.647; Lithostrotia only partial RMA in dotted orange: intercept = 0.569, slope = -0.084, *r^2^* = 0.000, *p* = 0.972. (e) PC5 against log-Femoral length, all taxa RMA in dark red: intercept = -0.346, slope = 0.05, *r^2^* = 0.097, *p* = 0.225; Titanosauria only partial RMA in dashed green: intercept = -0.346, slope = 0.051, *r^2^* = 0.186, *p* = 0.124; Lithostrotia only partial RMA in dotted orange: intercept = -0.401, slope = 0.06, *r^2^* = 0.252, *p* = 0.116. (f) PC6 against log-Femoral length, all taxa RMA in dark red: intercept = -0.323, slope = 0.047, *r^2^* = 0.097, *p* = 0.223; Titanosauria only partial RMA in dashed green: intercept = -0.336, slope = -0.049, *r^2^* = 0.104, *p* = 0.261; Lithostrotia only partial RMA in dotted orange: intercept = 0.386, slope = -0.058, *r^2^* = 0.002, *p* = 0.903. *Ae* – *Aeolosaurus*, *Amp* – *Ampelosaurus*, *An* – *Antarctosaurus*, *Bo* – *Bonatitan*, *Bon* – *Bonitasaura*, *Dre* – *Dreadnoughuts*, *Eu* – *Euhelopus*, *Ja* – *Jainosaurus*, *Li* – *Lirainosaurus*, *Lig* – *Ligabuesaurus*, *Lo* – *Lohuecotitan*, *Ma* – *Magyarosaurus*, *Me* – *Mendozasaurus*, *Mu* – *Muyelensaurus*, *Ne* – *Neuquensaurus*, Oc – *Oceanotitan*, *Sa* – *Saltasaurus*.

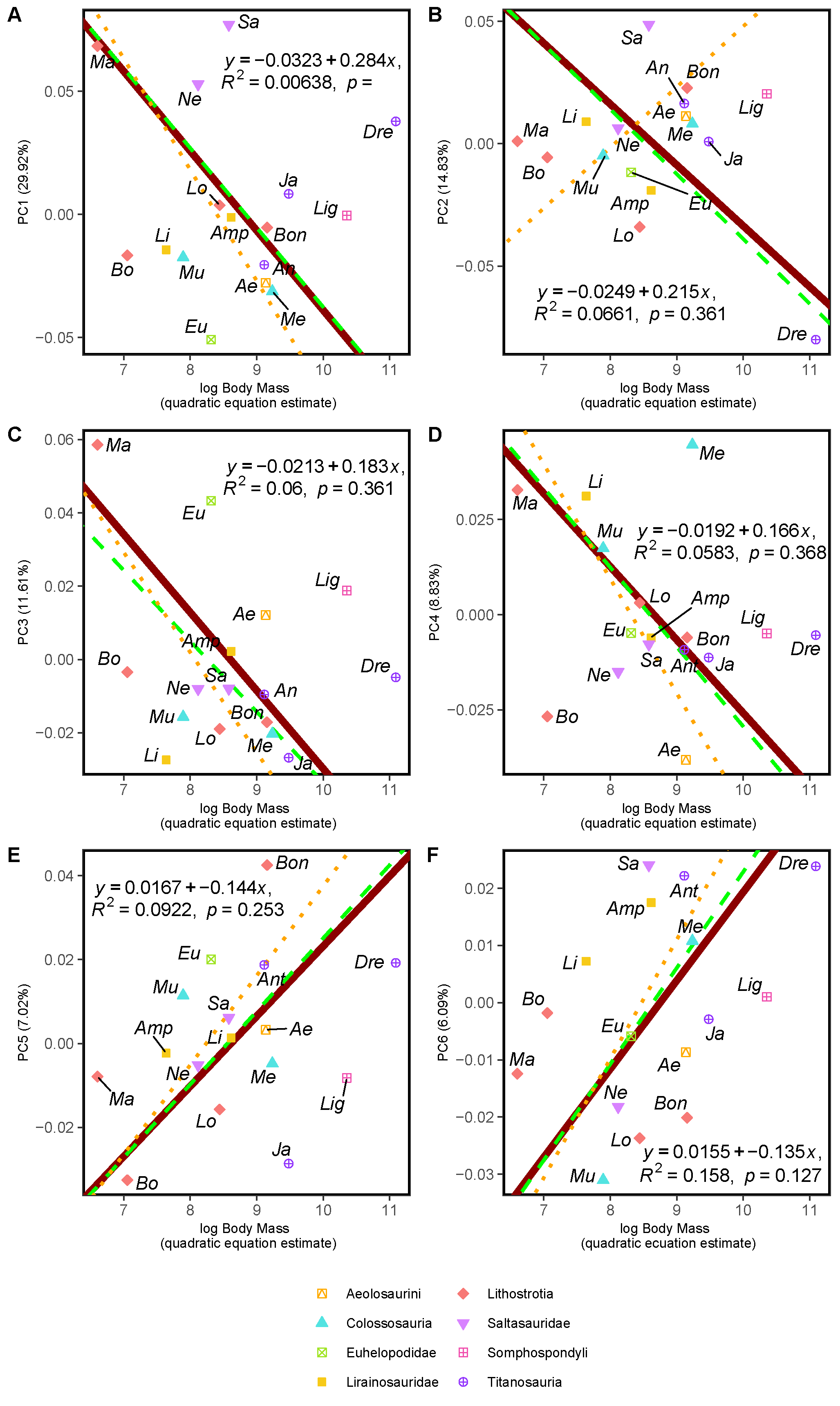

Fig. S3.2.3. RMA results of the first three shape PCs against the logarithm of the body mass in kilograms, estimated via quadratic equation method (Campione, 2017, 2015). (a) PC1 against log-body mass, all taxa RMA in dark red: intercept = 0.284, slope = -0.032, *r^2^* = 0.006, *p* = 0.769; Titanosauria only partial RMA in dashed green: intercept = 0.285, slope = -0.032, *r^2^* = 0.012, *p* = 0.712; Lithostrotia only partial RMA in dotted orange: intercept = 0.379, slope = -0.045, *r^2^* = 0.119, *p* = 0.299; (b) PC2 against log-Body mass, all taxa RMA in dark red: intercept = 0.215, slope = -0.025, *r^2^* = 0.066, *p* = 0.336; Titanosauria only partial RMA in dashed green: intercept = 0.226, slope = -0.026, *r^2^* = 0.119, *p* = 0.18; Lithostrotia only partial RMA in dotted orange: intercept = -0.2, slope = 0.025 *r^2^* = 0,053, *p* = 0.496; (c) PC3 against log-Body mass, all taxa RMA in dark red: intercept = 0.183, slope = -0.021, *r^2^* = 0.06, *p* = 0.361; Titanosauria only partial RMA in dashed green: intercept = 0.159, slope = -0.019, *r^2^* = 0.158, *p* = 0.159; Lithostrotia only partial RMA in dotted orange: intercept = 0.22, slope = -0.027, *r^2^* = 0.239, *p* = 0.127. (d) PC4 against log-Body mass, all taxa RMA in dark red: intercept = 0.166, slope = -0.019, *r^2^* = 0.058, *p* = 0.368; Titanosauria only partial RMA in dashed green: intercept = 0.18, slope = -0.021, *r^2^* = 0.059, *p* = 0.401; Lithostrotia only partial RMA in dotted orange: intercept = 0.25, slope = -0.03, *r^2^* = 0.038, *p* = 0.566. (e) PC5 against log-Body mass, all taxa RMA in dark red: intercept = -0.144, slope = 0.017, *r^2^* = 0.092, *p* = 0.253; Titanosauria only partial RMA in dashed green: intercept = -0.149, slope = 0.017, *r^2^* = 0.178, *p* = 0.133; Lithostrotia only partial RMA in dotted orange: intercept = -0.176, slope = 0.021, *r^2^* = 0.289, *p* = 0.088. (f) PC6 against log-Body mass, all taxa RMA in dark red: intercept = -0.100, slope = 0.046, *r^2^* = 0.086, *p* = 0.254; Titanosauria only partial RMA in dashed green: intercept = -0.145, slope = 0.017, *r^2^* = 0.173, *p* = 0.139; Lithostrotia only partial RMA in dotted orange: intercept = -0.176, slope = 0.021, *r^2^* = 0.025, *p* = 0.642. *Ae* – *Aeolosaurus*, *Amp* – *Ampelosaurus*, *An* – *Antarctosaurus*, *Bo* – *Bonatitan*, *Bon* – *Bonitasaura*, *Dre* – *Dreadnoughuts*, *Eu* – *Euhelopus*, *Ja* – *Jainosaurus*, *Li* – *Lirainosaurus*, *Lig* – *Ligabuesaurus*, *Lo* – *Lohuecotitan*, *Ma* – *Magyarosaurus*, *Me* – *Mendozasaurus*, *Mu* – *Muyelensaurus*, *Ne* – *Neuquensaurus*, Oc – *Oceanotitan*, *Sa* – *Saltasaurus*.

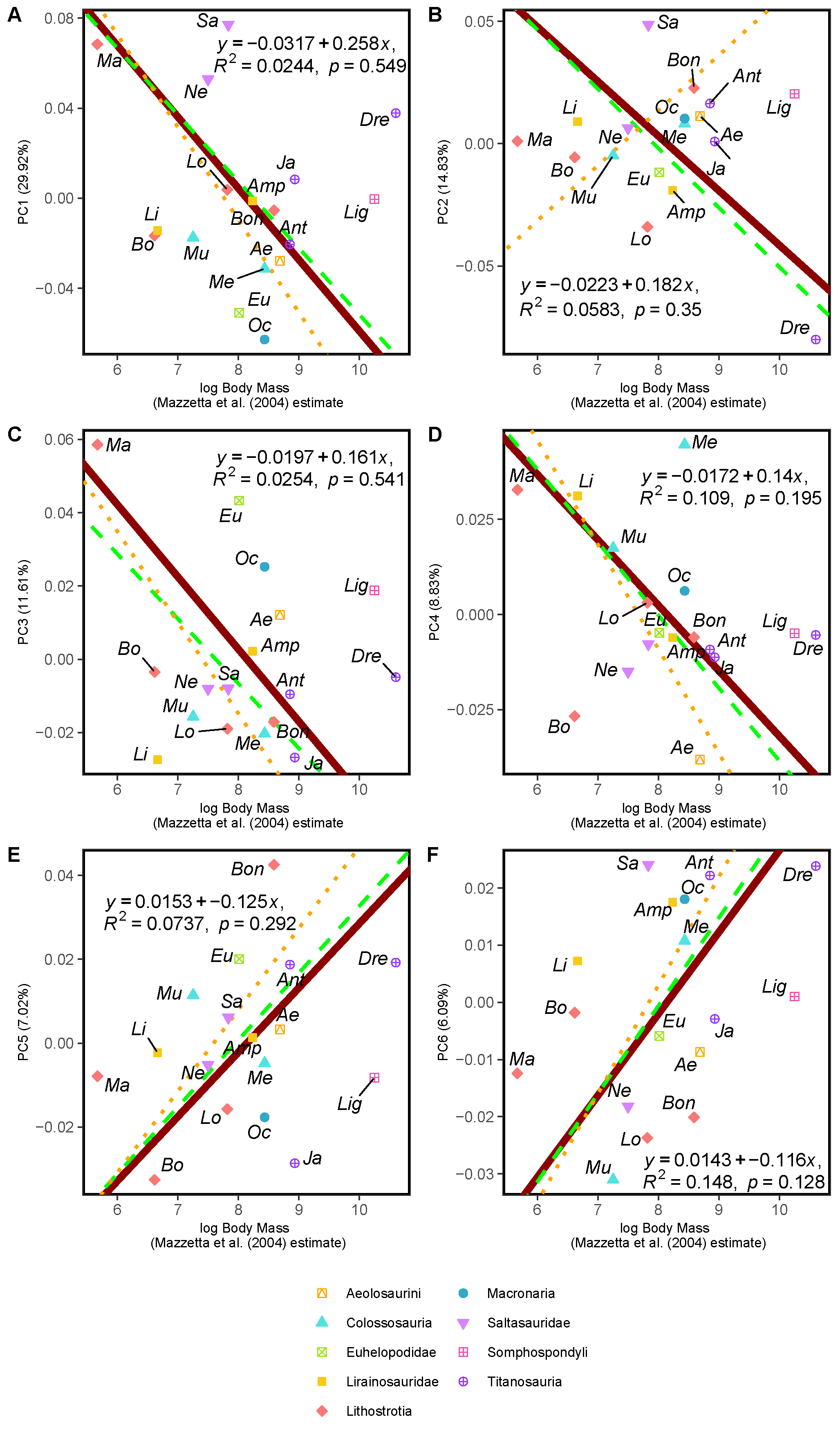

Fig. S3.2.4. RMA results of the first three shape PCs against the logarithm of the body mass in kilograms, estimated via hind limb multiple regression proposed by Mazzeta et al. (2004). (a) PC1 against log-body mass, all taxa RMA in dark red: intercept = 0.258, slope = -0.032, *r^2^* = 0.024, *p* = 0.549; Titanosauria only partial RMA in dashed green: intercept = 0.244, slope = -0.03, *r^2^* = 0.024, *p* = 0.597; Lithostrotia only partial RMA in dotted orange: intercept = 0.319, slope = -0.041, *r^2^* = 0.147, *p* = 0.245; (b) PC2 against log-Centroid size, all taxa RMA in dark red: intercept = 0.182, slope = -0.022, *r^2^* = 0.025, *p* = 0.35; Titanosauria only partial RMA in dashed green: intercept = 0.192, slope = -0.024, *r^2^* = 0.149, *p* = 0.173; Lithostrotia only partial RMA in dotted orange: intercept = -0.167, slope = 0.023, *r^2^* = 0.023, *p* = 0.657; (c) PC3 against log-Body mass, all taxa RMA in dark red: intercept = 0.161, slope = -0.02, *r^2^* = 0.025, *p* = 0.541; Titanosauria only partial RMA in dashed green: intercept = 0.135, slope = -0.018, *r^2^* = 0.148, *p* = 0.174; Lithostrotia only partial RMA in dotted orange: intercept = 0.184, slope = -0.025, *r^2^* = 0.215, *p* = 0.151. (d) PC4 against log-Body mass, all taxa RMA in dark red: intercept = 0.14, slope = -0.017, *r^2^* = 0.109, *p* = 0.195; Titanosauria only partial RMA in dashed green: intercept = 0.153, slope = -0.019, *r^2^* = 0.122, *p* = 0.221; Lithostrotia only partial RMA in dotted orange: intercept = 0.21, slope = -0.027, *r^2^* = 0.117, *p* = 0.302. (e) PC5 against log-Body mass, all taxa RMA in dark red: intercept = -0.125, slope = 0.015, *r^2^* = 0.074, *p* = 0.292; Titanosauria only partial RMA in dashed green: intercept = -0.127, slope = 0.016, *r^2^* = 0.169, *p* = 0.145; Lithostrotia only partial RMA in dotted orange: intercept = -0.147, slope = 0.019, *r^2^* = 0.242, *p* = 0.125. (f) PC6 against log-Body mass, all taxa RMA in dark red: intercept = -0.116, slope = 0.014, *r^2^* = 0.148, *p* = 0.128; Titanosauria only partial RMA in dashed green: intercept = -0.124, slope = 0.015, *r^2^* = 0.17, *p* = 0.142; Lithostrotia only partial RMA in dotted orange: intercept = -0.149, slope = -0.053, *r^2^* = 0.015, *p* = 0.716. *Ae* – *Aeolosaurus*, *Amp* – *Ampelosaurus*, *An* – *Antarctosaurus*, *Bo* – *Bonatitan*, *Bon* – *Bonitasaura*, *Dre* – *Dreadnoughuts*, *Eu* – *Euhelopus*, *Ja* – *Jainosaurus*, *Li* – *Lirainosaurus*, *Lig* – *Ligabuesaurus*, *Lo* – *Lohuecotitan*, *Ma* – *Magyarosaurus*, *Me* – *Mendozasaurus*, *Mu* – *Muyelensaurus*, *Ne* – *Neuquensaurus*, Oc – *Oceanotitan*, *Sa* – *Saltasaurus*.

### S3.3. Effects of Taphonomy and the Statistical Virtual Restoration on the shape analyses

In this section we will assess the effects of taphonomy and the virtual restoration and mean hind limb shape of the specimens, following the method of Lefebvre et al. (2020). Also, all the shape PCA results should be read as a composite between taphonomic deformation and shape variation between the analyzed specimens, or taphomorphospaces (Hedrick and Dodson, 2013). We estimated the mean shapes of the hind limb elements including only the best preserved specimens for each multi-specimen taxon previously analyzed in Páramo et al. (2020). As previous analyses suggest, effects of specimen deformation have low impact in the sample of reconstructed specimens (Páramo et al., 2020). However, some deformation in the input is unavoidable due lack of multiple specimens of the same taxon or general degree of deformation in the entire sample for the same taxon due preservation of a particular fossil site.

The reconstructed femur of *Oceanotitan dantasi* exhibit some degree of “sigmoid” curvature to posterior at the height of the fourth trochanter and then to anterior again at the proximal most area and the proximal end. This might have had some impact if we considered a larger set of landmarks and curves like in Páramo et al. (2020), but not in our reduced set of landmarks compared to previous studies. The virtual reconstruction of the holotype specimens shares some similarities with the other non-Titanosaurian, columnar hind limb of *Euhelopus zdanskyi* which also exhibit this morphology in the femur (PMU-234). In all the analyses this displacement of the femoral head does not seems to translate in an anomalous shape variation respective to the sample and the phylomorphospaces exhibit that it commonly falls in an area of the (tapho)morphospace dissimilar to the open hind limb of the titanosaurian sauropods due their “wide gauge” posture acquisition (e.g., S3.2). Maybe it would alter its position in PC5 which would have been in slightly higher positive values (Fig. S3.2.5-6), probably overlapping with saltasaurid specimens. The proximal part of the hind limb, especially the proximal end of the femur, may resemble the femora of some saltasaurid sauropods with the dorsally deflected femoral head, whereas the rest of the hind limb still exhibits the plesiomorphic non-Titanosauriformes macronarian morphology. It is noticeable that the fragmentary hind limb of *O. dantasi* also exhibits some lithostatic deformation, with slightly collapsed shafts that barely translate in noticeable effects in our analyses, but some degree of warping especially in the right tibia SHN-181-31. We cannot completely override that the proximal end of the tibia is lateromedially-narrow due lithostatic compaction (see Mocho et al., 2019b). However, the distal end of the tibia preserved its original lateromedial width and it would be uncommon that one end is affected and in the other one the lithostatic deformation is barely noticeable especially in absence of hints of complex tectonic deformations. The medial concavity of the shaft it is due deformation and fracturing and it does affect our results, as it conditions the articulation with the fibula and overall length of the tibia, but the effects are quite light. In the other hand, there are no hints of rotational deformation along the shaft axis that may alter the position of the distal end of the tibia, nor the placement of the fibular articulation with the tibia.

The holotype specimen of *Bonitasaura salgadoi* does not preserve the proximal end of the femur but also exhibits similar anterior bevelling of the lateral bulge (Fig. S3.3.1), which may produce a morphology closer to other lithostrotian sauropods like *Lohuecotitan pandafilandi*. When inspecting the holotype specimen of *Bonitasaura salgadoi* (MPCA-460) this anterior deflection of the lateral bulge can be a by-product of taphonomic deformation as well as our virtual restoration method or either a true anterior deflection as commented before (Fig. 3.3.1). However, none of the results exhibit insights of incorrect plotting of *B. salgadoi* as the anterior deflection of the lateral bulge has low impact alone. It is often separated even in PC4 and PC5 (S3.2.4-5) where it captures the anterior deflection of the lateral bulge and the area of the greater trochanter, despite exhibiting morphological similarities to *Ampelosaurus atacis* and *Lohuecotitan pandafilandi* due the virtual restoration process.

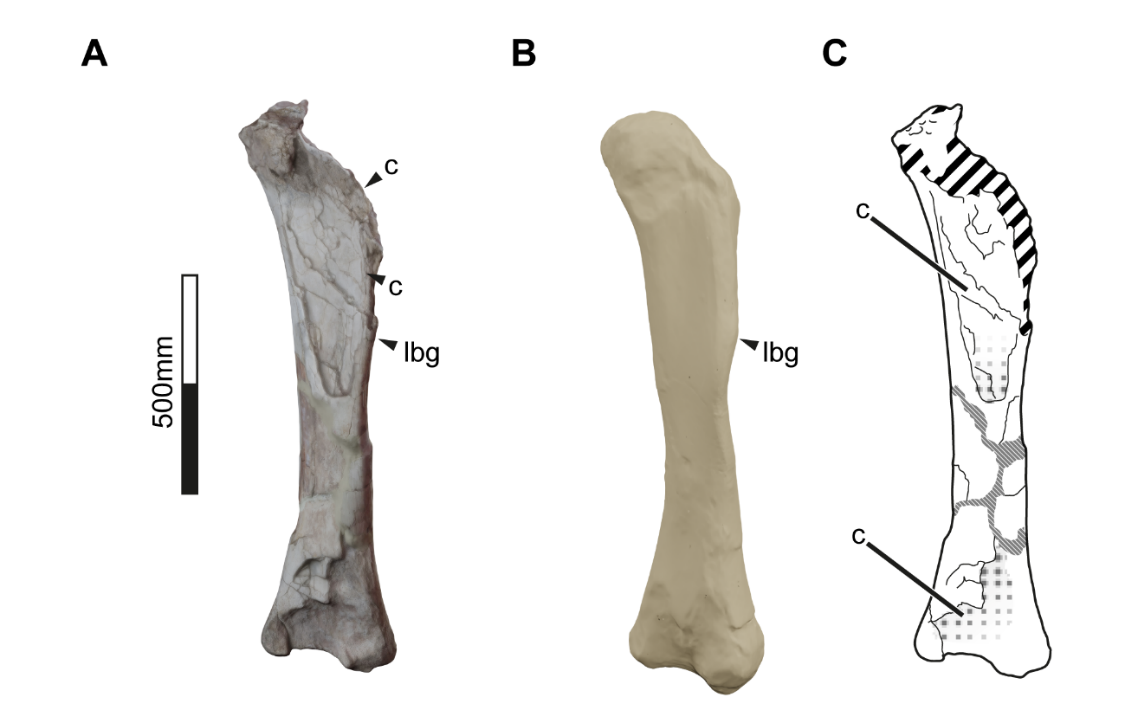

Fig. S3.3.1. 3D reconstruction of *Bonitasaura salgadoi* holotype (MPCA-460) left femur in anteolateral view. A. Original specimen 3D reconstruction. B. Complete femur as reconstructed by the statistical virtual restoration method. C. Diagram indicating the partial collapse of the anterior view of the shaft and the fragmentary proximal end. Abbreviations: c – collapse, lbg – lateral bulge. Discontinued lines/fill indicate fracture.

Despite possible bias in their overall position on the taphomorphospaces, our analyses seem to remain unaffected whether *B. salgadoi* lateral bulge is warped due the collapse of the proximal end or it truly does exhibit an anterior deflection similar to *E. zadanskyi* or some deeply-branched lithostrotians like *L. pandafilandi*. Small deviation in the position of *B. salgadoi* do no coerce large differences in the phylomorphospaces, neither it seems to increase the morphological variance in any of the main PCs.

The hind limb of *Dreadnoughtus schrani* exhibits some degree of lithostatic deformation especially the morphology of the femoral distal end (Ullmann and Lacovara, 2016). The femur distal end is medially rotated with an inward deflection of the condyles. As the authors suggest, this particular rotation apart of being characteristics does not correlated well with any biomechanical advantage and may not be biologically produced but rather taphonomic (Ullmann and Lacovara, 2016). The impact on our analyses is noticeable as *D. scharani* is recovered in a particularly different plotting compared to other deeply-branching titanosaurs due this medial rotation of the distal end of the femur and a slight distal bevelling of the condyles (PC2; Fig. S3.1.2). This extreme plotting is due the taphonomic deformation and cannot be interpreted as product of any evolutionary process. In fact, PC2 exhibit no significant phylogenetic signal and whether it is due the effects of taphonomic confounding factors or not is beyond the scopes of the current study.

Another factor observed in our sample is the lack of some information of the proximal and distal ends of some tibia. Especially the distal end which have been identified as one of the areas of the hind limb that contributes more to the sample variance (e.g., lateromedial expansion of the distal end in PC1, Fig. S3.1.1; anteroposterior orientation of the anterior ascending process on the distal condyle of the tibia in PC2, Fig. S3.1.2). Among the sampled specimens, *Aeolosaurus* sp. (specimen MPCA-27100-8) Has lost part of the lateral distal end of the tibia as well as part of the posterior ascending process of the distal condyle. Our reconstruction exhibits a conservative assumption of a rounded distal end with lateromedial width approximately the same as the anteroposterior width (Fig. S3.3.2). It’s uncertain if the lateral view of the distal end may be expanded as in *Elaltitan lilloi* (PVL-4628). But we can observe the morphology of the fossa between both ascending processes in medial view (Fig. S3.3.2) and the assumption of posterior orientation of the anterior ascending process of the virtual restoration fits the information preserved in the distal end of MPCA-27100-8.

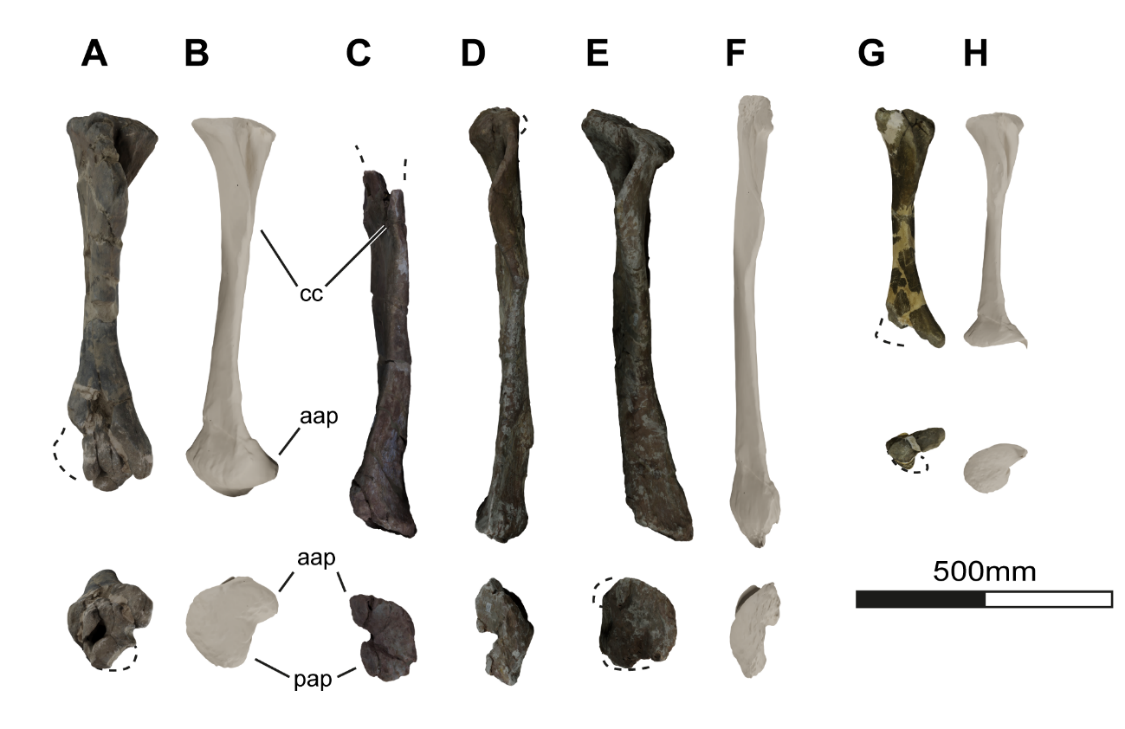

Fig. S3.3.2. Comparison between tibial specimens and their corresponding 3D virtual statistical rsestorations. *Aeolosaurus* sp.: A. Left tibia MPCA-27100-8 in anterior and distal views, B. Complete tibia as reconstructed by the statistical virtual restoration method in anterior and distal views. *Mendozasaurus neguyelap*: C. Right tibia IANIGLA-73-2 in anterior and distal views, D. Right tibia IANIGLA-73-3 in anterior and distal views, E. Right tibia IANIGLA-74-1 in anterior and distal views, F. Complete tibia as reconstructed by the statistical virtual restoration method in anterior and distal views. *Magyarosaurus* spp. G. Left tibia NHM-R3853 in anterior and distal views, H. Complete tibia as reconstructed by the statistical virtual restoration method in anterior and distal views. Abbreviations: aap – anterior ascending process, cc – cnemial crest, pap – posterior ascending process. Discontinued lines/fill indicate fracture.

The extremely gracile hind limb of *Mendozasaurus neguyelap* exhibits sights of lithostatic warping as longitudinal cracks and hints of shaft crushing, slight curvature of the midshaft, etc. The analyses recover *M. neguyelap* at some extremes of the robust-gracile hind limb spectrum (e.g., PC1-PC2; Fig. S3.1.1-2). However, several features like the rotation and morphology of the femoral and tibial proximal and distal ends seems slightly affected and thus translating in fewer taphonomic variance in the results of the PCs (e.g., PC1 as seen in Fig. S3.1.1). Such case is the rotation and shape of the distal end of the tibia, as seen in the specimen IANIGLA 73-2 which is completely preserved, but the incomplete specimens IANIGLA 73-3 and 74-1 also exhibits the lateromedially-narrow with rounded distal end, exhibiting an anterior and posterior ascending process slightly rotated toward posterior in distal view (Fig. S3.3.2). The lithostatic warping did not produce any sigmoidal component that might affect the proximodistal position or rotation of any feature. Among our results, PC2 and PC4 seems to be the most affected by this taphonomic deformation (see Fig. S.3.1.2 and S3.1.4), but these PCs exhibit no phylogenetic signal nonetheless (Table 6). It seems thought that some of our results that exhibit large variance in the morphology of the tibial proximal end may be altered by the reconstruction method (PC4-5, see Fig. S3.1.4-5), as the complete tibia IANIGLA-74-1 exhibits a slightly lateromedial-wider tibial proximal end compared with the reconstructed specimen. However, they sum up to a small portion of the total sample variance (PC4-PC5 sum up to 15.85% of the total variance) nonetheless. Similarly, *Muyelensaurus pecheni* exhibits a gracile and elongated hind limb and has also suffered from lithostatic compaction. However, all the specimens referred to *M. pecheni* from the same site exhibit some degree of crushing of the shaft, similar morphology of the proximal and distal ends, including a posterior rotation of the anterior and posterior ascending processes (e.g., MAU-PV-161 and 162). There is some room for lithostatic induced deformation as seen in the anterior fossa of the distal end, before the anterior ascending process, as seen in specimen MAU-PV-161 compared to the same fossa exhibited by specimens MAU-PV-162 and 430. But none of them suggest that either the proximal or distal ends rotations are affected as it may be the case of the distal end of the femur of *Dreadnoughtus schrani* (Ullmann and Lacovara, 2016).

The hind limb of *Magyarosaurus* is based on the classical collections from Nopcsa (1915). This material is sometimes fragmentary and scarce, with especial focus in the tibia as there is only one in the digitized sample (specimen NHM-R3853 referred to “*Magyarosaurus hungaricus*”). This also pose problems as *Magyarosaurus dacus* is an accepted taxon but *M. hungaricus*, *M. transsylvanicus* has been proposed as invalid taxa and preliminary referred to *M. dacus* (Upchurch et al., 2004). Among the dozen of individual recovered coming from different Maastrichtian fossil sites (Nopcsa, 1915), the re-study of the fossil remains deposited in the Romanian collections insight that some of the taxa referred to *Magyarosaurus* spp. might be chimeras and more than the original three morphotypes have been preliminary identified (Díez Díaz et al., 2023a). Assessment of the taxonomic distribution of titanosaurs from the Densuş-Ciula Formation (Maastrichtian) of Romania is still in debate (Díez Díaz et al., 2023a; Mocho et al., 2019a, 2024) and beyond the scopes of this study. For our study we reconstructed the hind limb of *Magyarosaurus* based on the combined sample of specimens previously referred to this taxon regardless of its current taxonomic status, thus this reconstruction may be biased by an unknown interspecific variability and known intraspecific variability like size differences between the recovered individuals (e.g., see S3.4. Ontogenetic status of the smaller specimens below).

Despite highly fragmentary, none of the studied specimens exhibit effects of lithostatic deformations imposed in the bone ends nor warping of the shafts. In general, *Magyarosaurus* spp. specimens are only affected by lack of preservation in several areas, up to the extreme of hindering the available sample. The tibial morphology have suffered the most of this lack of good preservation, as some other specimens were recovered by Nopcsa (1915), but lacks the proximal half (e.g., specimen NHM-R3859) or are only a fragment of the shaft (e.g., specimen NHM-R3849 referred to *M. dacus*). Lack of information in some distal or proximal ends in small percentages have not posed a problem before (Páramo et al., 2020) even if they represent missing information in the entire taxon (e.g., *Bonitasaura salgadoi* lack of the proximal end of the femur as discussed before). But large portions of missing information, especially in the same taxon has conduced our methods to errors and incapability of the algorithm to recover a stable solution, thus not returning any estimated landmark configuration. For this matter, we opted to use the left tibia NHM-R3853 previously referred to *M. hungaricus*. Robustness differences in this specimen are not appreciated or are not enough informative to be referred to a different taxon (APB pers. Obs.). Relative robustness does not differ enough to pose a different result in our analyses, as the position of *Magyarosaurus* plotted in the taphomorphospaces may not vary greatly. It is still noticeable that there is a small mesh reconstruction spike in the landmark corresponding to the anterior ascending process (Fig. S3.3.2.H) but only in the resulting restored specimen mesh. The analyzed landmarks are not affected by this mesh artifact produced during the restoration procedure in order to produce a series of complete specimen reconstructions for easily view the morphological variation. Lack of information in the posterior ascending process (Fig. S3.3.2.G) is more important we can guess that the distal end is anterolaterally oriented. However, lack of posterior ascending process and specimen virtual restoration is the only source of information for the articulation of the fibula. The impact of our own reconstruction of the *Magyarosaurus* hind limb based on the virtual restoration of the left tibia NHM-R3853 is larger than any of the other taphonomic factors as seen in the specimen alone. A slightly more rotated fibula due any other configuration of the posterior ascending process of the distal end of the tibia may produce less variance along PC3 (Fig. S3.1.3) on our analyses and a decreasing the scores along this axis, thus plotting *Magyarosaurus* in less negative PC3 values and also slightly centring the rest of the sample towards zero-score. But given the position of taxa such as *Euhelopus zdanskyi*, this effect on the whole PC3 may be less appreciable in the plotting of the other taxa.

The specimens from Bellevue area usually exhibit some degree of lithostatic compaction that usually translates in anteroposteriorly-narrower proximal and distal ends of the specimens as well as some anteroposterior crushing of the shafts. It is also important to note that there are at least two different morphotypes that may be attributed to two different taxa in the sample of Bellevue area with noticeable differences in the limb morphology (Díez Díaz et al., 2023b; Vila et al., 2012). For this reason we used only the specimens described by Le Loeuff (2005) with the hypothesis that this sample is referable solely to *A. atacis* and none to the second morphotype from Bellevue area (cf. *Lirainosaurus* after Vila et al., 2012). Taxonomic assessment is beyond the scopes of this work and this assumption will not include problematic specimens of the second gracile form. The specimens included in the hind limb of *A. atacis* exhibit the anteroposterior crushing and the proximal and distal end may be slightly anteroposteriorly-narrower due lithostatic compaction. The proximal end of the right femur MDE-C3-87 is extremely anteroposteriorly-narrow but despite some degree of compaction, the proximal end may be lightly-warped as the distal end are well preserved and does not suffer noticeable deformation. Similarly, the distal elements like the tibial proximal and distal ends are well preserved and are not compressed in any direction. The fibula also exhibits the anteromedial deflection of the anterior crest (e.g., MDE-C3-48), allowing us to disregard that some the lateromedially-narrow fibular shafts may also be a product of lithostatic compaction. Despite the fragmentary nature of many of the Bellevue specimens, no taphonomic deformation may have a great impact in our analyses.

Whereas there are several femoral and tibial specimens of *Antarctosaurus wichmannianus*, there is only one left fibula, specimen MACN-6804-21, which exhibits some degree of fracturing and taphonomic deformation of the shaft. The distal end has an oblique anteroposterior fracture and some posterior concavity produced due warping of the distal third of the shaft. This may affect in the plotting of *A. wichmannianus* with a slightly straighter fibula instead and thus closer to other deeply-branched non-colossosaurian non-aeolosaurini titanosaurs (e.g., in PC1; Fig. S3.1.1). The overall results may not alter but if *A. wichmannianus* exhibits less deflection of the fibula, it would remark much more the simplesiomorphic beam-like, less lateromedially-wider morphology of the colossosaurian and aeolosaurini hind limbs compared to other deeply-branched titanosaurians.

In general, we opted for light discussion and conservative conclusions in the evolutionary implications of the fibular rotation in lateral view, as many hind limbs of our study are affected by taphonomy, not directly in the shape of the fibula, but in lack of information on the articulation between the tibia and fibula. It is also important to note that this study lacks the analysis of the astragalus and some of the titanosaurs do not preserve one. The astragalus will dictate the distal articulation between the fibula and tibial pair (see Upchurch et al., 2004). The lack of astragalus impose a bias in how several of the hind limbs with tibia and fibula of subequal length (e.g., *Lirainosaurus astibiae*, most of Colossosauria sampled in this study, among others) articulates despite lack of apparent sigmoidal morphology despite an anterior projection of the distal end. The articulation with the femur tibial and fibular distal condyles coerces the height of the fibular proximal end, at the same height of the tibial proximal end or slightly below. The position and rotation of the tibial ascending processes of the distal end were taken into account. However, the lack of astragalus does not allow us to observe complex articulations beyond the medial fibular articulation with the available elements of the tibia. In the future we would like to expand on the biomechanical implications of the different configuration with the additional evolutionary information of the macronarian astragalus. However, it is beyond the scopes of this study and the sample is lacking to include in our current analyses.

### S3.4. Effects of the Ontogenetic status of the smaller specimens on the body size proxy and the shape analyses

The assessment of the presence of potential juvenile or sub-adult specimens in the sample is important as size differences due differences in ontogenetic status may alter greatly the results of our analyses. In order to control this altering factor, we excluded juvenile and early-juvenile specimens, considering only sub-adult specimens that exhibit some morphological features that indicate that it may have reached its maximum body size.

The relative age of a sauropod individual is usually established based on the standardized Histological Ontogenetical Stages (Klein and Sander, 2008). These stages were initially established through the histological analysis of sauropod long bones (Klein and Sander, 2008; Mitchell et al., 2017; Padian et al., 2001; Sander et al., 2011b) but can be determined using other parts of the skeleton such as the ribs (Klein et al., 2012; Waskow and Mateus, 2017; Waskow and Sander, 2014). The HOS are usually employed to determine life history and growth curves (Klein et al., 2012; Klein and Sander, 2008). Thanks to the advances in paleohistology they have allowed us to observe that sauropods grow extremely fast (Sander et al., 2011a) acquiring its body size earlier the its full maturity in few years (Sander et al., 2011b, 2011a). A “young adult” or probably equivalent to Histological Ontogenetic Stage of 11 or more (Klein and Sander, 2008; Stein et al., 2010) can be safely regarded that has reached its full size. Also, recent studies indicate that titanosaurs exhibit precocial growth and its morphology is acquired and locked at early postnatal ontogenetic stages (Curry Rogers et al., 2016). We opted to exclude smaller specimens, or those which exhibited hints of being subadult specimens (following th criteria established in Griffin and Nesbitt, 2016; Ikejiri, 2004; Ikejiri et al., 2005; applied to titanosaurs as in Páramo et al., 2019) from our sample prior obtaining the theoretical reconstructions of each taxon hind limb nonetheless.

We used the holotype hind limb of *Lohuecotitan pandafilandi* to include it in our database, despite Morphotypi I of Lo Hueco site has been suggested to pertain to the same taxa (Páramo et al., 2022) and none of the smaller, putative juvenile specimens, differ from the larger specimens nor the holotype elements (Páramo et al., 2022). The probable juvenile specimens referred to *Lirainosaurus astibiae* are all teeth or fore limb specimens (Díez Díaz, 2013) and were not considered in the multivariate statistic estimation of the body size.

*Muyelensaurus pecheni* may be problematic to assess which one may pertain to a probable juvenile or subadult specimen without a proper histological sample, as it has been proposed that there are some juvenile individuals in the sample of at least five individuals (Calvo et al., 2007). However, there are no size differences among the appendicular elements. Neither does any of the elements exhibit features that may indicate subadult status or related to earlier ontogenetic stage according to the criteria mentioned before.

There are some smaller specimens among the sample of appendicular elements of *Saltasaurus loricatus* that may pertain to putative juvenile individuals (see Powell, 2003). Specimens like the smaller tibia PVL-4017-87 exhibit not only size differences but less marked scars or rugosities in the proximal and distal ends (APB pers. obs.). However, there are no morphological differences with the larger individuals and previous analyses found this specimens plotted in the same area of the morphospace (e.g. Páramo et al., 2020). Their centroid size were excluded in the calculation of the mean size of each element of the operative taxonomic unit hind limb. Similarly, *Neuquensaurus* spp. includes some smaller specimens that may be referred to subadult individuals. In this case, they are more difficult to assess as some of them has lost the proximal and distal end, where usual criteria of the articular cap rugosity may apply in absence of histological sampling. We included all the sample specimens from previous analyses despite some of them like the tibiae MLP-CS-1123 and MLP-CS-1264. exhibit slightly smaller sized but no morphological differences in hand review nor in morphometric analyses. MLP-CS-1123 may or may not pertain to a juvenile individual as it does not preserve the proximal and distal ends, but in the other hand MLP-CS-1264 exhibit the same rugosity expected as in an adult individual and the larger specimens of the *Neuquensaurus* sample. We opted to include their sizes in the estimation of the operative taxonomic unit hind limb and consider the size differences as intraspecific variability not related to ontogenetic differences. However, we may take into account that *Neuquensaurus australis* and “*N. robustus*” morphological differences and taxonomic status of the latter is still in debate (Otero, 2010).

The sample of classical hind limb specimens referred to *Magyarosaurus* spp. (Nopcsa, 1915) exhibit some degree of size differences but none of the sampled specimens may be referred to a juvenile individual neither do their size differs greatly. Similarly to *Neuquensaurus* spp., taxonomic status of “*Magyarosaurus hungaricus*” and “*M. transylvanicus*” is still debated (e.g. Díez Díaz et al., 2023a; Mocho et al., 2024; Upchurch et al., 2004). Probable unknown interspecific variability that here is considered intraspecific variability could have greater impact than ontogenetic differences in our analyses. However, a preliminary study plotted all the *Magyarosaurus* specimens from Nopcsa classic collection in the same area of the morphospace, closer between them (Páramo, 2020).

Lastly, there are smaller specimens identified in the sample of Ampelosaurus atacis. In this case there is a lack of histological sample in all the sampled specimens, but we are more worried about the mixture of some specimens that may be referred to a second taxonomic unit. For now there is a proposal of the presence of a second morphotype, referable to another different undrescribed sauropod, cf. *Lirainosaurus* (Vila et al., 2012). In situ observations indicate there are large morphological differences as well as size differences between both morphotypes (Díez Díaz et al., 2023b; Vila et al., 2012) and our preliminary study found similar results in the morphometric analyses (Páramo, 2020). We feel that the interspecific variability due an undescribed titanosaur mixed in the sample of *A. atacis* may have a larger impact in our analyses than the ontogenetic status of some of the appendicular specimens itself. For this reason and until taxonomic status of the sample of *A. atacis* is assessed (e.g. Díez Díaz et al., 2023b) we used the referred specimen of Le Loeuff (2005) as the operative taxonomic unit hind limb of *A. atacis*. All the elements exhibit enough features to consider them at least subadult specimens.
